# Restoration of Cone Circuit Functionality in the Regenerating Adult Zebrafish Retina

**DOI:** 10.1101/2023.11.11.566441

**Authors:** Evelyn Abraham, Hella Hartmann, Takeshi Yoshimatsu, Tom Baden, Michael Brand

## Abstract

Unlike humans, teleosts like zebrafish exhibit robust retinal regeneration after injury from endogenous stem cells. However, understanding the functional recovery of the regenerated retina remains a challenge. In particular, it is unclear if regenerating cone-photoreceptors can regain physiological function and integrate correctly into post-synaptic circuits. To bridge this gap, we employed two-photon calcium imaging of living retina, focusing on photoreceptor responses before and after an intense light-induced lesion in the adult zebrafish retina. To assess the functional recovery of cones and their downstream circuits, we exploited the colour opponency in adult zebrafish cones. We find that UV cones exhibit an intrinsic Off-response to short-wavelength blue light, but an On-response to longer-wavelength green light, which depends on feedback signals from outer retinal circuits. Accordingly, we examined the presence and quality of Off-versus On-responses, to assess the functional recovery of cones and their correct integration into outer retinal circuits. We find that regenerated UV cones regain both Off-responses to short-wavelength and On-responses to long-wavelength within 3 months after light lesion. Hence, physiological circuit functionality is restored in regenerated cone photoreceptors, suggesting that inducing endogenous regeneration is a promising strategy for human retinal repair.

**Graphical abstract:** 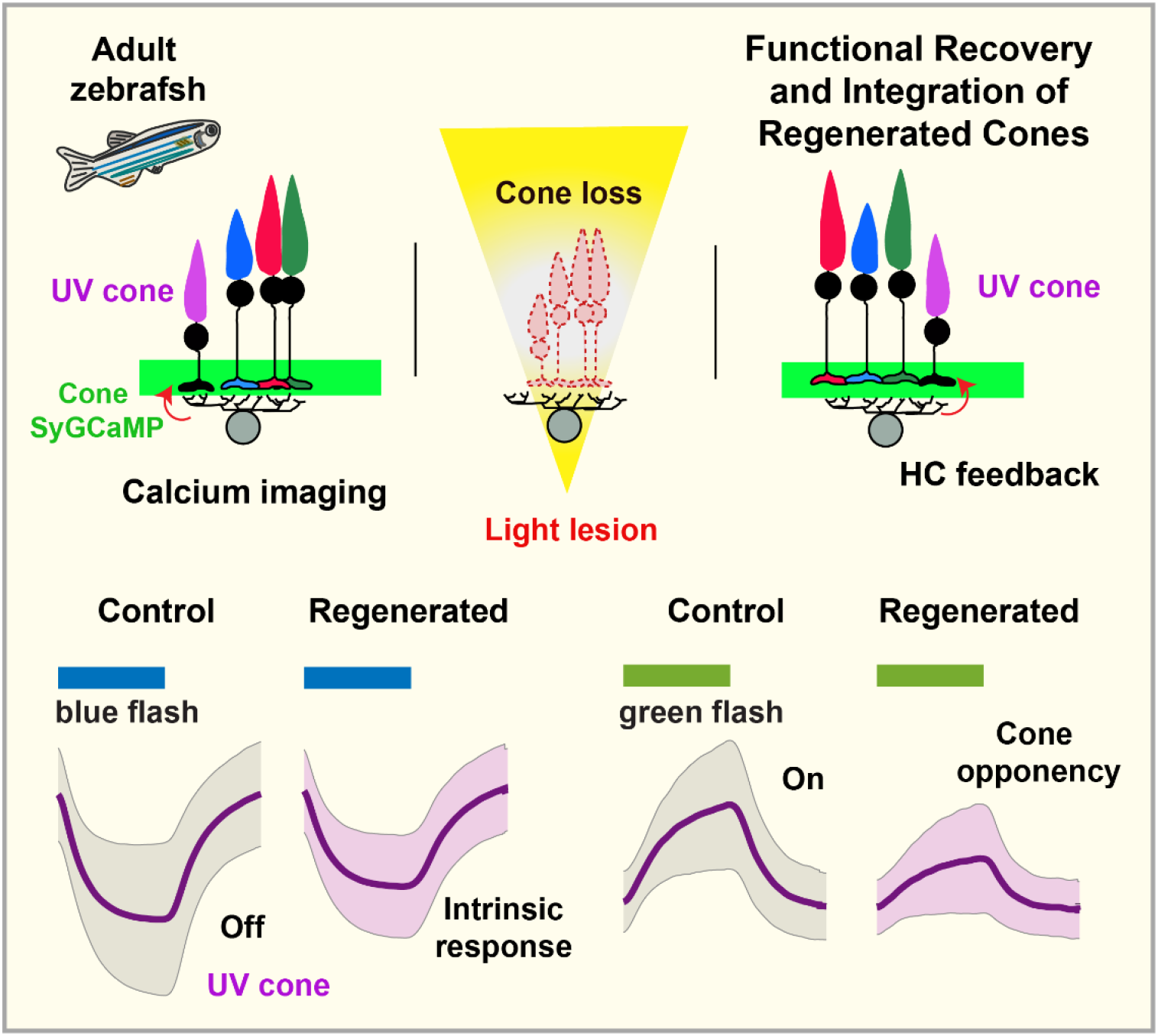

## Introduction

Visual impairment or blindness due to neuronal degeneration is a severe problem affecting humans (Karl & Reh, 2010). In contrast to mammals, zebrafish (*Danio rerio*) harbour a robust intrinsic capacity to regenerate photoreceptors and other retinal neurons shortly after injury and serve as an excellent model with potential translational impact (Otteson & Hitchcock, 2003; Wan & Goldman, 2016). Upon retinal damage, the radial glia in the adult zebrafish retina, Müller glia cells (MG), re-enter the cell cycle and function as the primary stem cell source (Bernardos et al., 2007). De-differentiated MG produces multipotent neuronal progenitors, which subsequently proliferate and differentiate to regenerate lost retinal cells, leading to a remarkable restoration of the retina’s cellular and synaptic layers (Lenkowski & Raymond, 2014; Sherpa et al., 2008). Although the regeneration of retinal morphology is evident through histological observations, achieving functional recovery remains a critical challenge, especially for newborn retinal neurons and the central nervous system (Fleisch et al., 2011).

Therefore, we asked two fundamental questions:

i. Do regenerated neurons themselves recover normal light-evoked physiology?
ii. Can regenerated neurons integrate functionally into post-synaptic circuits?

To this end, we developed a neural activity readout from cone photoreceptor terminals that enables to study light-evoked activity as well as synaptic feedback from post-synaptic circuits in the adult zebrafish retina and combined it with a reliable and non-invasive cell ablation paradigm to induce retinal regeneration. *Intense light lesion* is a sterile and effective photoreceptor-specific ablation method that enables to study newly regenerating photoreceptors (Qin et al., 2009). Intense light lesion with diffuse UV light has been characterized with respect to photoreceptor subtype specificity, cell death, retinal topography, histology, and optical coherence tomography in normally pigmented adult zebrafish (Weber et al., 2013). Shortly after lesion, UV cones are maximally ablated, followed by blue, green, and red cones in a wavelength-dependent manner across the retinal surface, with the central retina entirely ablated of all photoreceptors. Histological recovery of photoreceptor regeneration is complete within 28 days post-lesion (dpl) (Weber et al., 2013). Hence, diffuse UV light lesion is an effective method for examining the loss of photoreceptor function and its recovery. We thus sought to understand whether *UV cones* and the surrounding *non-UV* (blue, green, and red) cones regenerate normal physiology and light responses.

Here, we established a customized two-photon setup for adult zebrafish retinal explants and combined it with calcium imaging of cone photoreceptors and light stimulation. We found that the chromatic responses of regenerating UV cones showed robust functional recovery with response quality, kinetics and functional connectivity comparable to the unlesioned retina, gradually over 3 months post-lesion. Importantly, both before ablation and after regeneration, cones exhibited light-evoked Off- and On-signals. Since cones are intrinsically Off-cells, the presence of On-signals reflects feedback from the horizontal cell outer retinal network, substantiated through pharmacological validation. This suggests that functional recovery occurs not only of cones themselves, but also reflects their circuit integration into the outer retinal network. Our findings show that physiological function is restored in cone photoreceptors regenerating from endogenous MG stem cells and indicate for the first time that complex visual responses/pathways, such as synaptic processing in cones and colour circuits, are restored, culminating in functional regeneration of adult retina.

## Results

### Adult zebrafish UV cones are strongly colour-opponent

We initiated our study by establishing *ex vivo* adult zebrafish retinal whole-mount preparations (retinal explants) for live two-photon (2P) imaging using an inverse 2P microscope (**Fig. 1A, S1A-C**). To conduct calcium imaging in all cone photoreceptor terminals across the outer plexiform layer, we used a transgenic reporter carrying the transgene *gnat2:SyGCaMP6f*, in which the guanine nucleotide-binding protein, alpha transducing activity polypeptide 2 (gnat2)-promoter, drives GCaMP6f tethered to synaptic vesicles (SyGCaMP6f) (Dreosti et al., 2010; Kennedy et al., 2007) (**Fig. 1A-B**). For a comprehensive analysis, we simultaneously visualized cone SyGCaMP6f along with a UV cone marker line (Yoshimatsu et al., 2016) in the double transgenic background *Tg(gnat2:SyGCaMP6f); Tg(opn1sw1:nfsb-mCherry)*. Notably, adult zebrafish cones display a highly precise geometric mosaic pattern (Robinson et al., 1993); the arrangement of cone pedicles, as depicted in **Fig. 1B**, with single rows of alternating short-wavelength UV and Blue (B) cones, accompanied by double rows comprising pairs of Red (R) and Green (G) cones, with R cones flanking UV cones (Li et al., 2012), was evident in this preparation.

**Figure 1:**
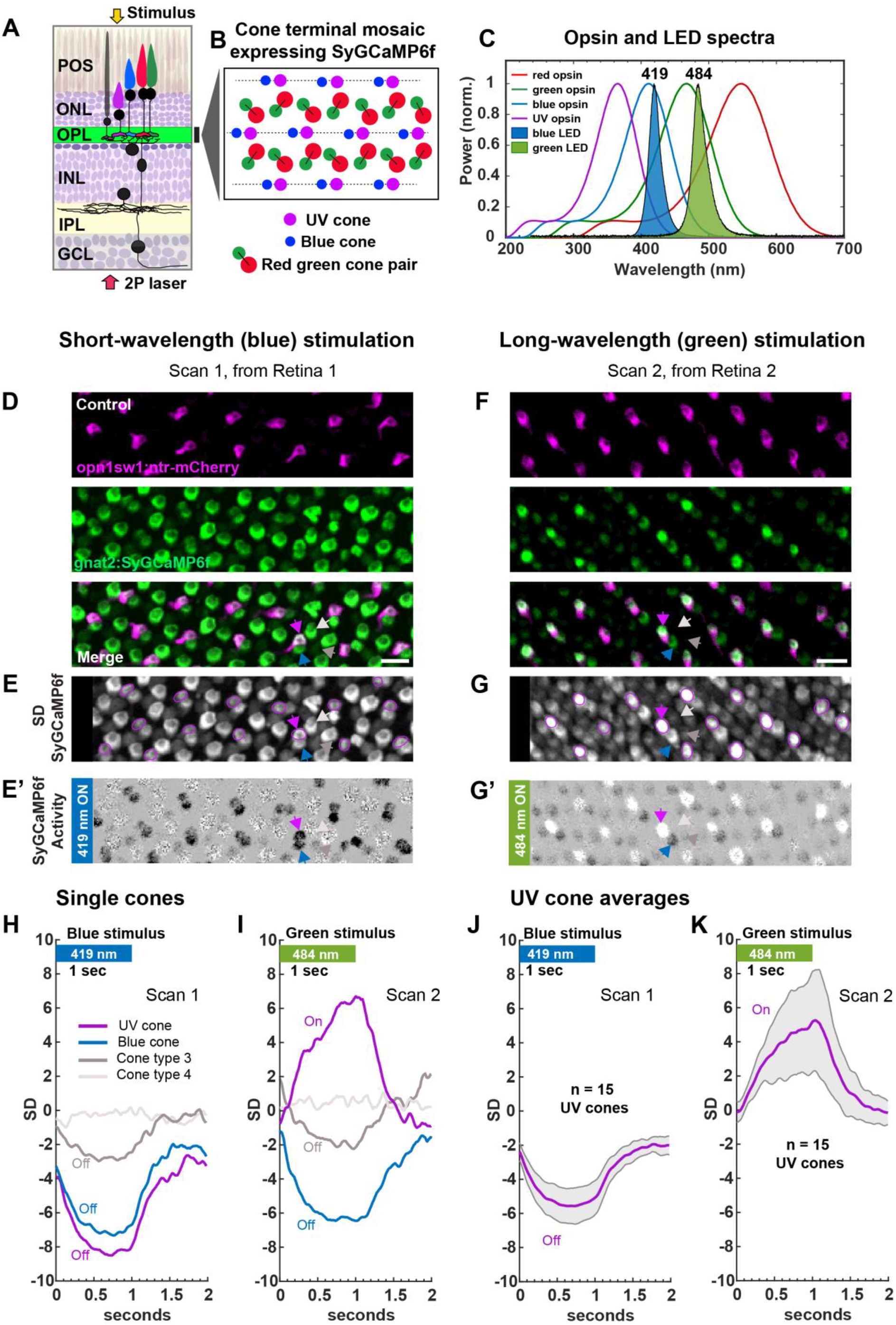
Adult zebrafish cone responses to short wavelength (blue, 419 nm) and long wavelength (green, 484 nm) light stimulation. (**A)** Schematic of adult zebrafish retina expressing SyGCaMP6f in all cone terminals in the outer plexiform layer (OPL). 2P imaging laser is delivered from the retinal ganglion cell side (red arrow), and light stimulation from the photoreceptor side (yellow arrow); see also **Fig S1A-C**. **(B)** Schematic of cone terminal mosaic of adult zebrafish (adapted from (Li et al., 2012)). POS: photoreceptor outer segments, ONL: outer nuclear layer, OPL: outer plexiform layer, INL: inner nuclear layer, IPL: inner plexiform layer, GCL: ganglion cell layer. **(C)** Blue LED, green LED spectrum peaking at 419 nm and 484 nm, overlayed with zebrafish cone opsin absorption spectrum. **(D and F)** Example functional scan fields from the central retina of double-transgenic *Tg(gnat2:SyGCaMP6f); Tg(opn1sw1:nfsb-mCherry)* retinae, stimulated with blue and green light, respectively. D and F are separate scans from different retinas. Green-SyGCaMP6f, magenta-UV cone mCherry; Scale bar = 10 micrometre. **(E and G)** Standard deviation (SD) projection of SyGCaMP6f from D and F with UV cone mosaic region of interest (ROI) marked in magenta circles. **(E’ and G’)** Activity images (see methods) of D and F, showing synaptic cone SyGCaMP6f calcium responses to blue and green stimulus respectively. Dark terminals represent Off responses, whereas lighter terminals represent On responses. Here numerous Off responses among UV and blue cones (dark hues), and weaker Off responses from unidentified cones in-between are seen to blue light. In contrast, UV cones elicit clear On responses to green light, while the remaining cones maintain Off responses. Quartet of cone terminals with UV cone (magenta arrow), blue cone (blue arrow), cone type 3 (dark grey arrow) and cone type 4 (light grey arrow) are indicated. Cone type 3 and 4 are presumably red or green cones however are unidentified. **(H and I)** Single cone light responses from quartet of cones marked with coloured arrows in E’ and G’ (also D-E, and F-G), respectively. **(J and K)** Light responses of n = 15 UV cones from E and G, respectively, represented as mean and standard deviation (shaded error bar). H-K: X-axis: time (seconds); Y-axis: standard deviation (SD). Blue bar: blue (419 nm) stimulus for 1 sec; Green bar: green (484 nm) stimulus for 1 sec; see also **Fig. S1A-C, Movies: 1-2** corresponding to E’ and G’.

Full-field photopic light stimulation was delivered to the flat-mounted retinae from above using a *short-wavelength blue* (419 nm) and a *long-wavelength green* (484 nm) LED targeting the central range of cone opsin spectra, allowing the activation of all cone subtypes (**Fig. 1A-C, S1A-C)**. Example scans from the central retina in response to blue and green light stimulation are shown in **Fig. 1D, 1F**. The distinct UV cone mosaic allows to morphologically identify its neighbouring blue cones, however, additional genetic labelling is required to sort by type, red and green cones occurring between UV and blue cones (**Fig. 1D-E, 1F-G**). The light responses of a group of four cone terminals that included UV, blue, and unidentified red/green cones to blue and green light are shown in **Fig. 1D-I**. Following the presentation of blue light, distinct arrays of cone terminal mosaics elicit Off responses, corresponding to terminals with darker-than background hues in the light-response (activity) image (**Fig. 1E’, Movie. 1)**. Conversely, the presentation of green light evokes both Off and On responses across arrays of cones, corresponding to terminals with darker and lighter hues, respectively (**Fig. 1G’, Movie. 2**). Notably, UV cones exhibit robust Off-responses to blue light, yet their responses inverted to On-responses under green light **Fig. 1E, 1E’, 1G, 1G’, 1H-K**. In contrast, blue cones elicit Off responses to both blue and green light, while the remaining cones show either Off or weakly On responses or were unresponsive under both conditions (see example traces in **Fig. 1H-K**). While intrinsic Off-responses at the cone pedicle stem from phototransduction, On-responses reflect feedback originating from the outer retinal circuits, as previously observed in the larval retina (Yoshimatsu et al., 2021). Therefore, the simultaneous occurrence of Off and On antiphase cone responses to green light in the adult zebrafish retina indicates not only functional cones, but also that these cones are connected within functional colour circuits (compare **Movies: 1-2**). Here, we observe that among the four cone types, UV cones invert their responses from Off to On in response to a shift from short to long wavelength stimulation. Our findings highlight substantial colour opponency in adult zebrafish UV cones, rendering them a valuable model for investigating functional restoration and circuitry integration of retinal neurons.

### Temporal Progression in Restoring Cone Function After Diffuse Light Lesion: Off Responses at 8 dpl to Complete Recovery by 8 mpl

To investigate the loss of cone function and its ‘recovery’ during regeneration, we used diffuse light lesion with high-intensity UV light (>200,000 lux) that induces a large centrally located lesion where all photoreceptors are ablated (**Fig. 2A-B**). Cell death of photoreceptors in the central retina peaks at 24 hours post-lesion, and by 3 days post-lesion, cone debris is cleared; photoreceptors are morphologically regenerated within 28 days post-lesion (Weber et al., 2013). We continuously monitored for regenerating cones in the central retina by recording light-evoked activity from light-lesioned retinae at short-term and long-term time points that included: Control (unlesioned), 4 dpl, 8-9 dpl, 10-11 dpl, 14-15 dpl, 28-29 dpl, 3 mpl, and 8 mpl (**Fig. 2C**) (dpl: days post-lesion, mpl: months post-lesion). Representative scans showing the cellular-level effects of lesion and regeneration in the central retina during the acquired time course are depicted in **Fig. S2**. After the lesion, at 4 dpl time point, the central retina is devoid of mCherry-positive and SyGCaMP6f-positive cell populations, which instead is sparsely filled with any remaining debris only (**Fig. S2**). By 8-9 dpl, the UV cone mCherry expression recovers, along with cone SyGCaMP6f, signifying the emergence of newly regenerating cones, which are present at all later time points until 8 mpl (**Fig. S2**). Our data suggests that differentiation into cone photoreceptors post-light lesion begins as early as 8 dpl after cell death and Müller glial cell activation and continues up to 8 mpl (see below).

**Figure 2:**
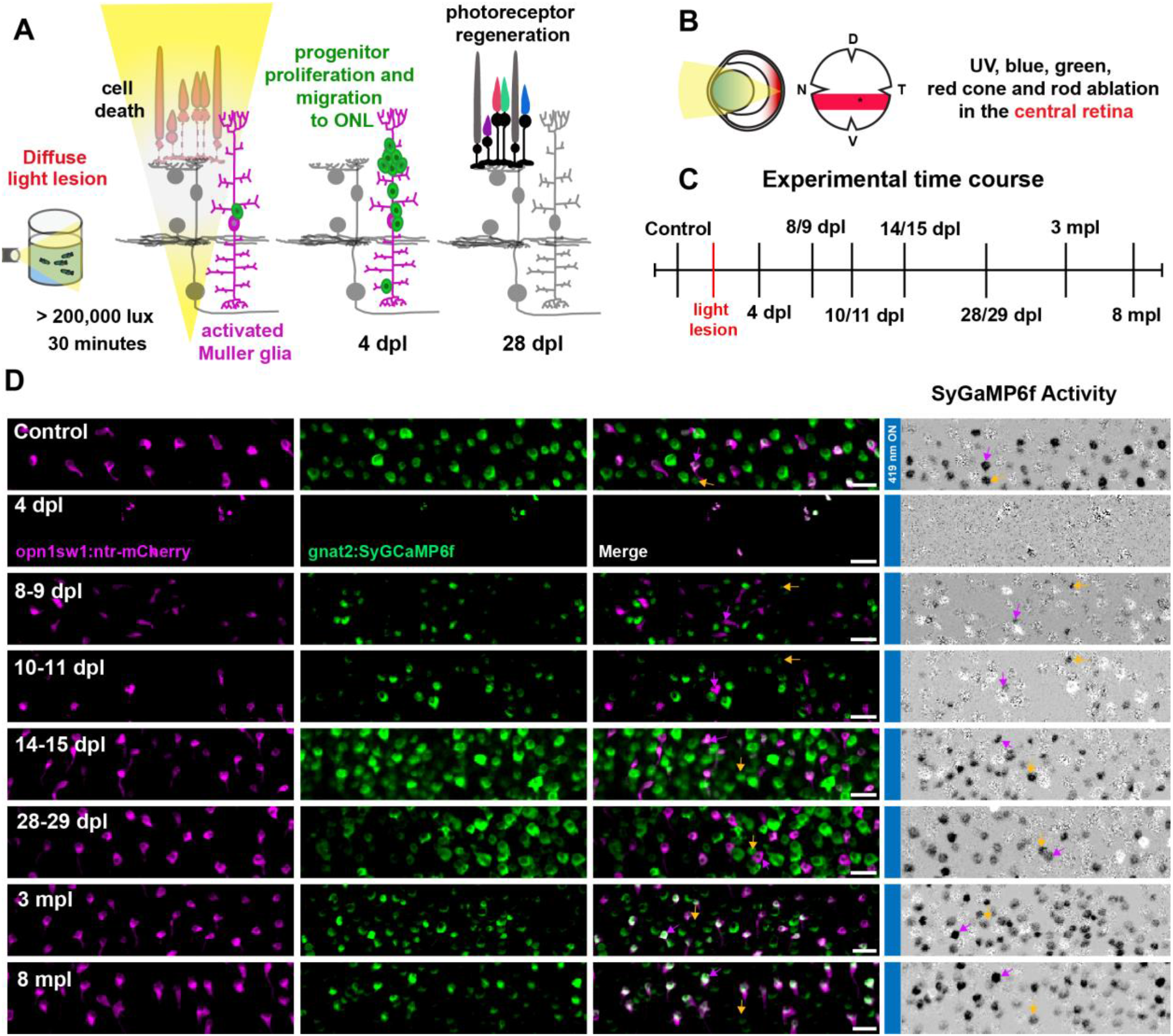
Recovery of Off responses in the regenerating retina after intense light lesion. **(A and B)** Schematic representation of diffuse light lesion induced photoreceptor ablation and regeneration in the adult zebrafish retina on retinal sections and flat mounts (Weber et al., 2013). Retinal regeneration following photoreceptor ablation is initiated by MG activation and dedifferentiation to form progenitors. Proliferating progenitors undergo interkinetic nuclear migration and differentiate to form regenerated rod and cone photoreceptors within 28 dpl. In wholemounts, diffuse light lesion leads to a large centrally located lesion (central stripe) shown in red, where all four cone photoreceptors subtypes and rods are ablated. UV and blue cones are also ablated in the dorsal retina, while red/green cones and rods are not. In contrast, the ventral retina photoreceptors are morphologically spared of the lesion. **(C)** Experimental time course of functional studies encompassing 8 time points: Control, 4 dpl, 8-9 dpl, 10-11 dpl, 14-15 dpl, 28-29 dpl, 3 mpl, and 8 mpl. **(D)** Representative functional scans and activity images from the central retina of double transgenic *Tg(gnat2:SyGCaMP6f); Tg(opn1sw1:nfsb-mCherry)*, during regeneration in response to blue light stimulation. Fourth column indicates SyGCaMP6f activity at the indicated time points of regeneration. Dark terminals represent Off responses, whereas lighter terminals represent On responses. Example UV cone Off responses are indicated with magenta arrow and non UV cone Off responses with yellow arrow, which can be seen at all time points except 4 dpl. Representative scans shown here are selected from 80, 20, 17, 32, 41, 99, 28 and 48 scans, respectively (N = 10, 3, 3, 6, 7, 6, 3 and 3 retinae), from the indicated time points: control, 4 dpl, 8-9 dpl, 10-11 dpl, 14-15 dpl, 28-29 dpl, 3 mpl and 8 mpl. In all experimental groups, only ‘responsive’ scans are included, where at least one cone responded to the stimulus. In total, the scans yielded n = 854, 250, 457, 744, 1281, 600 and 886 UV cones from control and the six regenerating time points 8-9 dpl to 8 mpl; of these, the proportion of responding UV cones steadily increased from 8-9 dpl until 8 mpl (**Fig. S3I**); see also **Movies: 3-10** corresponding to D, and **Fig. S1-S5.** Green-SyGCaMP6f, magenta-UV cone mCherry. Scale bar = 10 micrometre. ONL: outer nuclear layer, MG: Müller glia, dpl: days post-lesion, mpl: months post-lesion.

To assess the functionality of regenerating cones in the central retina, we presented blue light stimuli and specifically monitored UV cones; representative functional scans and corresponding activity images are shown in **Fig. 2D**; see also **Movies: 3-10**. No light responses were recordable within the lesioned central regions at 4 dpl, which exhibited distinct noise patterns, indicating loss of function. Newly emerging UV cones at 8-9 dpl, and 10-11 dpl elicited very weak Off-responses. However, the UV cones regain clearer Off-responses from 14-15 dpl until 28-29 dpl, which then become highly robust from 3 mpl until 8 mpl, resembling controls; **Fig. 2D, Movies: 3-10**. To further understand UV cone functionality at the population level, we evaluated their response traces during regeneration. UV cone light responses were evident in the control and, importantly, at all six regenerating time points starting from 8-9 dpl at both the single cone and population level (**Fig. S1D-E, S3**). As expected, ROI traces at 4 dpl were absent due to cell ablation. Furthermore, the proportion of “Off” responses within the regenerated light-responsive UV cones consistently remained >95%, indicating a high level of functional precision (**Fig. S3I-K**), despite a disrupted arrangement of the cone mosaic pattern of regenerated cone terminals (**Fig. 2D, S1D, S2**). Importantly, mCherry-negative, SyGCaMP6f-positive surrounding non-UV cones are also regaining their short-wavelength light responsivity (Off responses) during regeneration, functionally reflecting the normal responses of blue, green or red cones within 3 mpl (**Fig. 2D, Movies: 3-10**; see also **Fig. S8A-G, S8O-P)**. Further, the ventral retina remained unlesioned with intact cones (in accordance with (Weber et al., 2013; Raymond et al., 2014)), and the ventral UV cones continue to elicit light responses during lesion and regeneration, offering an excellent internal control (**Fig. S4, S5**). Consequently, the disrupted UV cone mosaic served as a primary landmark to identify regenerating cones within the lesioned central retina. Overall, the light-induced responses of regenerating UV cones demonstrate the gradual recovery of cone functionality over time.

### Progressive Functional Recovery culminates in Reliable, Sensitive, and Rapid UV Cone Off Responses

To examine regenerating UV cone physiology more quantitatively, we examined the Off Ca^++^-waveforms triggered by short-wavelength blue light over the time course of regeneration. We used a single exponential function to assign values for ‘amplitude’, ‘decay’ and ‘recovery’ time constants (Tau) from the measured UV cone Off response waveforms (methods). The amplitude measurement indicates response strength, while Tau signifies the speed of response during both decay and recovery phases. At 8-9 dpl, the waveform was highly diminished compared to that of controls, which improved only slightly until 14-15 dpl (**Fig. 3A**). The UV cone response waveform progressively improved over time, but still remains smaller than controls at 28-29 dpl. Our amplitude analysis revealed a substantial reduction in the Off-response amplitude at 8-9 days post-lesion (dpl) compared to unlesioned controls. This reduction persisted, with a significant effect size, until 28 dpl, and until 3 mpl with a moderate effect size, until near-complete restoration is reached by 8 mpl (81% of control values; **Fig. 3B, Table. 1**). Therefore, newly regenerated UV cones exhibit reduced sensitivity to light stimuli that is then regained over a period of 3-8 months of functional regeneration. Similarly, we quantified the decay and recovery speeds (kinetics) of UV cone Off responses using Tau decay and Tau recovery values (methods). Interestingly, Tau decay remained unchanged compared to controls, whereas Tau recovery exhibited a high value in the early stages of regeneration that eventually fully recovered by 8 mpl **(Fig. 3C-D, Table. 2-3)**. Thus, UV cones regenerated in the initial stages appear to be ‘immature’ with respect to their calcium (Ca++) recovery kinetics. Finally, at 3-8 mpl, the waveforms resemble control waveforms in size and shape (**Fig. 3A, 3B-D**).

**Figure 3:**
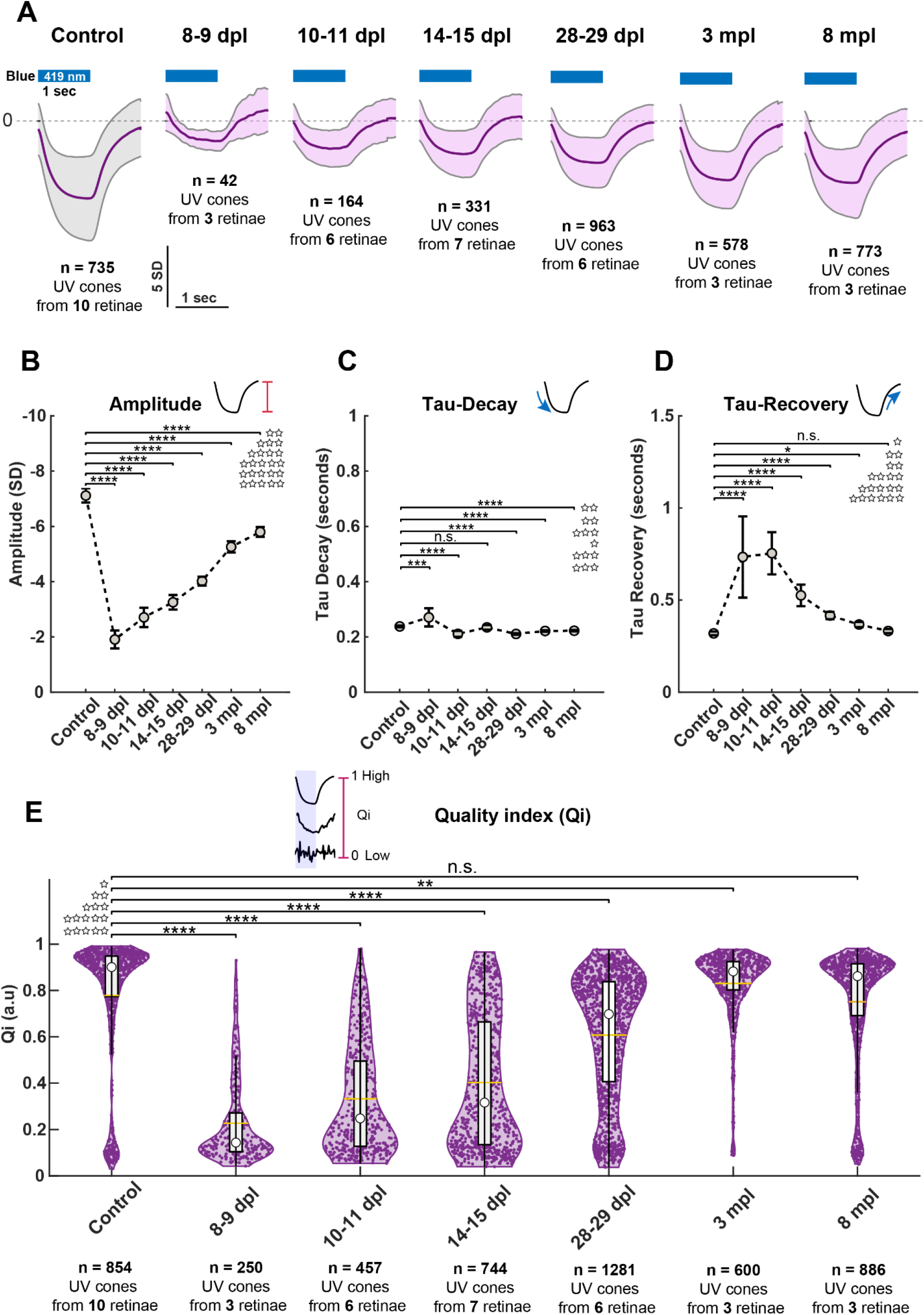
Quantitative evaluation of progressive functional recovery of UV cone Off responses. **(A)** Average response of all light responsive UV cones (Qi >= 0.4) during regeneration time course represented as mean and standard deviation (shaded error bar). Sample size (n): Control-735, 8-9 dpl-42, 10-11 dpl-164, 14-15 dpl-331, 28-29 dpl-963, 3 mpl-578, 8 mpl-773 (n = UV cones). Blue bars are stimulus presentations with blue (419 nm) light for 1 second; X-axis: time (seconds); Y-axis: standard deviation (SD). **(B-D)** Development of UV cone Off response amplitude, Tau decay and Tau recovery during regeneration | **(B)** Amplitude: The recovered amplitudes at 8-9 dpl, 14-15 dpl, 28-29 dpl, 3 mpl, and 8 mpl corresponded to 27%, 38%, 46%, 56%, 74%, and 81% of control levels. **(C)** Tau decay **(D)** Tau-recovery. X-axis: time points: Control – 8 mpl; Y-axis: Amplitude (SD) or Tau (seconds). B-D: Data represented as represented as mean and 95% confidence interval (95% CI); Sample size (n) for B-D: Control-576, 8-9 dpl-28, 10-11 dpl-89, 14-15 dpl-182, 28-29 dpl-643, 3 mpl-469, 8 mpl-571 (n = UV cone Off responses). **(E)** Violin plots of the development of UV cone response quality index (Qi) during regeneration. Each dot is one cell. Each violin plot is overlayed with boxplot. Yellow horizontal line shows the mean. Sample size (n): Control-854, 8-9 dpl-250, 10-11 dpl-457, 14-15 dpl-744, 28-29 dpl-1281, 3 mpl-600, 8 mpl-886 (n = UV cones); X-axis: time points: Control – 8 mpl; Y-axis: Quality index (Qi) in arbitrary units (a.u.). A-E from N= 10, 3, 6, 7, 6, 3, and 3 retinae at respective time points (Control - 8 mpl). Statistical test: One sample ANOVA, followed by Tukey-Kramer post-hoc multiple comparisons test at alpha level 0.05. **** p < 0.0001, *** p < 0.001, ** p < 0.01, * p < 0.05; n.s.-not significant; For all pairwise comparisons, we report the effect sizes as Cohen’s d (d): d > 2.0 huge, d > 1.20 very large, d > 0.8 large, d > 0.5 medium, d > 0.2 small, d > 0.01 very small. Group means, confidence intervals, summary statistics and effect sizes for B-E in **Table 1-4.** A-E: extended analysis of **Fig. S3**; see also **Movies: 3-10.**

In order to quantify the reliability of UV cone responses to light, we calculated a quality index (Qi) reflecting the response likelihood across 10 serial stimulus trials (methods). Qi tends to ‘1’ for high quality and reliable responses, and ‘Qi>=0.4’ was used as a threshold to define ‘light responsive’ cones. The resulting Qi violin plots (**Fig. 3E, Table. 4**) revealed that newly regenerating UV cones at early time points (8-9 dpl) respond poorly, with many low-quality responses (Qi: 0-0.4), reflecting random noise or very poor Off responses after blue light stimulation. Subsequently, the Qi values improve until very high-quality light responses with Qis of 0.8-1 are reached from 3 to 8 mpl that are indistinguishable from control Qis. Thus, our Qi analysis reveals that the quality of the UV cone response progressively recovers over time. Taken together, the quantification of response amplitude and kinetics of the regenerating UV cones suggest that the newborn UV cones are initially significantly impaired in sensitivity, and noisy and slow to recover normal kinetics of their Off responses. Remarkably, over the time course of regeneration, Off responses become stronger, highly reliable and fast, culminating in complete functional restoration by 8 mpl.

### Renewing Colour Circuitry: Restoration of UV Cones and Reinstatement of Horizontal Cell Feedback

Our findings above show that intrinsic physiological properties of UV cones regenerate; next, we asked whether they also re-establish their embedding into outer retinal circuitry. Specifically, we tested whether the regenerated UV cones regain their ‘opponent’ response characteristics (**see Fig. 1**) when exposed to long-wavelength green light. We find that regenerated UV cones invert their responses from Off to On in response to a shift from short to long wavelength stimulation. Regenerated UV cones recovered their On responses, with the proportion of “On” responses within the 2.5 mpl regenerated light-responsive UV cones at 78% (**Fig. 4A, S6, Movie. 11**). The response amplitude and quality were reduced (**Fig. 4B-C, Table: 5-6**), probably due to still ongoing maturation. Because cone opponency is mediated by horizontal cell (HC) feedback, we pharmacologically inhibited HCs using a competitive antagonist drug, CNQX, which blocks AMPA and Kainate receptors (Yoshimatsu et al., 2021). Green light stimulation was performed before and after CNQX treatment at identical scans in control, and 2.5 mpl retinae, and both UV and surrounding non-UV cone responses were analysed. Example activity scans and single cone responses are shown in **Fig. 4D, 4E-E’, 4F, 4G-G’** (see also **Movies: 12-13** and population activity in **Fig. S7**). Strikingly, horizontal cell blocking by CNQX effectively abolished all On responses in both control and 2.5 mpl regenerated retinae, while Off responses in surrounding non-UV cones remained unaffected (see activity scans: **Fig. 4D, 4F** and traces in **Fig. 4E-E’, 4G-G’; Movies: 12-13**). In conclusion, the recovery of ‘On’ responses provides functional evidence for the appropriate integration of regenerating UV cones within the horizontal cell-outer retinal network and proves that cones are connected within colour circuits. Furthermore, the collective responses observed in regenerated non-UV cones indicate the restoration of normal light responses of blue, green or red cones to long-wavelength as well (**Fig. S8H-N, S8Q-R**). Despite the disrupted cone mosaic, non-UV cones responses closely resemble those of the control group, providing robust evidence in support of the successful re-establishment of outer retinal colour circuits (**Fig. S8**).

**Figure 4:**
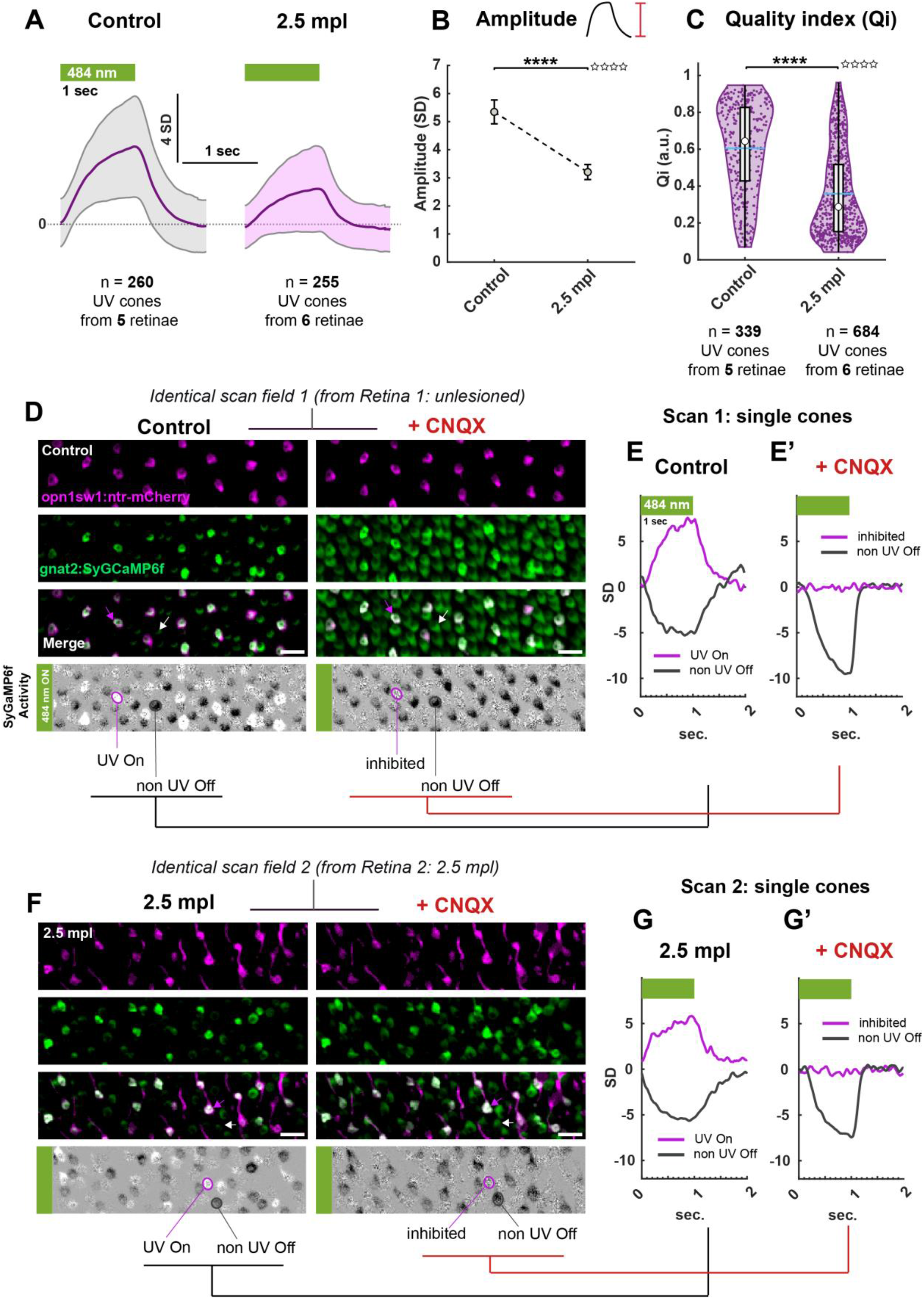
Functional recovery of UV cone On-responses and horizontal cell circuits in the regenerated retina. (**A)** Average response of all light responsive UV cones (Qi >= 0.4) in control and 2.5 mpl regenerated retinae, represented as mean and standard deviation (shaded error bar). Sample size (n): Control-260, 2.5 mpl-255; (n = UV cones) in respective time points. These light-responsive UV cones are from 32 and 44 scans; n = 339, and 684 total UV cones respectively. In all experimental groups, only ‘responsive’ scans are included, where at least one cone responded to the stimulus; X-axis: time (seconds); Y-axis: standard deviation (SD). **(B)** Amplitude of UV cone On responses represented as mean and 95% confidence interval (95% CI). Sample size (n): Control-182, 2.5 mpl-138, (n = UV cone On responses) **(C)** Violin plots of the development of UV cone response quality index (Qi) during regeneration. Each dot is one cell. Each violin plot is overlayed with boxplot. Blue horizontal line shows the mean. Sample size (n): Control-339, 2.5 mpl-684 (n = UV cones); X-axis: time points: Control and 2.5 mpl; Y-axis: Quality index (Qi) in arbitrary units (a.u.). A-C: Data from N= 5 and 6 retinae at respective time points; extended analysis shown in **Fig. S6**; see also **Movie: 11. (D)** Representative unique functional scan and its activity from the central retina of double transgenic *Tg(gnat2:SyGCaMP6f); Tg(opn1sw1:nfsb-mCherry)*, in control retina before and after CNQX addition. Example UV On and surrounding non-UV Off cone before and after CNQX are represented in activity images in magenta and grey respectively (or magenta and white arrows). Corresponding single cone responses to green (484 nm) stimulus is shown in **(E and E’)**. **(F)** Representative unique functional scan and its activity from the central retina of double transgenic *Tg(gnat2:SyGCaMP6f); Tg(opn1sw1:nfsb-mCherry)*, in 2.5 mpl regenerated retina before and after CNQX addition. Example UV On and surrounding non-UV Off cone before and after CNQX are represented in activity images in magenta and grey respectively (or magenta and white arrows). Corresponding cone responses to green (484 nm) stimulus is shown in **(G and G’)**. Activity images: Dark terminals represent Off responses, whereas lighter terminals represent On responses. HC block successfully abolished all On responses before and after regeneration, while Off responses remain. Data from N = 4, and 3 retinae in control and 2.5 mpl respectively; D-G’: extended analysis shown in **Fig. S7**; see also **Movie: 12-13** corresponding to D and F; E, E’, G, G’: X-axis: time (seconds); Y-axis: standard deviation (SD). Green bars are stimulus presentations with green (484 nm) light for 1 second. Green-SyGCaMP6f, magenta-UV cone mCherry, Scale bar = 10 micrometre. Statistical test: Two-sample t-test at alpha level 0.05; **** p < 0.0001, *** p < 0.001, ** p < 0.01, * p < 0.05; n.s.-not significant; we report the effect sizes as Cohen’s d (d): d > 2.0 huge, d > 1.20 very large, d > 0.8 large, d > 0.5 medium, d > 0.2 small, d > 0.01 very small. Group means, confidence intervals, summary statistics and effect sizes for B-C in **Table: 5-6.**

## Discussion

While existing studies indicate remarkable histological recovery of regenerated neurons, the functionality and functional circuit reintegration of regenerating photoreceptors has remained elusive. In this work, we analyse quantitatively if endogenous regeneration in the adult zebrafish retina achieves normal physiological function of photoreceptors, as the cell type that is most frequently affected in human blinding diseases. Notably, our results signify for the first time that regenerating UV cones exhibit excellent functional recovery of intrinsic Off responses and functionally integrate within the existing post-synaptic horizontal cell circuitry, restoring also their cone-opponent On responses in a wavelength-dependent manner; reinstating cone circuits in the regenerated retina. These remarkable insights emphasize the degree to which zebrafish endogenous retinal regeneration can be successful.

### Zebrafish Photoreceptor Regeneration: Insights and Challenges

The processes of photoreceptor structural breakdown, regional loss, and morphological reconstruction are well characterized in the context of light lesions in adult zebrafish (Thomas et al., 2012; Weber et al., 2013). Key cellular stages include: 1 dpl: peak cell death, 3 dpl: MG progenitor proliferation, 5 dpl: MG progenitor differentiation initiation, 7 dpl: first regenerated photoreceptors, 10 dpl: photoreceptor differentiation, 14 dpl: ongoing photoreceptor growth, and 28 dpl: complete anatomical regeneration (Weber et al., 2013; Thummel et al., 2008; Kramer et al., 2021; Kassen et al., 2007; Lenkowski & Raymond, 2014; Ranski et al., 2018; Thomas et al., 2012, 2016; Hammer et al., 2022; Celotto et al., 2023; Vihtelic & Hyde, 2000). Our cone regeneration study aligns with this scheme and explores further time points, both *short-term* and *long-term*. We detect cone loss at 4 dpl followed by the appearance of regenerating cones by *8 dpl* that persist and mature through subsequent regeneration stages, up to *3-8 mpl*. While there are indications of developmental fate specification programs being recapitulated in regeneration (Lahne et al., 2021; Hoang et al., 2020; Celotto et al., 2023), histological abnormalities such as excess neurons, differences in retinal lamination and neurogenic clusters are frequently reported in different lesion models (Barrett et al., 2022; McGinn et al., 2018, 2019; Powell et al., 2016; Sherpa et al., 2014; Vihtelic & Hyde, 2000). In particular, altered spacing, cone mosaic patterning errors, and density differences are observed after photoreceptor ablation and regeneration (Powell et al., 2016; Raymond et al., 2014; Vihtelic & Hyde, 2000). Similarly, we noted a disordered cone terminal mosaic pattern throughout regeneration; although the detailed nature of these patterning disruptions is still unclear, it is interesting to note that they do not prevent the complete functional recovery of cone photoreceptors. Nonetheless, additional research is necessary to understand the functional significance of neuronal mosaics in adult zebrafish retina.

### Progressive Maturation of Photoreceptor Function Triggers Visual Recovery

Our observations on the progressive nature and timeline of maturation of photoreceptor function confirm and extend studies that examined visual recovery using electroretinogram (ERG) recordings and behavioural assays in teleost fish (Maurer et al., 2014; Lindsey & Powers, 2007; Kästner & Wolburg, 1982; Hammer et al., 2022; Sherpa et al., 2008; Mensinger & Powers, 2007, 1999; Sherpa et al., 2014; Wang et al., 2017; Allison et al., 2006; Barrett et al., 2022; McGinn et al., 2018; Mensinger & Powers, 1999, 2007). These studies indirectly supported the functional restoration of photoreceptors. Hammer et al. (2022) demonstrated the quantitative and gradual recovery of visual behaviour in adult zebrafish within 28 days of light lesion using optokinetic response (OKR), social preference, and light/dark preference assays. The progressive physiological recovery of cone photoreceptors over time that we observed here conceivably triggers the gradual recovery of the OKR for more difficult stimuli, i.e., with lower contrast or spatial resolution, from 14 dpl onwards (Hammer et al., 2022). Similarly, ERG studies operating at the tissue level detected partial re-establishment of ERG a- and b-wave activity following regeneration from neurotoxic lesions (McGinn et al., 2018; Barrett et al., 2022). Together, our findings argue that pre-processed physiological output signals from regenerated cones can be forwarded to complex circuits of the inner retina and brain to inform colour vision and visual behaviour.

### Why is Functional Regeneration of UV Cones Progressive?

UV cones (SWS1) in the teleost photoreceptor system, are also evolutionary homologs of mammalian ‘blue’/’S’ cones, and invaluable for functional studies in regeneration due to their high amenability and reproducibility to intense UV light lesions (Baden, 2021; Weber et al., 2013; Raymond et al., 2014). Our study evaluated UV cone function directly at the cone pedicle, revealing strong recovery in light-evoked Off responses in terms of quality, sensitivity, and kinetics after an intense light lesion. While light-induced calcium activity is present in regenerated UV cones by 8-9 dpl, quantitative analysis reveals progressive maturation to eventually achieve normal responses by 3 months. Initial UV cone Off responses (8-15 dpl) are noisy, low-amplitude, and slow, indicating reduced sensitivity in the newly regenerated eye. However, UV cone response kinetics stabilize by 15 dpl, implying an unknown ‘limiting entity’ influencing the light response quality and amplitude, which progressively improves to achieve complete functional recovery by 3 mpl. Speculatively, this limiting entity might be, for instance, a synaptic calcium channel that increases in expression over time or processes like outer segment growth, enhancing light sensitivity for reliable pedicle responses. Consistent with this, previous studies suggests photoreceptor outer segment growth during regeneration (Kramer et al., 2021; Weber et al., 2013; Hammer et al., 2022). A further possibility would be changes in cone terminal size or synaptic plasticity. Taken together, our UV cone data underscores that progressive functional regeneration is required to ultimately yield accurate physiology, reliable synaptic transmission, and robust light responses.

### Integration of Regenerated Cones Reinstates Colour Circuitry

Regenerating neurons after injury requires more than restoring their intrinsic physiological properties; new photoreceptors also need to correctly (re-)integrate into their synaptic networks. The vertebrate cone synapse is an excellent site for studying functional integration since horizontal cell-mediated synaptic transmission can invert cone responses at the pedicle as a function of space or colour in a process called cone-opponency in colour vision (Baden, 2021; Thoreson & Dacey, 2019). Consistently, larval cone pedicle tuning functions confirm that when distinct colour sensitive cones are stimulated with full-field colour stimuli, the resulting cone responses reflect their integration state within the outer retinal circuitry (Yoshimatsu et al., 2021). However, to date, no data was available in adult and regenerating zebrafish. Our findings show that the adult zebrafish UV cones in their original circuit, i.e., in the control condition, show On responses to green light, revealing a “UV vs green” colour circuit existing between UV cones and other spectral cone(s), which we utilized here to investigate circuit functionality and horizontal cell feedback during regeneration. We observed the recovery of UV cone opponency at 2.5 mpl, indicating “correct” circuit integration of regenerated UV cones into the remaining network for the first time. Further, inhibiting glutamate receptors in horizontal cells confirmed their role in outer retinal feedback to regenerated cones. It highlights the integration of regenerated cones within post-synaptic horizontal cell networks, which facilitates spectral processing in the regenerated retina. Furthermore, proper functional integration of regenerated UV cones hinges on accurately integrating other colour cones (R, G, and B) into the network; and indeed, we observed numerous light responses among the surrounding R, G, and B cones before and after regeneration, indicating their normal functional recovery. Notably, teleost R and G cones (LWS) are homologous to both mammalian ‘green’/’M’ and ‘red’/’L’ cones as well (Baden, 2021). In summary, (i) regenerated cones recover their physiological light responses, and (ii) regenerated cones are integrated into the circuitry of the outer retina, allowing the preprocessing of cone output signals via HCs, signifying that regeneration reconstructs ‘correct’ functional circuits, instead of randomly regrowing cones.

### Exploring Adult Zebrafish Retinal Organization with 2P imaging

To date, most of what is known about the functional organization in the zebrafish retina stems from *in vivo* 2P imaging in larval retina (Baden, 2021), partly due to technical constraints of studying a light sensor (retina), with light microscopy in its flat-mounted configuration (Euler et al., 2019). Our 2P recording of light-driven responses from cone photoreceptors and its combination with light lesion and regeneration is unique for adult zebrafish. It combines excellent resolution of the cellular substrate with accessibility to many neurons, making it a novel and valuable addition to the existing toolkit of functional studies, which would otherwise rely on systemic measurements at lower than cellular resolution. The UV cone hyperpolarization (Off) to blue light that we report in adult retina agrees with the observations in larval retina, where only G and B cones are spectrally opponent, while UV and R are non-opponent (Yoshimatsu et al., 2021). However, the strong UV cone depolarization (On) response to green light we see in adults is substantially weaker and regionally restricted in larvae, possibly reflecting the progressive maturation of neuronal circuits from larval to fully adult stages. Also, the stereotypic cone mosaic arrangement that is characteristic of the adult zebrafish retina is still absent in larval stages (Allison et al., 2010). Thus, our findings open fresh avenues for studying functional organization of neuronal circuits during development.

### Future Directions

In this work, we have shown that zebrafish cone photoreceptors achieve not only anatomical regeneration in response to injury, but also appropriate physiology and functional integration within the outer retinal circuits in the adult retina using 2P calcium imaging. Our findings offer new avenues of exploration, including understanding retinal signal propagation and re-establishment of retinal networks to assess functional recovery in retinal processing at the inner plexiform layer, where intricate computations occur, and at the functional retinal ganglion cell output channels level. These investigations illuminate functional recovery in retinal contexts and, since the retina is essentially a developmentally everted piece of diencephalon, may apply more generally to the regenerating zebrafish brain (Kyritsis et al., 2012; Kroehne et al., 2011). In conclusion, endogenous-regeneration-based strategies offer significant potential for advancing therapies in neurodegenerative diseases using mammalian models and potentially benefiting human treatments.

## Supporting information

Movies 2 Photon

## Movies

### SyGCaMP6f light response and corresponding activity (grey)

Activity (see methods) shows synaptic cone SyGCaMP6f calcium responses to blue and green stimulus respectively. Dark terminals represent Off responses, whereas lighter terminals represent On responses.

Stimulus: Flash ON for 1 sec, and Flash OFF for 1 sec.

Total duration: 2 sec.

**Movie 1:**
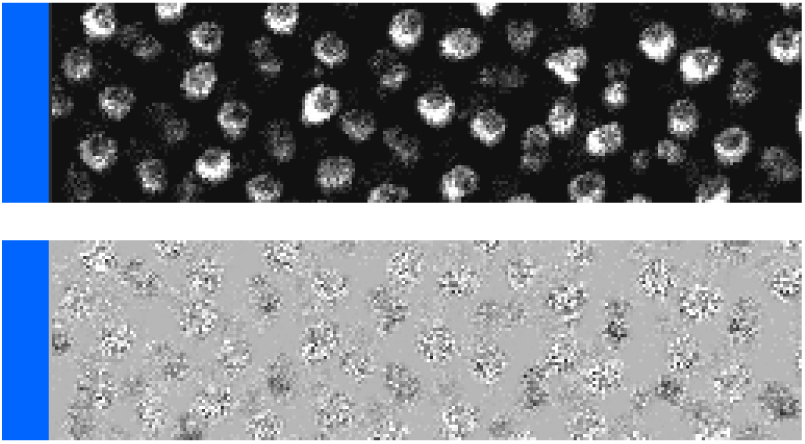
Fig. 1D, Control_1 *(blue, 419 nm)*

**Movie 2:**
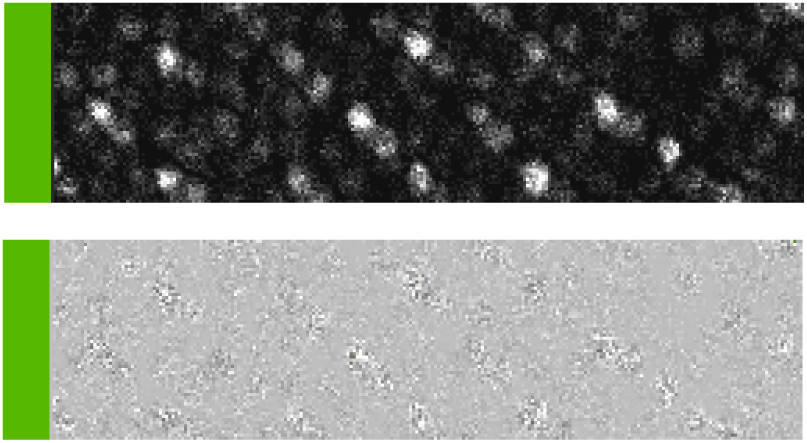
Fig. 1F, Control_2 *(green, 484 nm)*

**Movie 3:**
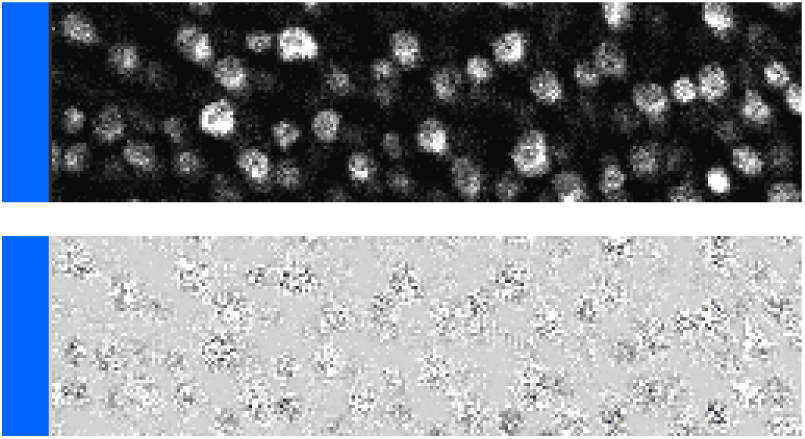
Fig. 2D, Control_3 *(blue, 419 nm)*

**Movie 4:**
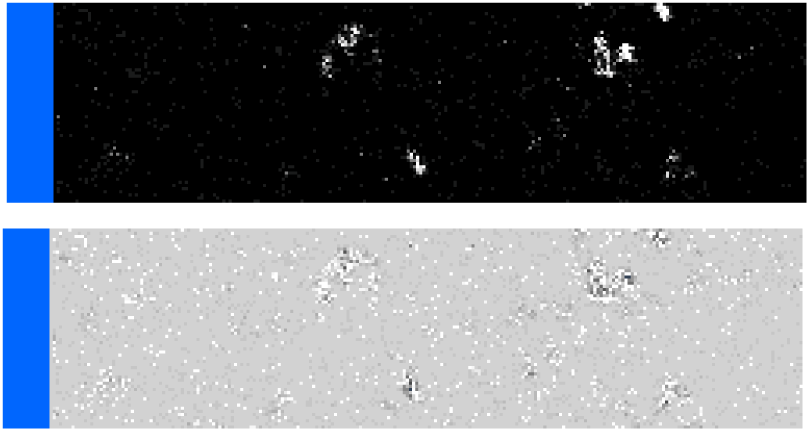
Fig. 2D, 4 dpl *(blue, 419 nm)*

**Movie 5:**
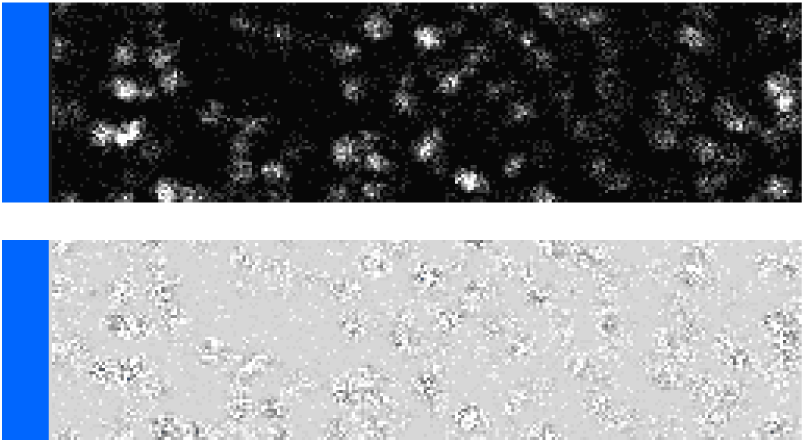
Fig. 2D, 8-9 dpl *(blue, 419 nm)*

**Movie 6:**
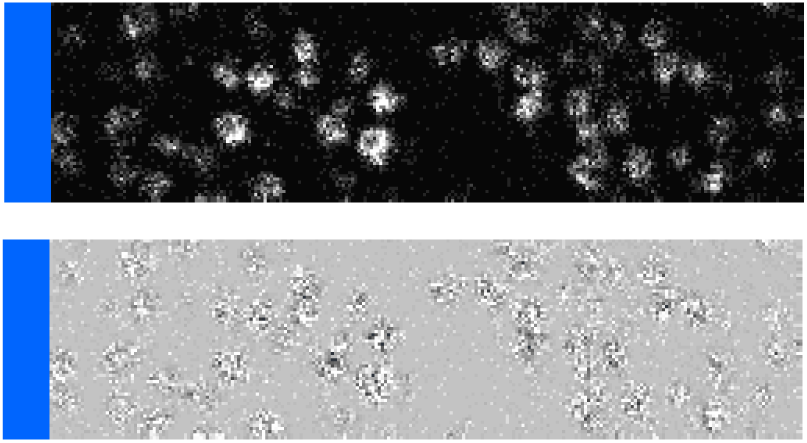
Fig. 2D, 10-11 dpl *(blue, 419 nm)*

**Movie 7:**
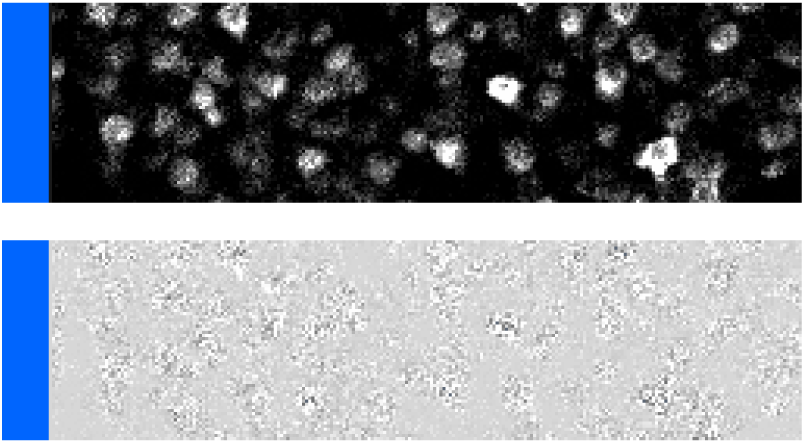
Fig. 2D, 14-15 dpl *(blue, 419 nm)*

**Movie 8:**
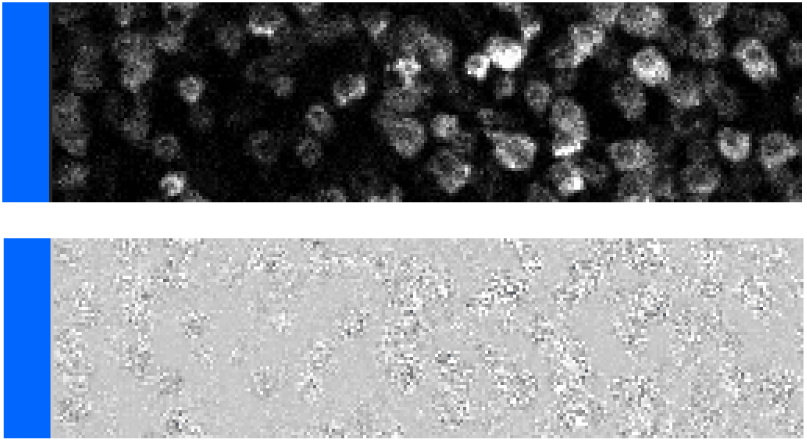
Fig. 2D, 28-29 dpl *(blue, 419 nm)*

**Movie 9:**
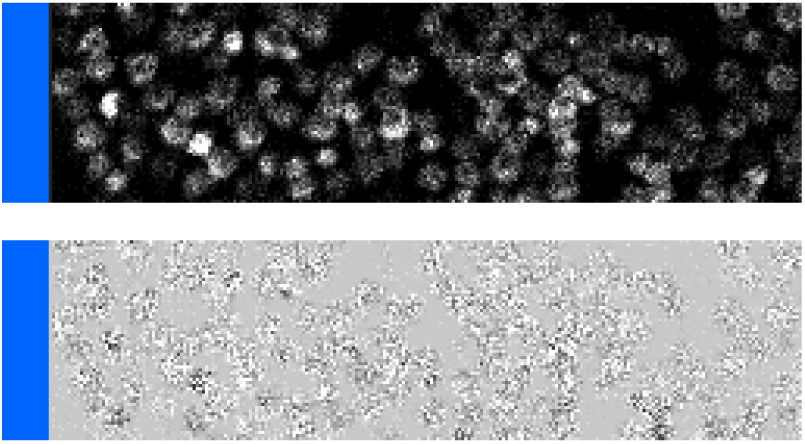
Fig. 2D, 3 mpl *(blue, 419 nm)*

**Movie 10:**
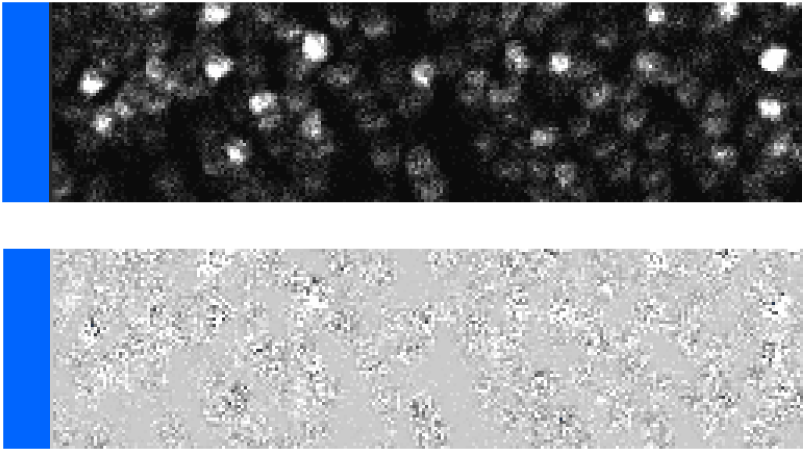
Fig. 2D. 8 mpl *(blue, 419 nm)*

**Movie 11:**
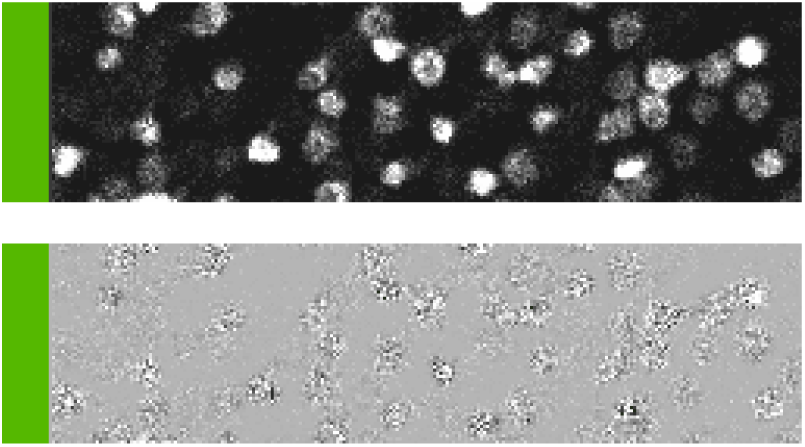
Fig. S6A, 2.5 mpl_1 *(green, 484 nm)*

**Movie 12:**
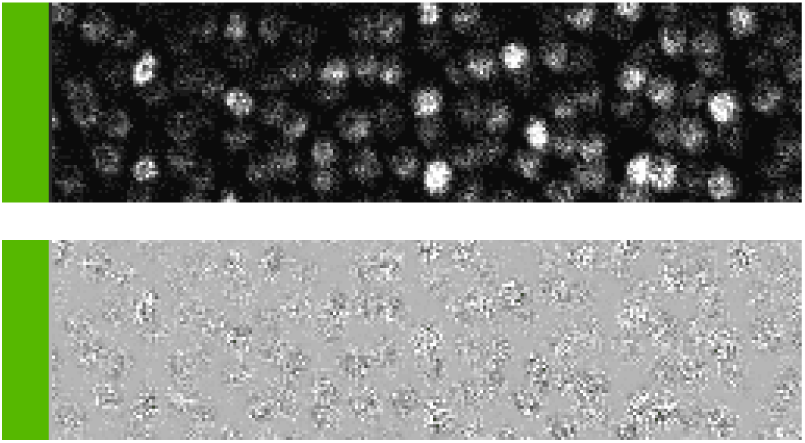
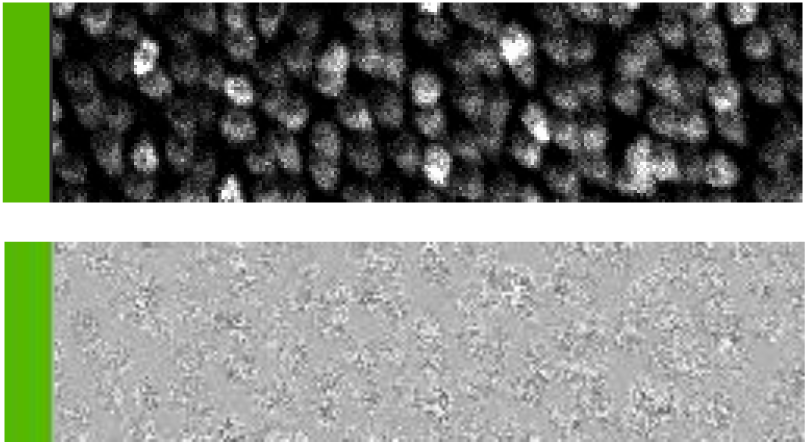
Fig. 4D, Control_4 +CNQX *(green, 484 nm)*

**Movie 13:**
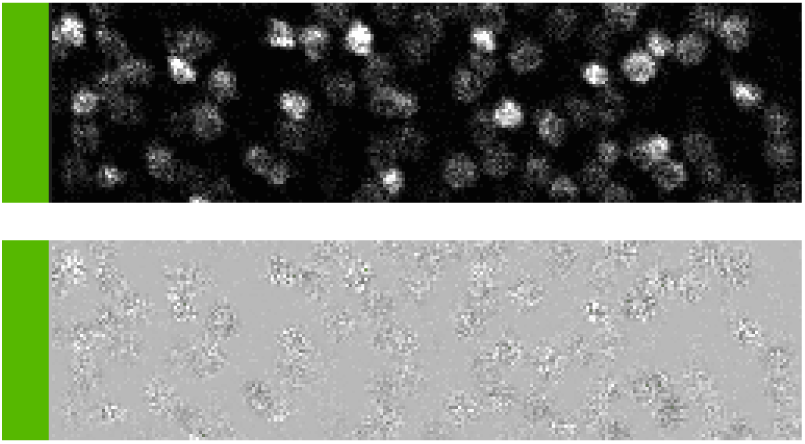
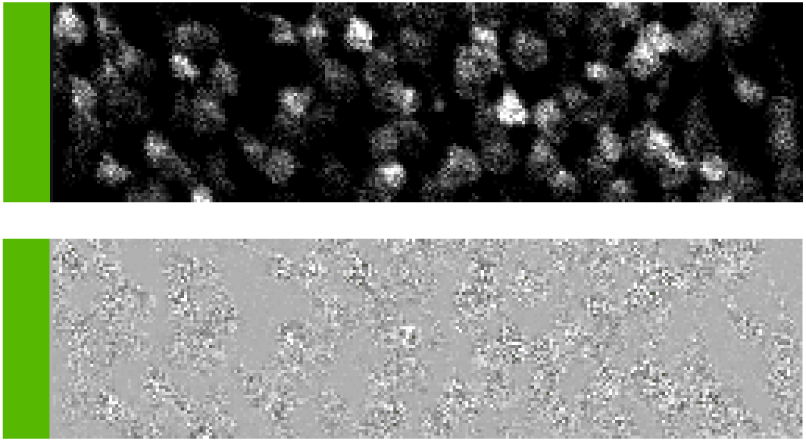
Fig. 4F, 2.5 mpl_2 + CNQX *(green, 484 nm)*

## Acknowledgements

We thank all members of the Brand laboratory for their continued support, scientific discussions, as well as helpful comments on this project. We thank Judith Konantz, M. Fischer, S. Kunadt and D. Mögel for excellent zebrafish care. We would particularly like to thank Falk Hartmann for developing, with H.H., the electronics, and custom software for controlling the LED light stimulation synchronized to the 2-photon confocal laser scanning measurements, as part of our Light Microscopy Facility, a core facility of the CMCB at TU Dresden. Aristides Arrenberg, MPI Tübingen, kindly gave us an earlier GCaMP6f fish line. This work was supported by grants to M.B. from the Deutsche Forschungsgemeinschaft (BR 1746/11-1), the European Union (European Research Council AdG Zf-BrainReg), the German Excellence Initiative (Wissenschaftsrat and Deutsche Forschungsgemeinschaft) (EXC 168), and by funds from TU Dresden. The funders had no role in study design, data collection and analysis, decision to publish, or preparation of the manuscript.

## Author contributions

Conceptualization: M.B., E.A.; Investigation: E.A.; Analysis: E.A., Writing – original draft: E.A., M.B.; Writing - review & editing: E.A., H.H., T.Y., T.B. and M.B.; Supervision: M.B.; Funding acquisition: M.B.

## SUPPLEMENTAL INFORMATION

Supplemental Information can be found online.

## STAR METHODS

### KEY RESOURCES TABLE

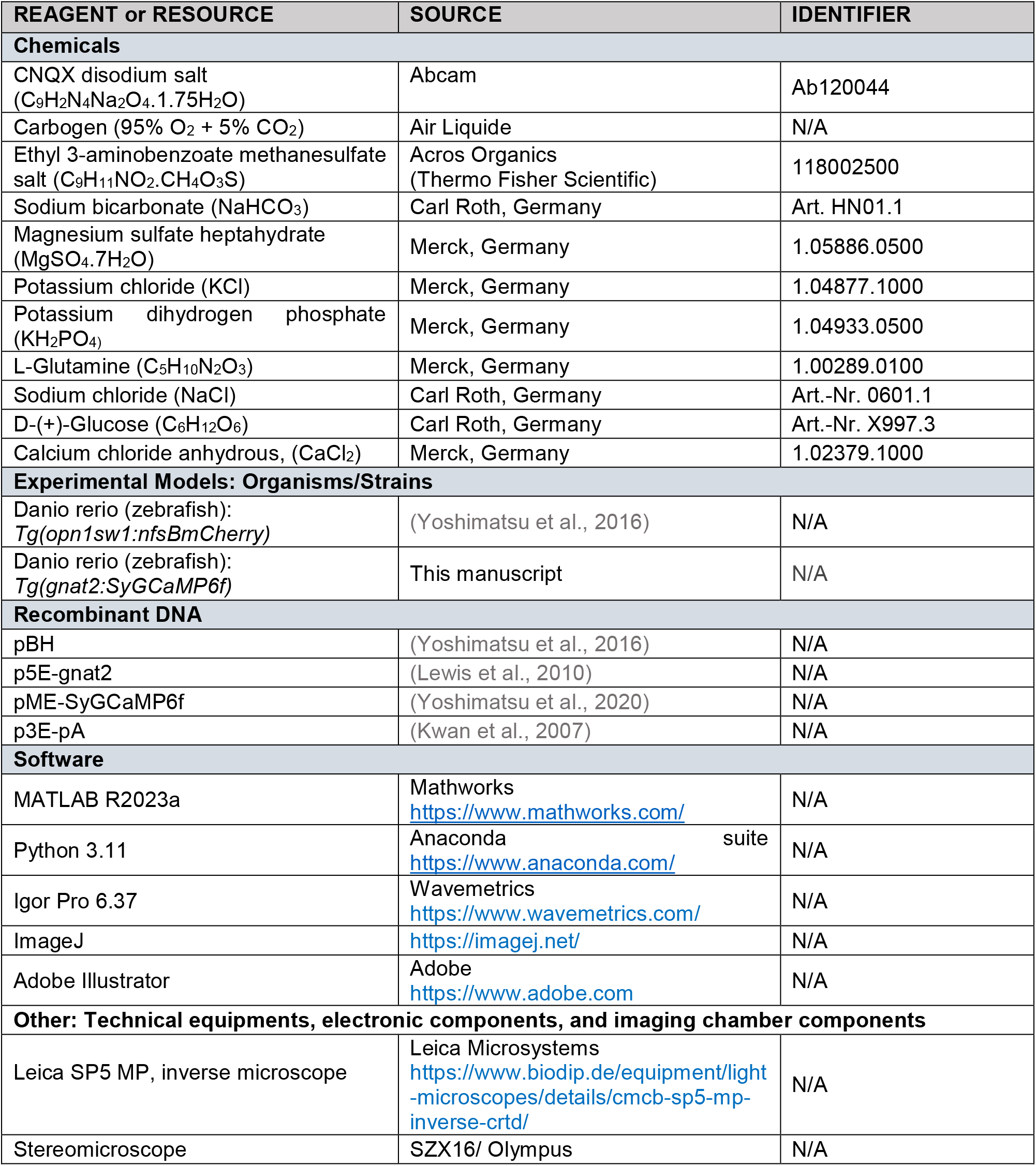
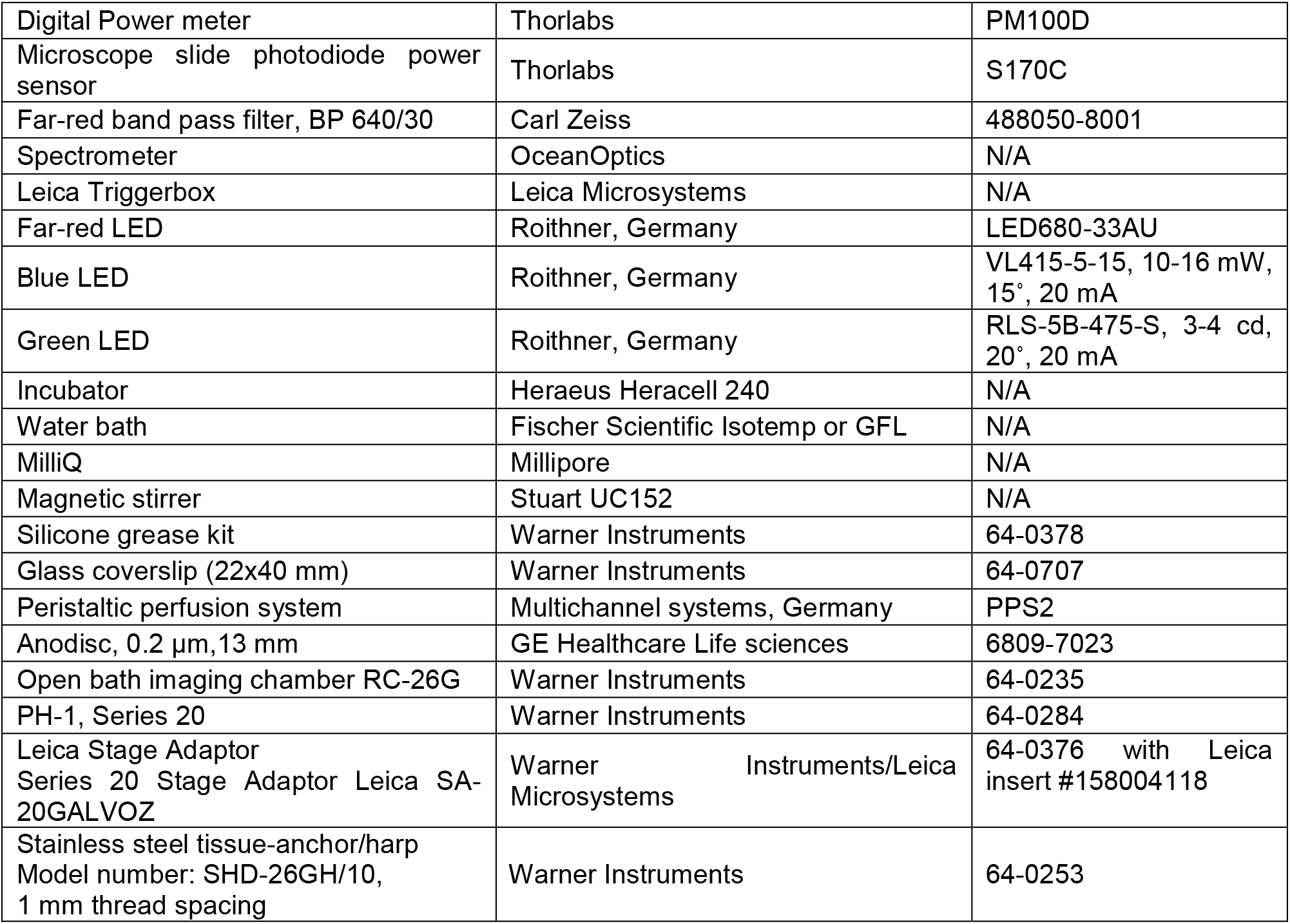

| REAGENT or RESOURCE | SOURCE | IDENTIFIER |
| --- | --- | --- |
| **Chemicals** |  |  |
| CNQX disodium salt (C<sub>9</sub>H<sub>2</sub>N<sub>4</sub>Na<sub>2</sub>O<sub>4</sub>·1.75H<sub>2</sub>O) | Abcam | Ab120044 |
| Carbogen (95% O<sub>2</sub> + 5% CO<sub>2</sub>) | Air Liquide | N/A |
| Ethyl 3-aminobenzoate methanesulfate salt (C<sub>9</sub>H<sub>11</sub>NO<sub>2</sub>·CH<sub>4</sub>O<sub>3</sub>S) | Acros Organics (Thermo Fisher Scientific) | 118002500 |
| Sodium bicarbonate (NaHCO<sub>3</sub>) | Carl Roth, Germany | Art. HN01.1 |
| Magnesium sulfate heptahydrate (MgSO<sub>4</sub>·7H<sub>2</sub>O) | Merck, Germany | 1.05886.0500 |
| Potassium chloride (KCl) | Merck, Germany | 1.04877.1000 |
| Potassium dihydrogen phosphate (KH<sub>2</sub>PO<sub>4</sub>) | Merck, Germany | 1.04933.0500 |
| L-Glutamine (C<sub>5</sub>H<sub>10</sub>N<sub>2</sub>O<sub>3</sub>) | Merck, Germany | 1.00289.0100 |
| Sodium chloride (NaCl) | Carl Roth, Germany | Art.-Nr. 0601.1 |
| D-(+)-Glucose (C<sub>6</sub>H<sub>12</sub>O<sub>6</sub>) | Carl Roth, Germany | Art.-Nr. X997.3 |
| Calcium chloride anhydrous, (CaCl<sub>2</sub>) | Merck, Germany | 1.02379.1000 |
| **Experimental Models: Organisms/Strains** |  |  |
| Danio rerio (zebrafish): *Tg(opn1sw1:nfsBmCherry)* | (Yoshimatsu et al., 2016) | N/A |
| Danio rerio (zebrafish): *Tg(gnat2:SyGCaMP6f)* | This manuscript | N/A |
| **Recombinant DNA** |  |  |
| pBH | (Yoshimatsu et al., 2016) | N/A |
| p5E-gnat2 | (Lewis et al., 2010) | N/A |
| pME-SyGCaMP6f | (Yoshimatsu et al., 2020) | N/A |
| p3E-pA | (Kwan et al., 2007) | N/A |
| **Software** |  |  |
| MATLAB R2023a | Mathworks<br>https://www.mathworks.com/ | N/A |
| Python 3.11 | Anaconda suite<br>https://www.anaconda.com/ | N/A |
| Igor Pro 6.37 | Wavemetrics<br>https://www.wavemetrics.com/ | N/A |
| ImageJ | https://imagej.net/ | N/A |
| Adobe Illustrator | Adobe<br>https://www.adobe.com | N/A |
| **Other: Technical equipments, electronic components, and imaging chamber components** |  |  |
| Leica SP5 MP, inverse microscope | Leica Microsystems<br>https://www.biodip.de/equipment/light-microscopes/details/cmcb-sp5-mp-inverse-crtd/ | N/A |
| Stereomicroscope | SZX16/ Olympus | N/A |
| Digital Power meter | Thorlabs | PM100D |
| Microscope slide photodiode power sensor | Thorlabs | S170C |
| Far-red band pass filter, BP 640/30 | Carl Zeiss | 488050-8001 |
| Spectrometer | OceanOptics | N/A |
| Leica Triggerbox | Leica Microsystems | N/A |
| Far-red LED | Roithner, Germany | LED680-33AU |
| Blue LED | Roithner, Germany | VL415-5-15, 10-16 mW, 15°, 20 mA |
| Green LED | Roithner, Germany | RLS-5B-475-S, 3-4 cd, 20°, 20 mA |
| Incubator | Heraeus Heracell 240 | N/A |
| Water bath | Fischer Scientific Isotemp or GFL | N/A |
| MilliQ | Millipore | N/A |
| Magnetic stirrer | Stuart UC152 | N/A |
| Silicone grease kit | Warner Instruments | 64-0378 |
| Glass coverslip (22x40 mm) | Warner Instruments | 64-0707 |
| Peristaltic perfusion system | Multichannel systems, Germany | PPS2 |
| Anodisc, 0.2 µm, 13 mm | GE Healthcare Life sciences | 6809-7023 |
| Open bath imaging chamber RC-26G | Warner Instruments | 64-0235 |
| PH-1, Series 20 | Warner Instruments | 64-0284 |
| Leica Stage Adaptor Series 20 Stage Adaptor Leica SA-20GALVOZ | Warner Instruments/Leica Microsystems | 64-0376 with Leica insert #158004118 |
| Stainless steel tissue-anchor/harp Model number: SHD-26GH/10, 1 mm thread spacing | Warner Instruments | 64-0253 |

### RESOURCE AVAILABILITY

#### Lead Contact

Further information and requests for resources and reagents should be directed to and will be fulfilled by the Lead Contact, Michael Brand, CRTD, TU Dresden, Germany.

#### Materials Availability

Materials generated in this study are available upon request to the Lead Contact.

#### Data and Code Availability

Processed calcium responses, data and codes used for generating figures are deposited online. Raw 2P imaging datasets, and any remaining data used in this study will be provided by the Lead Contact upon reasonable requests.

### EXPERIMENTAL MODEL AND SUBJECT DETAILS

#### Ethics statement

All animal experiments in this study were performed according to the animal welfare legislation and were approved by the animal ethics committee of the TU Dresden and the Landesdirektion Sachsen (Permit: TVV 59/2018). All possible measures were taken to reduce animal suffering and limit the number of animals utilized.

#### Animals

Zebrafish (*Danio rerio*) were kept as previously described (Brand et al., 2002) in our fish facility in a light-dark alternation (14:10 hr cycle) per day. Transgenic embryos were sorted at 4-5 dpf for red heart marker (*gnat2:SyGCaMP6f transgene*) and red eye marker (*opn1sw1:nfsb-mCherry transgene*) and raised until adulthood. *Ex vivo* functional recordings for all 2P imaging experiments were conducted using a mixed group of adult zebrafish, both female and male, in the double transgenic *Tg(gnat2:SyGCaMP6f); Tg(opn1sw1:nfsb-mCherry)* AB background. The zebrafish selected were aged between 6-18 months post-fertilization. Experimental group names were approximated to either days post lesion (dpl) or months post lesion (mpl) depending on the harvested time points.

### METHODS

#### Generation of Transgenic lines

*Tg(gnat2:SyGCaMP6f)* line was generated by T.Y. by injecting pBH-gnat2-SyGCaMP6f-pA plasmids into single-cell stage eggs. Injected fish were out-crossed with wild-type fish to screen for founders. Progeny was raised to establish transgenic lines. The plasmid was made using the Gateway system (ThermoFisher, 12538120) with combinations of entry and destination plasmids as follows: pBH (Yoshimatsu et al., 2016), p5E-gnat2 (Lewis et al., 2010), pME-SyGCaMP6f (Yoshimatsu et al., 2020), p3E-pA (Kwan et al., 2007).

#### Diffuse light lesion

Adult zebrafish were continuously dark adapted for 3 days prior to light lesion to maximize light sensitivity. Diffuse light lesions were performed as described in (Weber et al., 2013) consistently with N = 6 fish each at a time, freely swimming in a 400 ml beaker for all experiments. Aluminum foil was wrapped around the beaker to reflect the incoming light, leaving enough width (∼5 cm) for light entry. A bright UV-rich light source (∼200,000 lux) EXFO X-Cite 120W metal halide lamp (EXFO Photonic Solutions, Mississauga, Canada) was projected via a light guide carefully fixed at a horizontal distance between 0.5-1 cm at the beaker’s half-height. The light source was switched off after 30 minutes, and the lesioned fish were returned to standard fish facility lighting/housing.

#### Physiological solution

Ames medium formulated to support retinal tissue in relatively short-term culture was used for live-imaging zebrafish retinal explants. 2 litre basic Ames buffer was freshly prepared in deionized water at room temperature. Composition for 2 litre Ames buffer: NaHCO_3_ (2.8g/16.7mM), CaCl_2_ (0.255g/1.2mM), MgSO_4_.7H_2_O (0.609g/1.2 mM), KCl (0.462g/3.1mM), KH_2_PO_4_ (0.136g/499.7 μM), L-Glutamine (0.146g/499.5 μM), NaCl (14.02g/120 mM), C_6_H_12_O_6_, D-(+)-Glucose (2.16g/6mM). The prepared Ames media was mixed for ≥ 30 min on a magnetic stirrer until the salts were dissolved, following which it was incubated in a 28-degree water bath and bubbled continuously with carbogen (95% O_2_ and 5% CO_2_) prior to start of experiments.

#### Retinal explant preparation and mounting

Experimental animals were dark adapted for at-least 3 hours before starting all functional recordings. All procedures were conducted in a dark room under dim red illumination (> 640 nm). Single adult zebrafish were anesthetized in 0.02% MS-222 until cessation of opercular movements, followed by decapitation. The head was immediately immersed in 30°C warm Ames buffer and bubbled continuously with 95% O_2_ and 5% CO_2_ in a petri dish. Eyes were removed surgically by excision of the optic nerve and quickly transferred into another petri dish filled with continuously oxygenated Ames. The retina was dissected from the eyecup, Retinal Pigment Epithelium removed, and transferred onto the glass coverslip of the diamond open bath recording chamber (Warner instruments) filled with Ames media. The retinal ganglion cell side aligned with the coverslip. Next, an anodisc (GE Healthcare) filter membrane was placed on the photoreceptor side from the top to flatten the retina and hold it down. The retina was further stabilized with a U-shaped harp (Warner instruments) with nylon strings of 1 mm spacing placed over the anodisc. Next, the infusion and suction cannulas for tissue perfusion were tightly positioned, and the whole chamber was mounted on the microscope stage (Leica). The microscope was incubated at 28 degrees in a custom temperature control unit (Leica). A perfusion pump (Multichannel systems) supplied freshly oxygenated Ames media (maintained at ∼30°C) at 3ml/min, and the waste media was removed at the rate of 3.5ml/min. The retina was focused to position under far-red illumination (∼680 nm) through the microscope eye piece before the start of 2P imaging.

#### Pharmacology

Horizontal cell block was performed using the AMPA/kainate antagonist CNQX. 2mM working solutions of CNQX were freshly prepared in 1X Ames media, and aliquots were incubated at 37 degrees to ensure maximum solubility. Perfusion was briefly switched off prior to drug addition, and 200 microlitre warm, oxygenated CNQX solution was directly added into the open bath chamber with the retinal explant and mixed with a 200-microliter pipette under dim red illumination. Post-drug recordings were acquired from the undisturbed unique spot after CNQX addition, following which perfusion was switched on.

#### Two-photon calcium imaging

For *ex vivo* recordings, an inverse Leica SP5 MP two-photon microscope equipped with a Spectra-Physics MaiTai Ti: Sapphire laser tuned to 920 nm was used. For image acquisition, Leica LAS AF software, a water immersion objective (25x/0.95 HC IRAPO L, Leica), descanned HyD detectors allowing spectral detection for imaging SyGCaMP6f (495-560 nm) and mCherry (609-700 nm) was employed. For the 4 dpl sample and in a few cases mCherry was detected within 650-700 nm due to technical issues. 3-6% of 2P laser power, corresponding to a displayed ∼0.87 Watts in LAS AF software, was required for the SyGCaMP6f signal. The tissue was kept under 2P scanning for at least 5-6 seconds before light stimulus presentations for all functional recordings. A consistent protocol in which either the blue or green LED was switched ON for 1 sec (flash) followed by 1 sec OFF (interval) of 2P scanning only, with 10 repeats is defined was the experimental paradigm to study light-correlated activity in cones. Morphological data was recorded at 512×512 pixels (frame average 12 or 20), while functional data was recorded at 256×64 pixel resolution (conventional scanner, 1000 Hz line frequency, unidirectional scan, with no frame averaging) at 70ms per frame (∼14.28Hz).

#### Light stimulation

A custom-built stimulator unit, allowing for Köhler illumination of stimulus light through the condenser lens using the transmitted light arm of the microscope, was used for light stimulation. Single light emitting diodes (‘blue’ 419 nm, VL415-5-15, 10-16 mW, 15°, 20 mA; and ‘green’ 484 nm, RLS-5B-475-S, 3-4 cd, 20°, 20 mA; Roithner, Germany) were mounted on the optical axis within a Leica halogen lamp house using a 3D printed holder. The field diaphragm was adjusted to restrict stimulus size to 400 microns. The aperture diaphragm was minimized in steps to decrease light intensity further. LED stimulations were synchronized with the line scan retrace at 1000Hz to prevent interference of the stimulation light with the fluorescence recordings (Euler et al., 2019). An Adafruit HUZZAH ESP32 microcontroller and custom software (via a browser interface) were used to set the stimulation sequence. All associated build files used for the light stimulator are available on bitbucket. The stimulus light was focused on the photoreceptors, through the condenser with Köhler illumination prior to functional recordings.

#### Stimulus intensities

The blue (419 nm) and green (484 nm) LED spectra approximated the peak spectral sensitivity of zebrafish B- and G-opsins, respectively, with no additional bandpass filters. Effective stimulus light intensity during scanning (256×64 pixels) was measured with a power meter (Thorlabs PM100D equipped with an S106 photodiode sensor) to measure effective power (in μWatt) passing through the anodisc at the sample location just above the objective lens. The power values were directly read from the instrument after zeroing the background before turning ON the LEDs. The power values were calibrated into radiometric units (photons/sec/m^2^) at specific wavelengths (blue-419 nm, green-484 nm). First, the energy of a single blue/green photon was calculated (in eV) at respective wavelengths using the Planck-Einstein relation: E= (1.2398)/(λ (in μm)); where λ is the LED wavelength in μm. The calculated E is converted into Joules (J) using the relation 1 eV = 1.602176634 x 10-19 J. Next, the number of photons striking the sensor per sec is calculated by taking the ratio of measured LED power (in Watt) and E (in J). Since the stimulus size is defined by Koehler illumination, the area enclosed by a circular region of radius 200 μm on the sensor was computed. From here, the number of photons striking the sensor per sec per unit area per nm (photons/sec/m^2^/nm) for respective blue and green LEDs was calculated. With the measured power of blue LED at 0.06 μW and green LED at 0.02 μW at the lowest aperture diaphragm position, the final power values are at 1.01e+18 photons/sec/m^2^/nm for blue LED and at 3.88e+17 photons/sec/m^2^/nm for green LED. The LED stimulus artifact detected by the microscope detectors during functional imaging and visible in the imaging datasets is cueing for temporal alignment of neural activity with stimulus onset for data analysis. Precisely aligning the light stimulus with imaging data is met using the LED light spectrally detected during imaging and embedded in the data. Zebrafish opsin template values are provided by T.Y and extracted using (Govardovskii et al., 2000).

### QUANTIFICATION AND STATISTICAL ANALYSIS

#### Image processing

Functional scan fields are represented as AVG (average) projection of the raw TIFF imaging time series stack to show co-localization of UV cones (mCherry) with SyGCaMP6f activity. The standard deviation (STD) projections of raw SyGCaMP6f functional scans were used for placing UV and surrounding non-UV cone regions of interest (ROIs) for further analysis of Ca^++^ traces in Igor Pro. Activity images of functional scans were generated using the averaged movies (28 frames) exported from Igor Pro: the AVG projection of the last 4 frames (25-28) was subtracted from the raw TIFF averaged stack in FiJi. Further, the subtracted stack was AVG projected (frames 5-20) and processed in FiJi for similar background shade (grey). Darker than background (black) terminals represent Off responses, whereas lighter than background (white) terminals represent On responses. For creating light response movies with stimulus bars, a 28 frame RGB stack of 15×64 pixels, where the first 14 frames were colour coded for either blue or green was combined with the SyGCaMP6f RGB stack of 241×64 pixels. All 2P microscopy images are processed in FiJi. Final figures are compiled using Adobe Illustrator.

#### Analysis

Functional recordings (raw TIFF data) were pre-processed in Igor Pro using custom-written routines. The recordings were combined systematically by time points, recording zone (central or ventral), and stimulus and retina identity. All cone regions of interest ROIs are placed manually by hand in Igor Pro on the standard deviation projection of SyGCaMP6f time series channel to best visualize all morphologically present cones. The extracted response quality, stimulus temporal information, and cone activity information were post-processed in Python and MATLAB using custom-written scripts. In all experimental groups, only ‘responsive’ scans are included, where at least one cone ‘responded (Qi >= 0.4)’ to the stimulus. All data visualization is done in MATLAB. Final figures are compiled using Adobe Illustrator.

#### Data pre-processing

Calcium (Ca++) traces for each ROI were extracted at 1 ms precision based on the centre of mass of each ROI. The traces were z-normalized based on the 1-6 sec time interval at the beginning of recordings. The stimulus timing was detected in the red or blue channel and used to align the Ca++ traces relative to the visual stimulus with a temporal precision of 1 ms. For quantifying response amplitudes and Tau, the averaged Ca++ traces from 10 repetitive visual stimuli were used by aligning single trial traces at 1 ms precision. Example Ca++ response movies were generated by averaging Ca++ traces for each pixel in the scan images.

The quality index was computed as follows:

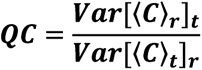

where C is the T by R response matrix (time samples by stimulus repetitions) and 〈 〉x and Var[]x denotes the mean and variance across the indicated dimension, respectively. If all trials are identical such that the mean response perfectly represents the response, QC equals 1. If all trials are random with fixed variance, QC equals 1/R. All pre-processing was done using custom-written routines under IGOR Pro (Yoshimatsu et al., 2021)

#### Data post-processing

*Exponential fit:* A quality index (Qi) was used to estimate the reliability of light responses among the 10 trials. Qi >= 0.4 cutoff was used to assign cones that responded to light and were considered as the ‘responsive’ fraction that gave reliable responses for further analysis. The remaining cones were grouped as the ‘non-responsive’ cones. For all analysis, the averaged cone response (in Z scores) is used.

For kinetics, single exponential function of the form:

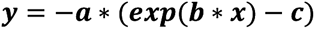

are fit separately on the average response (in Z scores) decay and recovery phases of the Off response (and similarly for the On response) to calculate the best-fit estimates of amplitude and speed, Tau. The amplitude was calculated as the fitted exponential curve’s value corresponding to the average response’s maximum value. The best fit Tau was calculated as the sign-independent inverse value of the best-fit rate parameter ‘b’. To determine Off (hyperpolarization to light) and On (depolarization to light) responses (as used in percentage estimations in bar charts), the averaged curve is offset to start at zero and values summated (sum). ‘Off’ responses are grouped as curves with sum negative and amplitude positive. On responses are grouped are curves with sum positive and amplitude positive. For kinetics analysis of Off or On responses, the averaged curves are first sorted into Off or On group and only the responses satisfying the coefficient of determination (R²) cutoff 0.8 are further considered. For analysis of amplitude and Tau, a 2.5 x median absolute deviation window (MAD) is used to exclude extreme outliers in all samples in the time course, first applied to Tau recovery, then to Tau decay (Leys et al., 2013). These outliers represent extremely slow cone relaxations and violate the exponential model, becoming more linear. These slow and noisy responders are classified as non-physiological responses (outliers).

#### Statistics

No analysis was done to predetermine sample size/power. One-way analysis of variance or ANOVA followed by Tukey Kramer post-hoc analysis for unequal sample sizes was used whenever multiple pairwise comparisons of independent control and multiple experimental groups were done. ANOVA results prior to Tukey Kramer post-hoc analysis for unequal sample sizes summarized in respective tables. For comparing two independent groups, a Two-sample t-test was used. Summary of groups, sample size, mean, 95% confidence interval (95% CI) of mean, pairwise comparisons, p values, and effect size of the difference for each comparison in terms of Cohen’s d (d value) are reported in respective tables. All Statistical tests are done in MATLAB (Statistics toolbox) with custom scripts.

## Supplementary information

### SUPPLEMENTARY FIGURES

**Figure S1:**
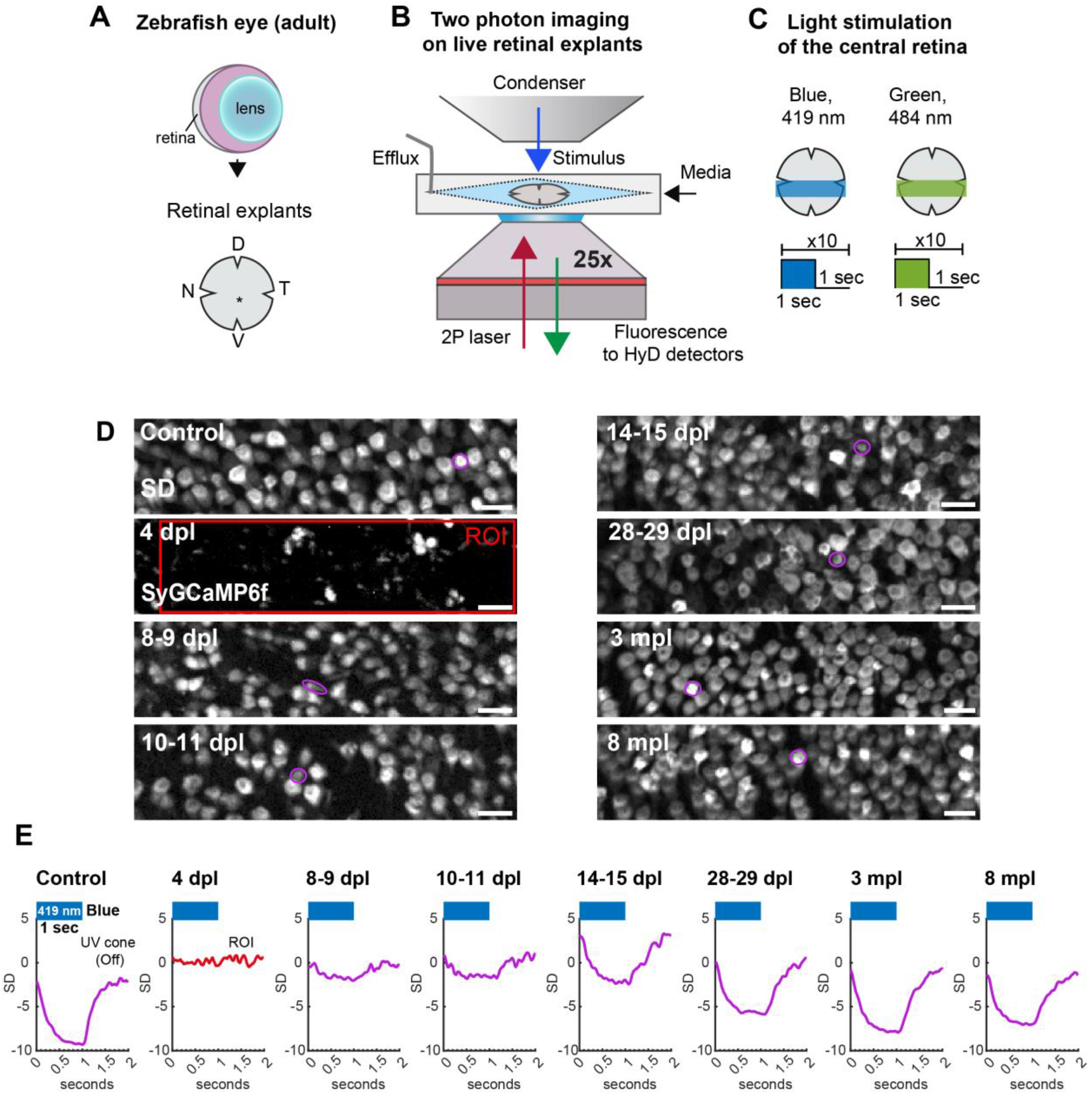
(A-C) 2P calcium imaging with light stimulation in adult zebrafish retinal explants. **(A)** Zebrafish adult retinal whole mount preparation (explants), orientation preserved. **(B)** Retinal explant live-imaging with continuous oxygenated media perfusion under an inverse 2P microscope. 2P excitation (red arrow), light stimulus through condenser (blue arrow), emitted fluorescence (green arrow). **(C)** Light stimulation protocol: LED ON for 1 second and interval for 1 sec, repeated 10 times for each scan field in the central retina. nasal (N), temporal (T), dorsal (D), and ventral (V). **(D-E) UV cone functional recovery in the central retina at the single cone level | (D)** Standard deviation (SD) projection of SyGCaMP6f imaging stack from representative scans shown in Fig. 2D. SD projection is used for placing cone ROIs; here single UV cone ROIs are shown encircled in magenta. At 4 dpl, cones are still ablated in the central retina, with only debris remaining; large rectangular ROI shown in red. Note that non-UV cones ROIs can also be analyzed in this way. **(E)** Light responses of single UV cones from A. Blue bar indicates blue LED stimulus presentation for 1 sec. Control, and 8-9 dpl - 8 mpl shows UV cone Off responses, whereas 4 dpl shows loss of function or random noise. Y-axis: standard deviation (SD) and X-axis: time (seconds). Grey: SyGCaMP6f signal. Scale bar = 10 micrometre. See also Fig. 1-2.

**Figure S2:**
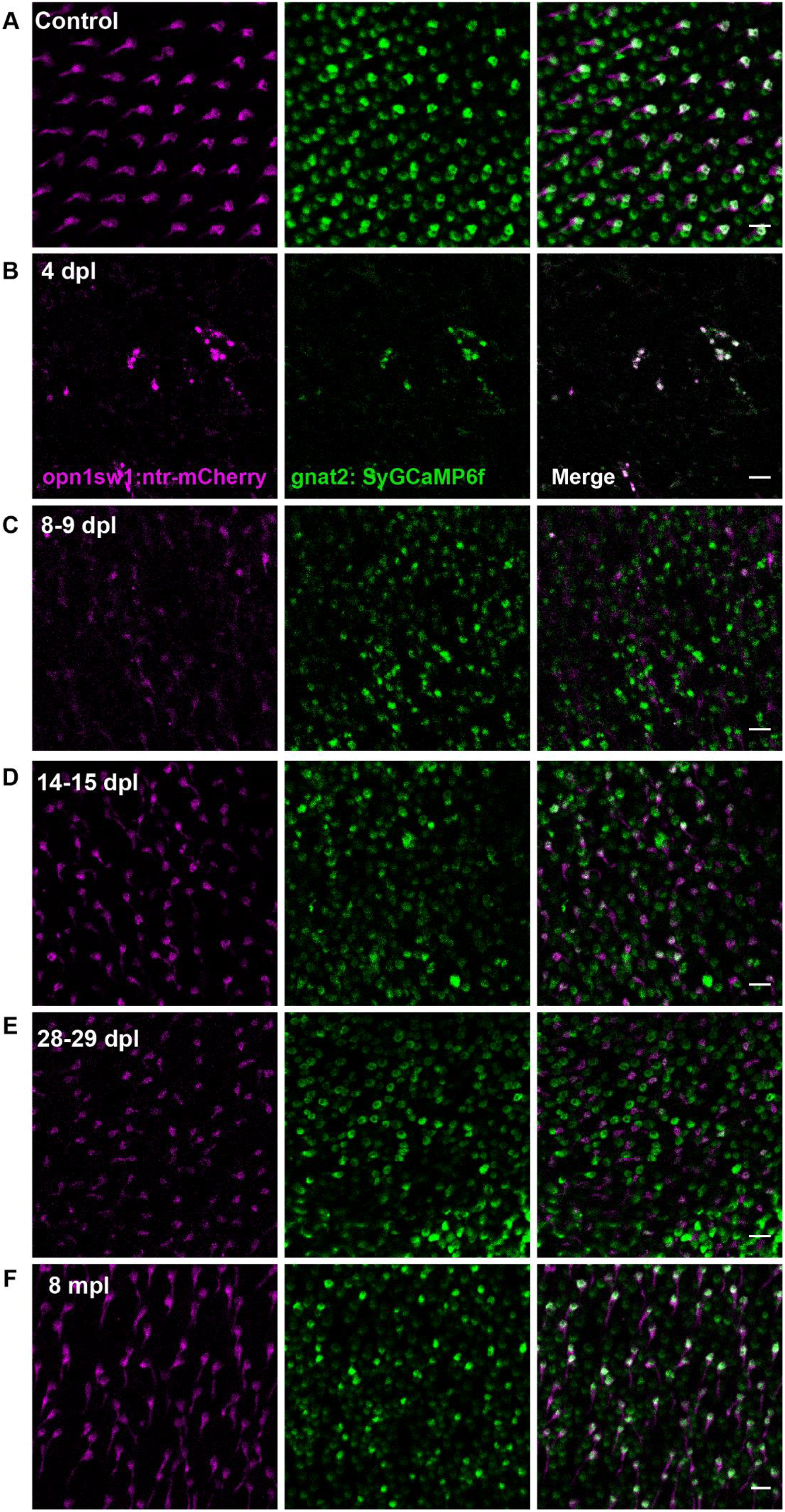
Cone photoreceptors in the central retina are completely ablated after light lesion, but continue to regenerate from 8 dpl to 8 mpl-exemplary morphological scans shown. Regenerating cones in the central retina in double transgenic *Tg(gnat2:SyGCaMP6f); Tg(opn1sw1:nfsb-mCherry)* reporter after diffuse light lesion **(A)** Control, **(B)** 4 dpl, **(C)** 8-9 dpl, **(D)** 14-15 dpl, **(E)** 28-29 dpl, **(F)** 8 mpl. Several morphological scan fields from respective time points were examined during functional recordings, and representative images are shown. Compared to controls, 4 dpl central retina is completely ablated of UV cones and the bulk of blue, green and red cones. The bulk of regenerated cones are captured in the central retina where the UV cone row mosaic is disrupted, starting at 8-9 dpl and remains prominent at all other examined time points 10-11 dpl (not shown), 14-15 dpl (shown), 28-29 dpl (shown), 3 mpl (not shown) and 8 mpl (shown). Green: SyGCaMP6f, magenta: UV cone mCherry. Scale bar = 10 micrometre.

**Figure S3:**
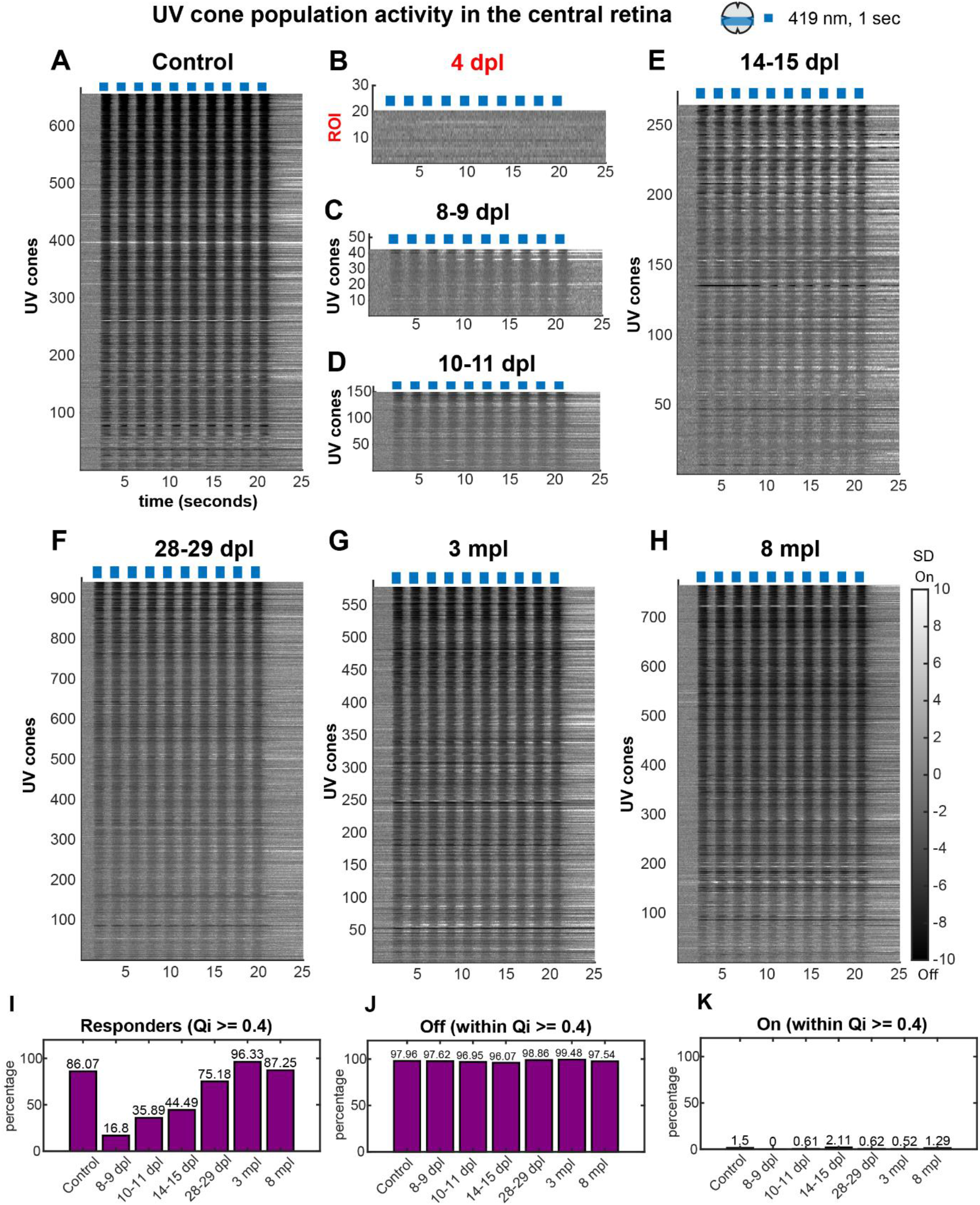
A-H: Progressive functional recovery of UV cone Off responses in the central retina at the population level after light lesion. (**A-H)** Heatmaps representing UV cone responses to blue (419 nm) light at the population level in Z scores (SD), in control and regenerating retinas. **(A)** Control, **(B)** 4 dpl, **(C)** 8-9 dpl, **(D)** 10-11 dpl, **(E)** 14-15 dpl, **(F)** 28-29 dpl, **(G)** 3 mpl, **(H)** 8 mpl. Darker shades indicate a drop in calcium level relative to baseline, denoting a cone’s intrinsic light response (Off), while lighter shades indicate a rise in calcium level, and reveal sign-inverted inputs from the outer retinal network (On). Qualitative examination of the heatmaps of UV cone-blue light responses in the control group reveals lots of dark responses indicative of a reduction in calcium levels relative to the baseline, signifying Off-responses (A); notably, UV cone responses from the 8-9 dpl to 14-15 dpl reveal attenuated Off responses (C-E). Finally, the responses return to control-like dark shades from 28-29 dpl until 8 mpl (F-H). Intriguingly, the much lighter tones indicating elevated calcium levels above the baseline or On-responses are negligible throughout the time course, validating the recovery of spectrally accurate “Off” responses to short-wavelength blue light in regenerating UV cones. At 4 dpl, the effect of diffuse light lesion is still evident, with loss of cones and function; 4 dpl traces do not represent UV cones; rather a large ROI encompassing the entire scan field, including the debris, as shown in Fig. S1D; 4 dpl. Sample size (n) Control-656, 8-9 dpl-42, 10-11 dpl-150, 14-15 dpl-264, 28-29 dpl-941, 3 mpl-578, 8 mpl-765 (n = UV cones); from N=10, 3, 6, 7, 6, 3, and 3 retinae at respective time points; 4 dpl: 20 ROIs from 3 retinae. UV cone traces indicate UV cones which passed cutoff QI >= 0.4 sorted for QI (most reliable responses on top). Only the UV cone response traces where all 10 stimulus triggers were detected during pre-processing are shown. Blue bars are stimulus presentations with blue (419 nm) light for 1 second. Grayscale bars are in Z scores (SD). **(I-K) Percentage of UV cone responders, Off and On responses to blue light recovered during regeneration | (A)** Percentage of UV cones satisfying QI >= 0.4 cutoff out of total UV cones; called UV cone responders; that reliably responded to stimulus. **(B)** Percentage of UV cone Off responses within UV cone responder population from A. **(C)** Percentage of UV cone On responses within the UV cone responder population from A. Sample size: Total UV cones: Control-854, 8-9 dpl-250, 10-11 dpl-457, 14-15 dpl-744, 28-29 dpl-1281, 3 mpl-600, 8 mpl-886; Responding UV cones: Control-735, 8-9 dpl-42, 10-11 dpl-164, 14-15 dpl-331, 28-29 dpl-963, 3 mpl-578, 8 mpl-773; Off responses: Control-720, 8-9 dpl-41, 10-11 dpl-159, 14-15 dpl-318, 28-29 dpl-952, 3 mpl-575, 8 mpl-754; On responses: Control-11, 8-9 dpl-0, 10-11 dpl-1, 14-15 dpl-7, 28-29 dpl-6, 3mpl-3, 8 mpl-10; n = UV cones. See also Fig. 3.

**Figure S4:**
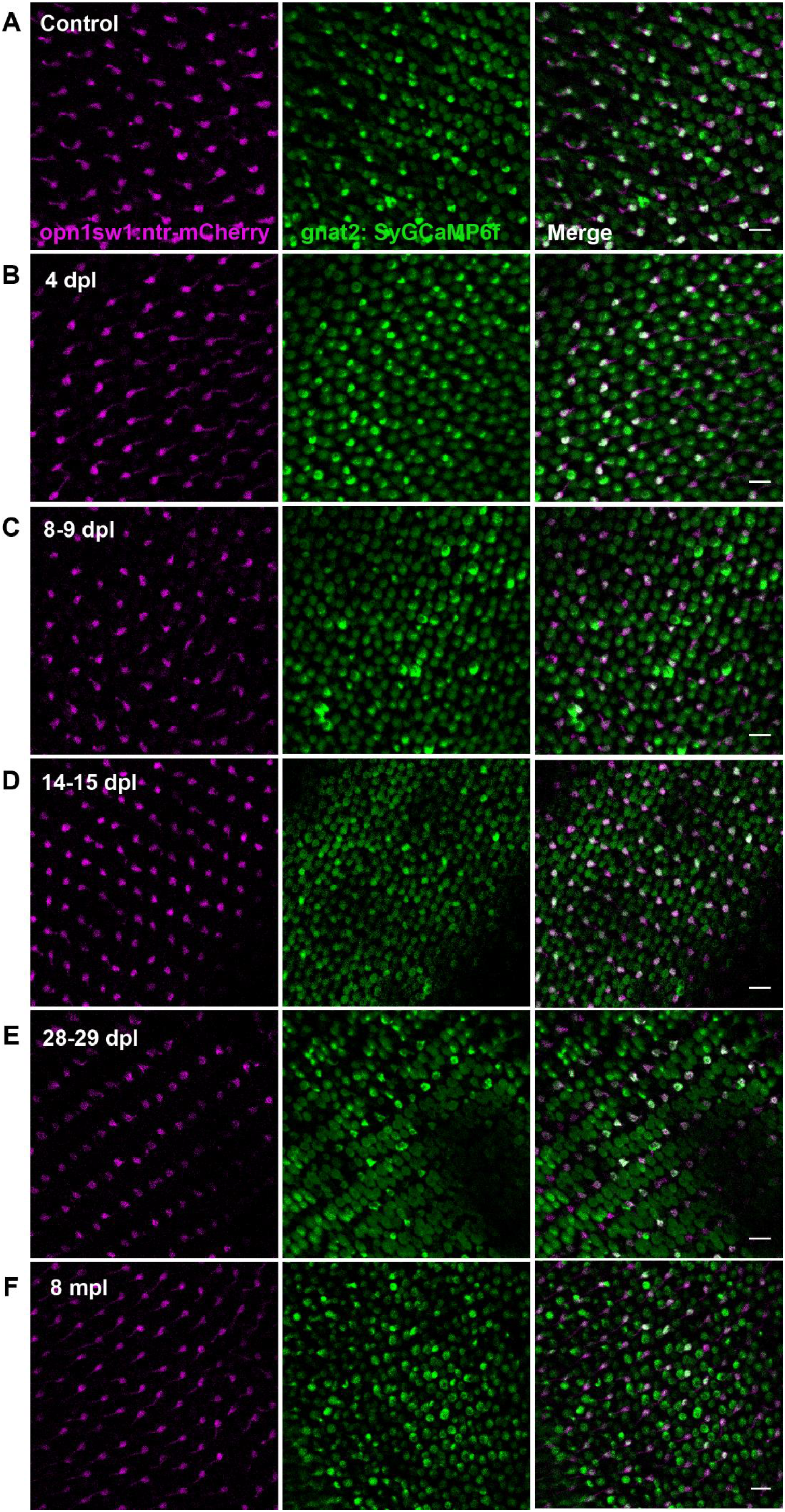
Cone photoreceptors in the ventral retina remain morphologically unlesioned after diffuse light lesion: exemplary morphological scans shown. Cones in the ventral retina in the double transgenic *Tg(gnat2:SyGCaMP6f); Tg(opn1sw1:nfsb-mCherry)* reporter after diffuse light lesion remain unlesioned. **(A)** Control**, (B)** 4 dpl, **(C)** 8-9 dpl, **(D)** 14-15 dpl, **(E)** 28-29 dpl, **(F)** 8 mpl. Similar to controls, 4 dpl ventral retina is intact with UV, blue, green and red cones, characterized by the intact cone row mosaic arrangement at all examined time points 10-11 dpl (not shown), 14-15 dpl (shown), 28-29 dpl (shown), 3 mpl (not shown) and 8 mpl (shown). Green: SyGCaMP6f, magenta: UV cone mCherry. Scale bar = 10 micrometre.

**Figure S5:**
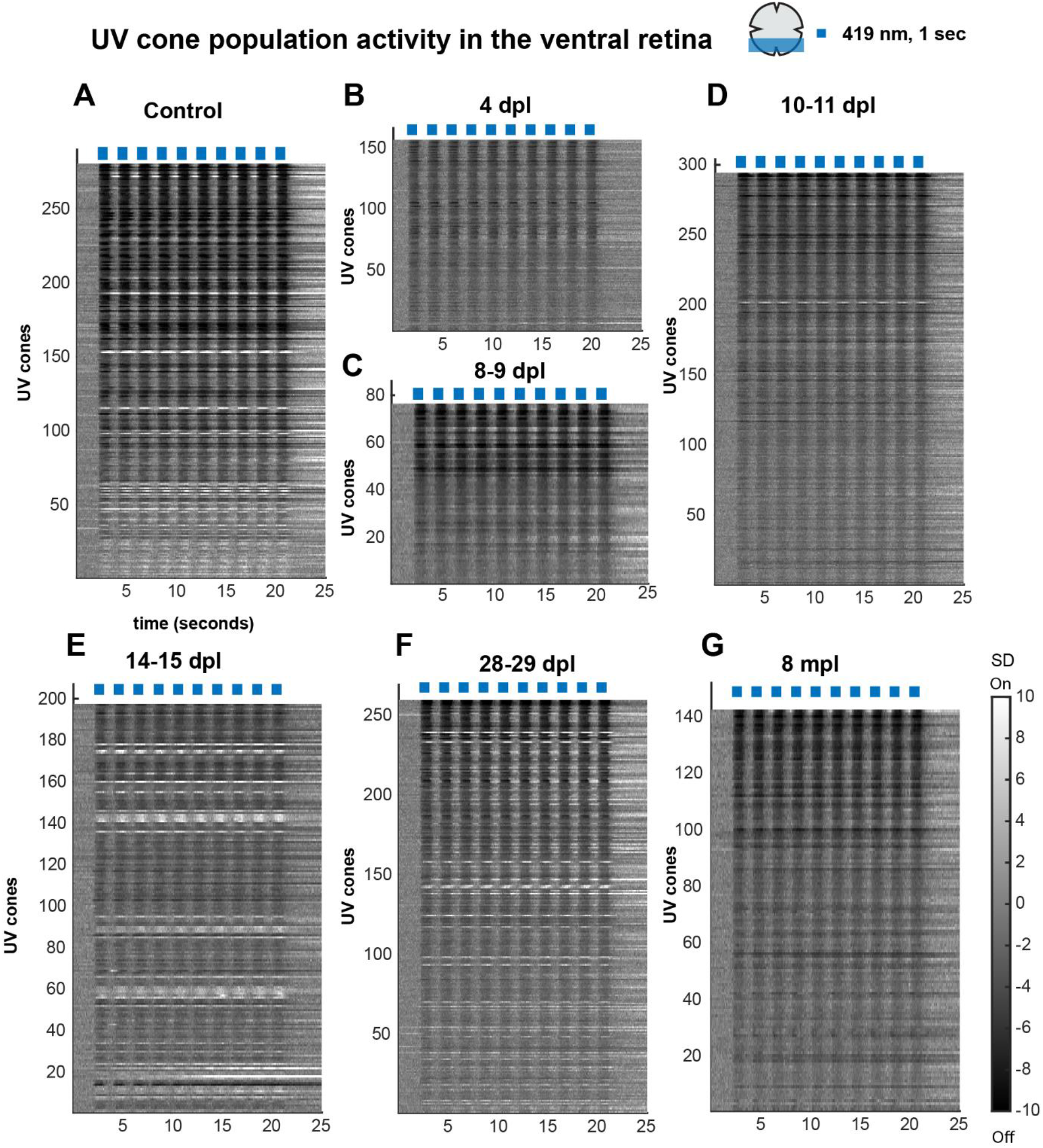
UV cone Off responses in the ventral retina is not affected by light lesion at 4 dpl and is an excellent internal control during regeneration. **A-G:** Heatmaps representing UV cone responses to blue (419 nm) light at the population level in the ventral unlesioned zone in Z scores (SD) in control and regenerating retinas. (**A)** Control, **(B)** 4 dpl, **(C)** 8-9 dpl, (**D)** 10-11 dpl, (**E)** 14-15 dpl, **(F)** 28-29 dpl, (**G)** 8 mpl. Sample size (n) Control-280, 4 dpl-156, 8-9 dpl-76, 10-11 dpl-294, 14-15 dpl-197, 28-29 dpl-259, 8 mpl-142 (n = UV cones); from N= 4, 3, 3, 5, 5, 5, and 3 retinae at respective time points. UV cone traces indicate UV cones which passed cutoff QI >= 0.4 sorted for QI (most reliable responses on top). Only the UV cone response traces where all 10 stimulus triggers were detected during pre-processing are included. Blue bars are stimulus presentations with blue (419 nm) light for 1 second. Grayscale bars are in Z scores (SD). Darker shades indicate a drop in calcium level relative to baseline, denoting a cone’s intrinsic light response (Off), while lighter shades indicate a rise in calcium level, and reveal sign-inverted inputs from the outer retinal network (On).

**Figure S6:**
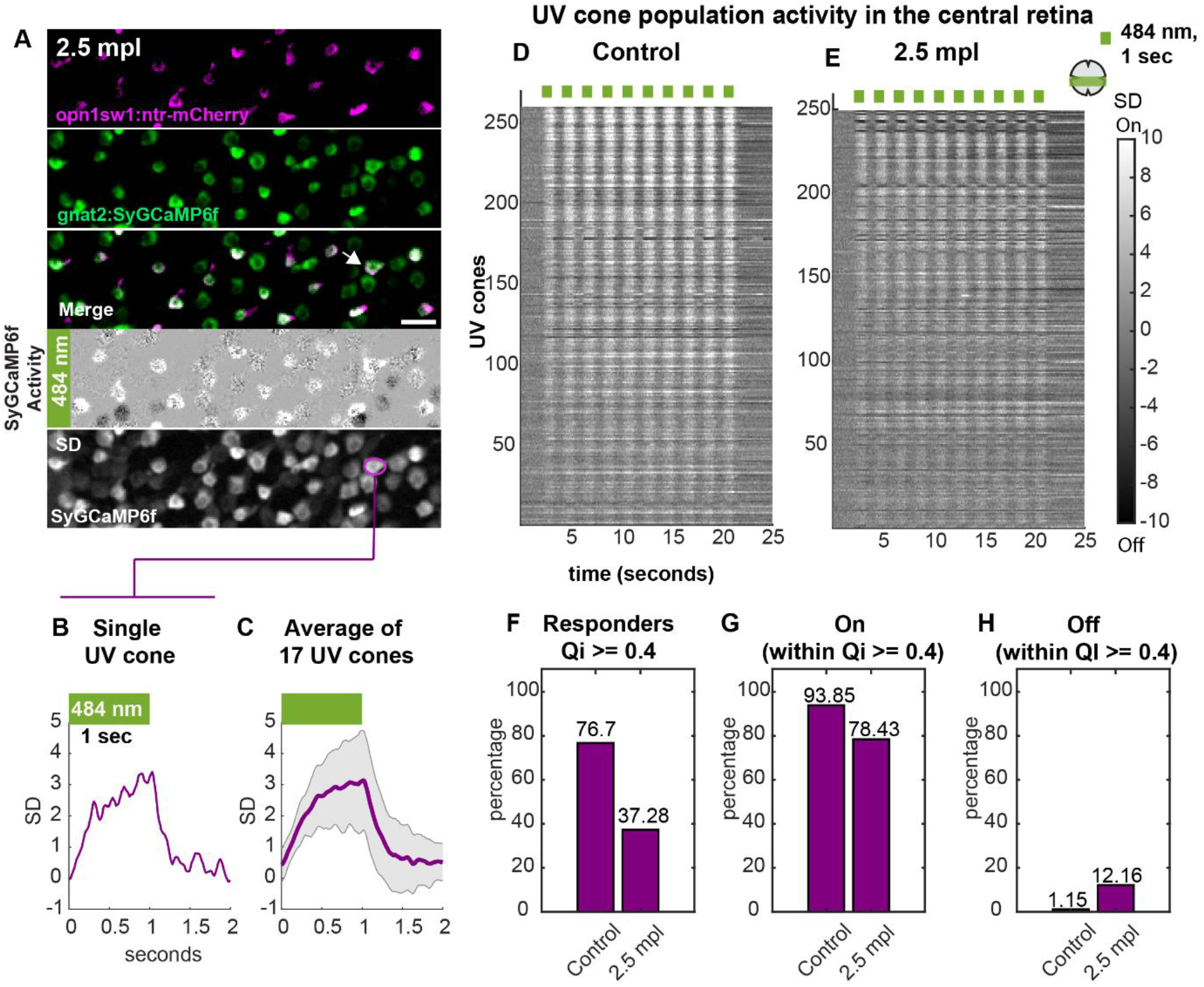
Functional recovery of On response in regenerating UV cones in the central retina after light lesion. **(A)** Example functional scan fields from the 2.5 mpl regenerated central retina of the double transgenic *Tg(gnat2:SyGCaMP6f); Tg(opn1sw1:nfsb-mCherry)*, stimulated with green (484 nm) LED, Green-SyGCaMP6f, magenta-UV cone mCherry, Scale bar = 10 micrometre. Cone activity and standard deviation (SD) projection of SyGCaMP6f from A shown; with an example UV cone (ROI) marked in magenta. Activity image: Dark terminals represent Off responses, whereas lighter terminals represent On responses; see also **Movie 11**. **(B)** Single UV cone light On response from A. **(C)** Averaged light responses from all n = 17 UV cones from A, represented as mean and standard deviation (shaded error bar). Y-axis: standard deviation (SD) and X-axis: time (seconds). **(D, E)** Heatmaps representing UV cone responses to green (484 nm) light at the population level in Z scores (SD) in control and regenerating retinas. **(D)** Control, **(E)** 2.5 mpl, Sample size (n) Control-260, 2.5 mpl-249, (n = UV cones); from N= 5, and 6 retinae, at respective time points. UV cone traces indicate UV cones which passed cutoff QI >= 0.4 sorted for QI (most reliable responses on top). Only the UV cone response traces where all 10 stimulus triggers were detected during pre-processing are shown. Grayscale bars are in Z scores (SD). Darker shades indicate a drop in calcium level relative to baseline, denoting a cone’s intrinsic light response (Off), while lighter shades indicate a rise in calcium level, and reveal sign-inverted inputs from the outer retinal network (On). **(F)** Percentage of UV cones satisfying QI >= 0.4 cutoff out of total UV cones; called UV cone responders; that reliably responded to stimulus. **(G)** Percentage of UV cone On responses within UV cone responder population from D. **(H)** Percentage of UV cone Off responses within UV cone responder population from D. Sample size: Total UV cones: Control-339, 2.5 mpl-684; Responding UV cones: Control-260, 2.5 mpl-255; On responses: Control-244, 2.5 mpl-200; Off responses: Control-3, 2.5 mpl-31. Green bar: green (484 nm) stimulus for 1 sec. See also Fig. 4.

**Figure S7:**
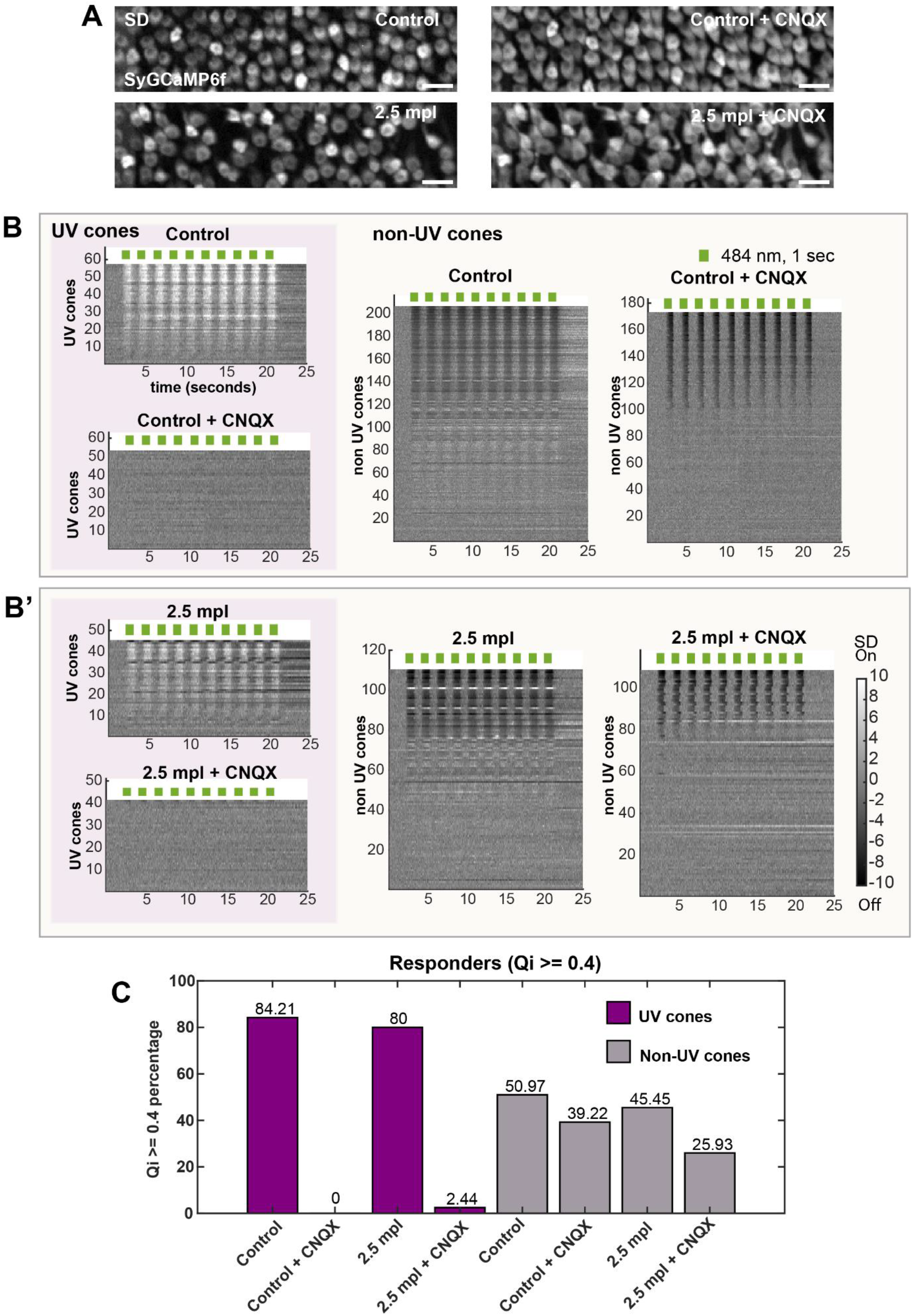
CNQX treatment in unique scan fields from the same retina in the control and 2.5 mpl regenerated retina successfully eliminates all On responses and preserves Off responses. **(A)** Standard deviation (SD) projection of SyGCaMP6f from representative scans shown in Fig. 4D and 4F. SD projection is used for placing cone ROIs. Grey: SyGCaMP6f signal. Scale bar = 10 micrometre. **(B)** Heatmap of response traces of UV cones and non-UV cones (in Z scores) from 4 unique scan fields (N= 4 retinae) fields in the control group to green light flashes (green bars on top) before and after HC block, as indicated. Sample size (n): Control-57, control + HC block-53 (n = UV cones); Sample size (n): Control-206, control + HC block-173, (n = non-UV cones) **(B’)**-Heatmap of response traces of UV cones and non UV cones (in Z scores) from 3 unique scan fields (N= 3 retinae) in 2.5 mpl group to green light flashes (green bars on top) before and after HC block, as indicated. Sample size (n): 2.5 mpl-45, 2.5 mpl + HC block-41, (n = UV cones); Sample size (n): 2.5 mpl-110, 2.5 mpl + HC block-108, (n = non-UV cones). Grayscale bars are in Z scores (SD). Darker shades indicate calcium drop relative to baseline, denoting a cone’s intrinsic light response (Off), while lighter shades indicate a rise in calcium and reveal sign-inverted inputs from the outer retinal network (On). In all heatmaps, cone response traces are sorted for increasing QI (most reliable responses on top) and all responses passed cutoff QI >= 0, where 10 stimulus triggers were detected are shown. **(C)** Percentage of UV cone responders before and after CNQX treatment in the control and 2.5 mpl groups. Sample size: Total UV cones: Control-57, Control + CNQX-53; Responding UV cones: Control-48, Control + CNQX-0; Total UV cones: 2.5 mpl-45, 2.5 mpl + CNQX-41; Responding UV cones: 2.5 mpl-36, 2.5 mpl + CNQX-1; Total non-UV cones: Control-206, Control + CNQX-204; Responding non-UV cones: Control-105, Control + CNQX-80; Total non-UV cones: 2.5 mpl-110, 2.5 mpl + CNQX-108; Responding non-UV cones: 2.5 mpl-50, 2.5 mpl + CNQX-28. See also Fig. 4.

**Figure S8:**
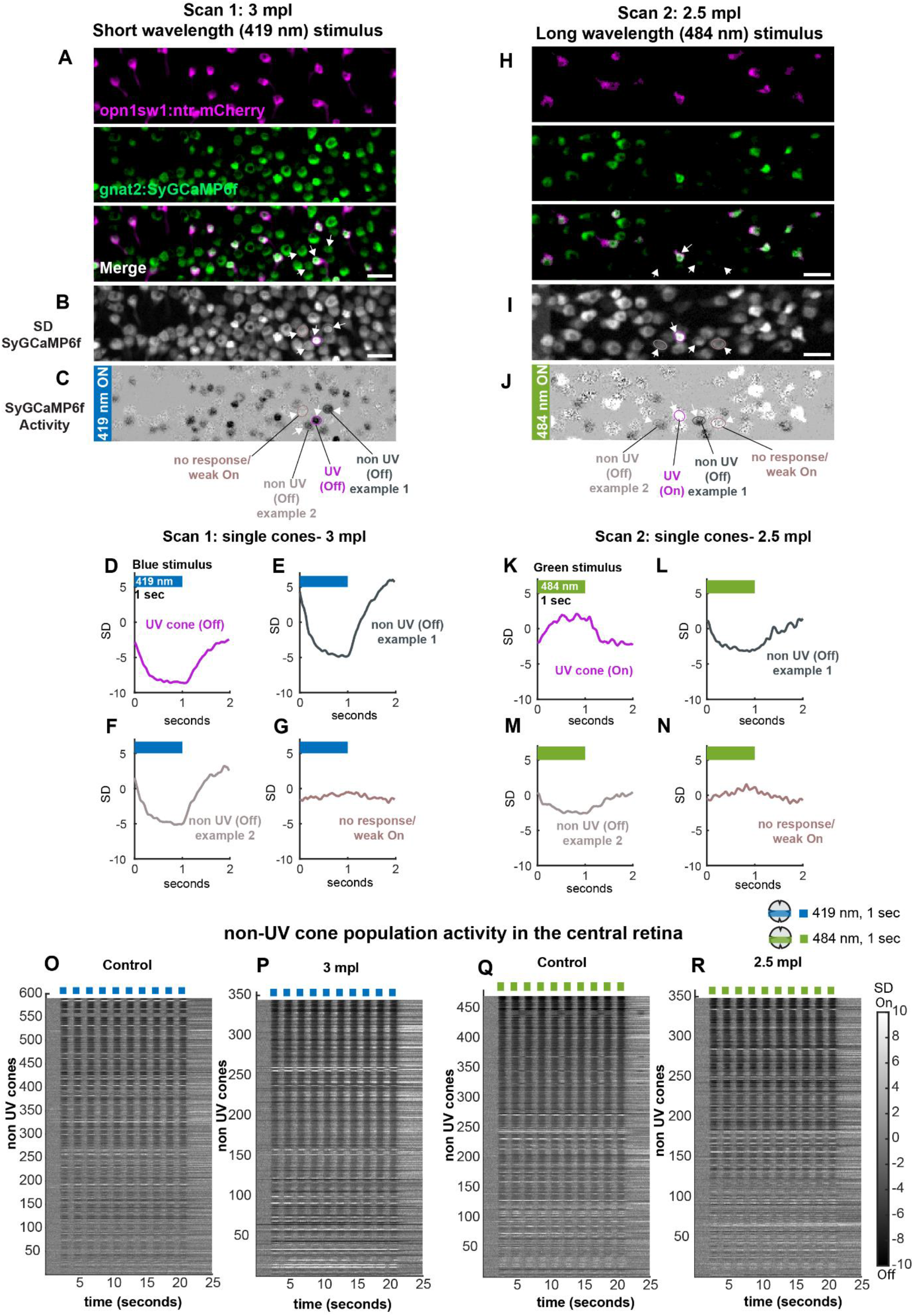
Non-UV cone responses to short wavelength (blue, 419 nm) and long wavelength (green, 484 nm) light stimulation are recovered in the regenerating retina. **(A)** Example functional scan fields from the central retina of double-transgenic *Tg(gnat2:SyGCaMP6f); Tg(opn1sw1:nfsb-mCherry)* retinae at 3 mpl, stimulated with blue light. **(B)** Standard deviation (SD) projection of SyGCaMP6f from A with quartet of ROIs, one UV cone and three non-UV cone ROI marked (also in A and C). **(C)** Activity image of A, showing cone SyGCaMP6f responses to blue light. Dark terminals represent Off responses, whereas lighter terminals represent On responses. **(D-G)** Example cone responses from ROIs in B (also in A and C): UV cone Off response, non UV cone Off response example 1, non UV cone Off response example 2, and non UV cone (non-responsive/ weakly On), respectively. **(G)** Example functional scan fields from the central retina of double-transgenic *Tg(gnat2:SyGCaMP6f); Tg(opn1sw1:nfsb-mCherry)* retinae at 2.5 mpl, stimulated with green light. **(I)** Standard deviation (SD) projection of SyGCaMP6f from H with quartet of ROIs, one UV cone and three non-UV cone ROI marked (also in H and J). **(J)** Activity image of H, showing cone SyGCaMP6f responses to green light. Dark terminals represent Off responses, whereas lighter terminals represent On responses. **(K-N)** Example cone responses from ROIs in I (also in H and J): UV cone On response, non UV cone Off response example 1, non UV cone Off response example 2, and non UV cone (non-responsive/ weakly On), respectively. A and H are separate scans from different retinas. Green-SyGCaMP6f, magenta-UV cone mCherry; Scale bar = 10 micrometre. **(O)** Control, **(P)** 3 mpl, **(Q)** Control, **(R)** 2.5 mpl. In some control scans we also see an On response artifact from blue cones to blue light; this was reasoned as a consequence of calcium responses from an imaing plane closer to the soma. Sample size(n): blue stimulus: Control-590, 3 mpl-344, (n = non UV cones); from N= 5, and 3 retinae; green stimulus: Control-470, 2.5 mpl-348, (n = non UV cones); from N= 5, and 6 retinae. Non-UV cone traces indicate non-UV cones which passed cutoff QI >= 0.4 sorted for QI (most reliable responses on top). Only the non UV cone response traces where all 10 stimulus triggers were detected during pre-processing are included. Blue and green bars are stimulus presentations with blue (419 nm) and green (484 nm) light for 1 second respectively. Grayscale bars are in Z scores (SD). Darker shades indicate a drop in calcium level relative to baseline, denoting a cone’s intrinsic light response (Off), while lighter shades indicate a rise in calcium level, and reveal sign-inverted inputs from the outer retinal network (On). See also Fig. 1, 2 and 4.

### SUMMARY STATISTICS

**Table 1:**
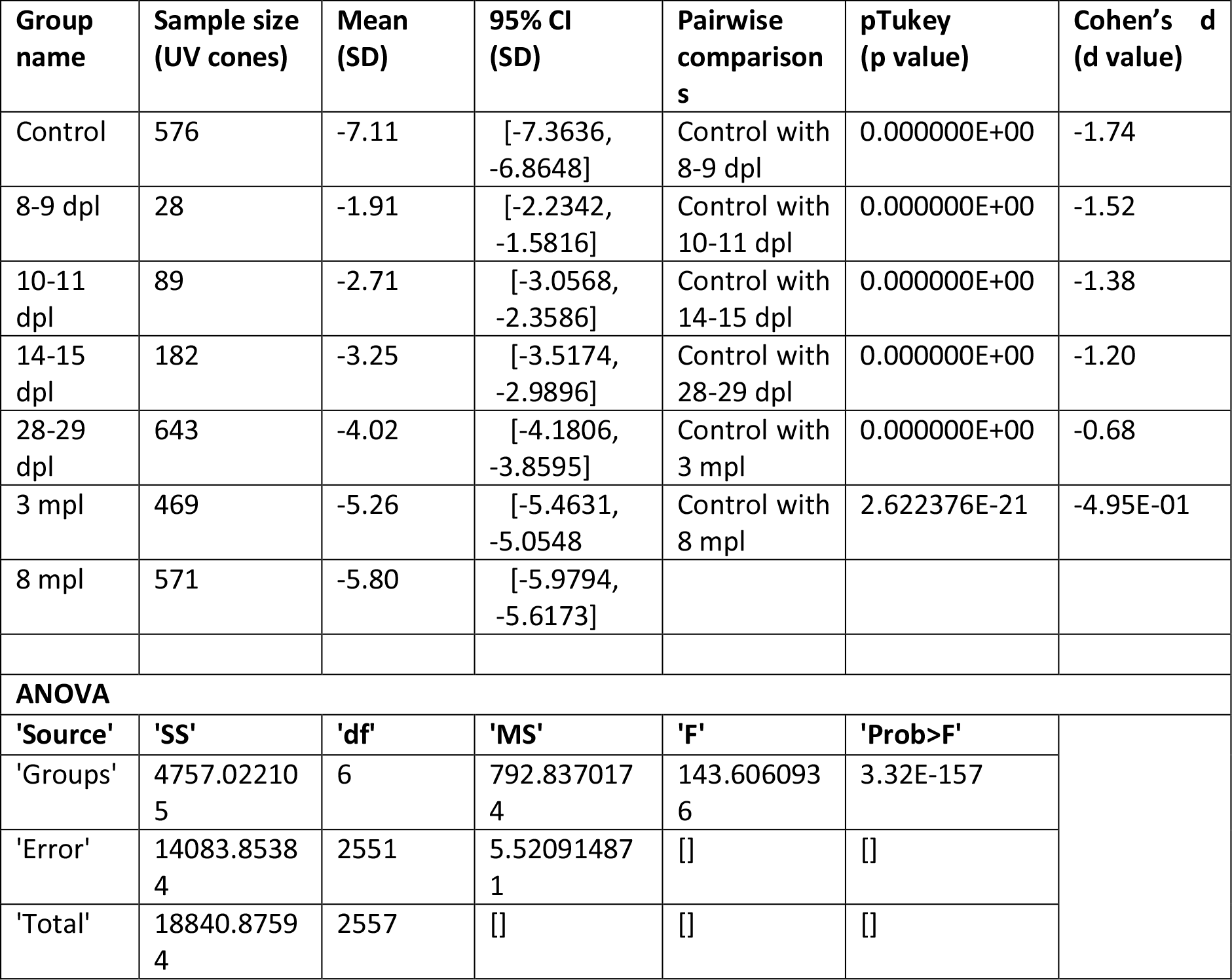
Summary statistics of UV cone Off response amplitude during regeneration: related to Figure 3B. Time course of development of UV cone Off response amplitude during regeneration time course. One-way analysis of variance or ANOVA followed by Tukey Kramer post-hoc analysis for unequal sample sizes was used whenever multiple pairwise comparisons of independent control and experimental groups were done. Summary of groups, sample size, mean, 95% confidence interval (95% CI) of mean, pairwise comparisons, p values corresponding to significance of multiple comparisons of control group to 6 regenerated time points, and effect size of the difference for each comparison in terms of Cohen’s d. ANOVA results prior to Tukey Kramer post-hoc analysis for unequal sample sizes summarized. ‘SS’-sum of squares, ‘df’ - degrees of freedom, ‘MS’-mean squared error, ‘F’-*F* statistic. The ‘Prob’-is the *p*-value or probability that the test statistic can take a value greater than the value of the computed test statistic. Cohen’s d (d value) can be read as Very small-0.01, Small-0.20, Medium-0.50, Large-0.80, Very large-1.20, Huge-2.0. Sign indicates direction of the effect.

**Table 2:**
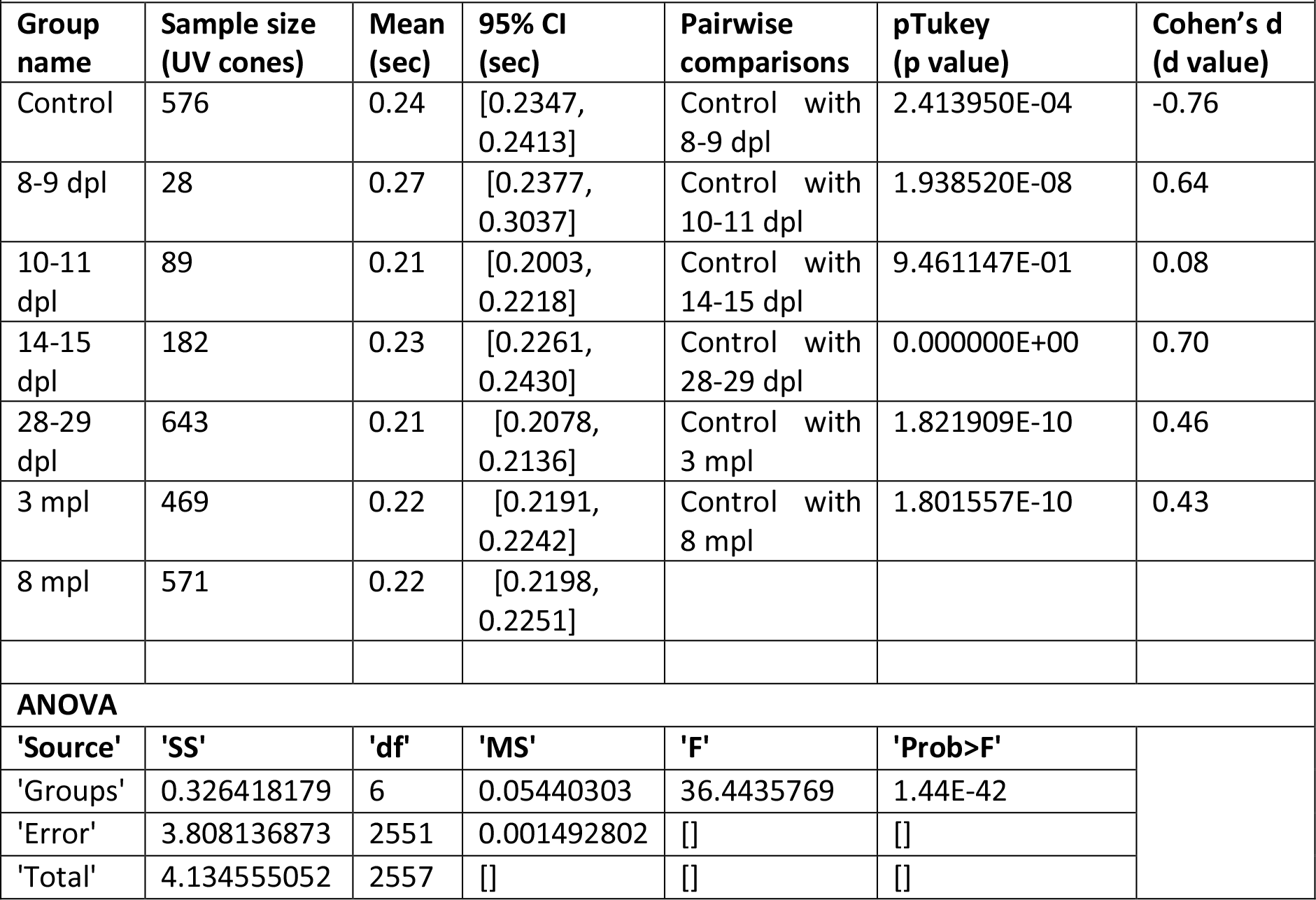
Summary statistics of UV cone Off response Tau decay during regeneration: related to Figure 3C. Time course of development of UV cone Off response Tau decay during regeneration time course. One-way analysis of variance or ANOVA followed by Tukey Kramer post-hoc analysis for unequal sample sizes was used whenever multiple pairwise comparisons of independent control and experimental groups were done. Summary of groups, sample size, mean, 95% confidence interval (95% CI) of mean, pairwise comparisons, p values corresponding to significance of multiple comparisons of control group to 6 regenerated time points, and effect size of the difference for each comparison in terms of Cohen’s d. ANOVA results prior to Tukey Kramer post-hoc analysis for unequal sample sizes summarized. ‘SS’-sum of squares, ‘df’ - degrees of freedom, ‘MS’-mean squared error, ‘F’-*F* statistic. The ‘Prob’-is the *p*-value or probability that the test statistic can take a value greater than the value of the computed test statistic. Cohen’s d (d value) can be read as Very small-0.01, Small-0.20, Medium-0.50, Large-0.80, Very large-1.20, Huge-2.0. Sign indicates direction of the effect.

**Table 3:**
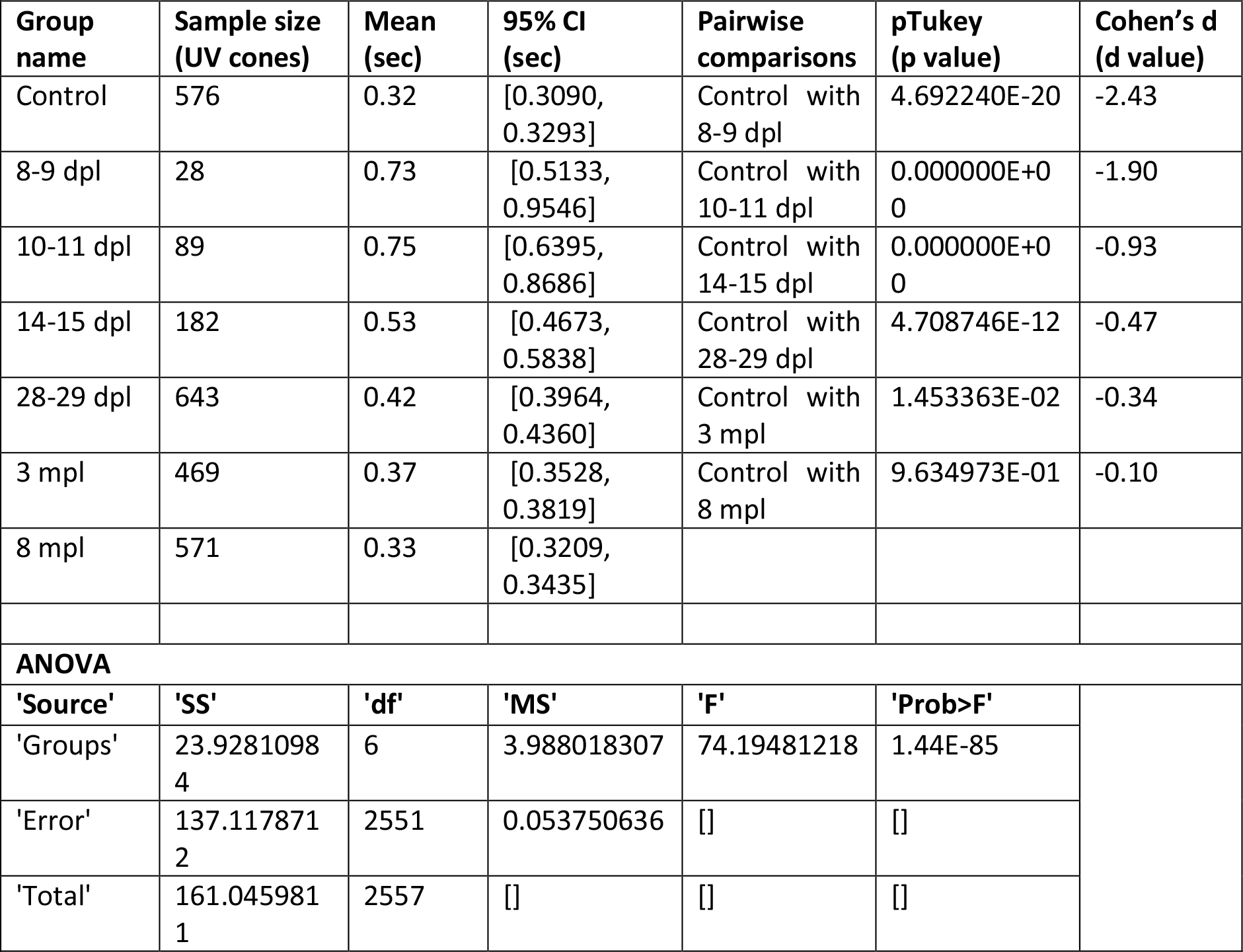
Summary statistics of UV cone Off response Tau recovery during regeneration: related to Figure 3D. Time course of development of UV cone Off response Tau recovery during regeneration time course. One-way analysis of variance or ANOVA followed by Tukey Kramer post-hoc analysis for unequal sample sizes was used whenever multiple pairwise comparisons of independent control and experimental groups were done. Summary of groups, sample size, mean, 95% confidence interval (95% CI) of mean, pairwise comparisons, p values corresponding to significance of multiple comparisons of control group to 6 regenerated time points, and effect size of the difference for each comparison in terms of Cohen’s d. ANOVA results prior to Tukey Kramer post-hoc analysis for unequal sample sizes summarized. ‘SS’-sum of squares, ‘df’ - degrees of freedom, ‘MS’-mean squared error, ‘F’-*F* statistic. The ‘Prob’-is the *p*-value or probability that the test statistic can take a value greater than the value of the computed test statistic. Cohen’s d (d value) can be read as Very small-0.01, Small-0.20, Medium-0.50, Large-0.80, Very large-1.20, Huge-2.0. Sign indicates direction of the effect.

**Table 4:**
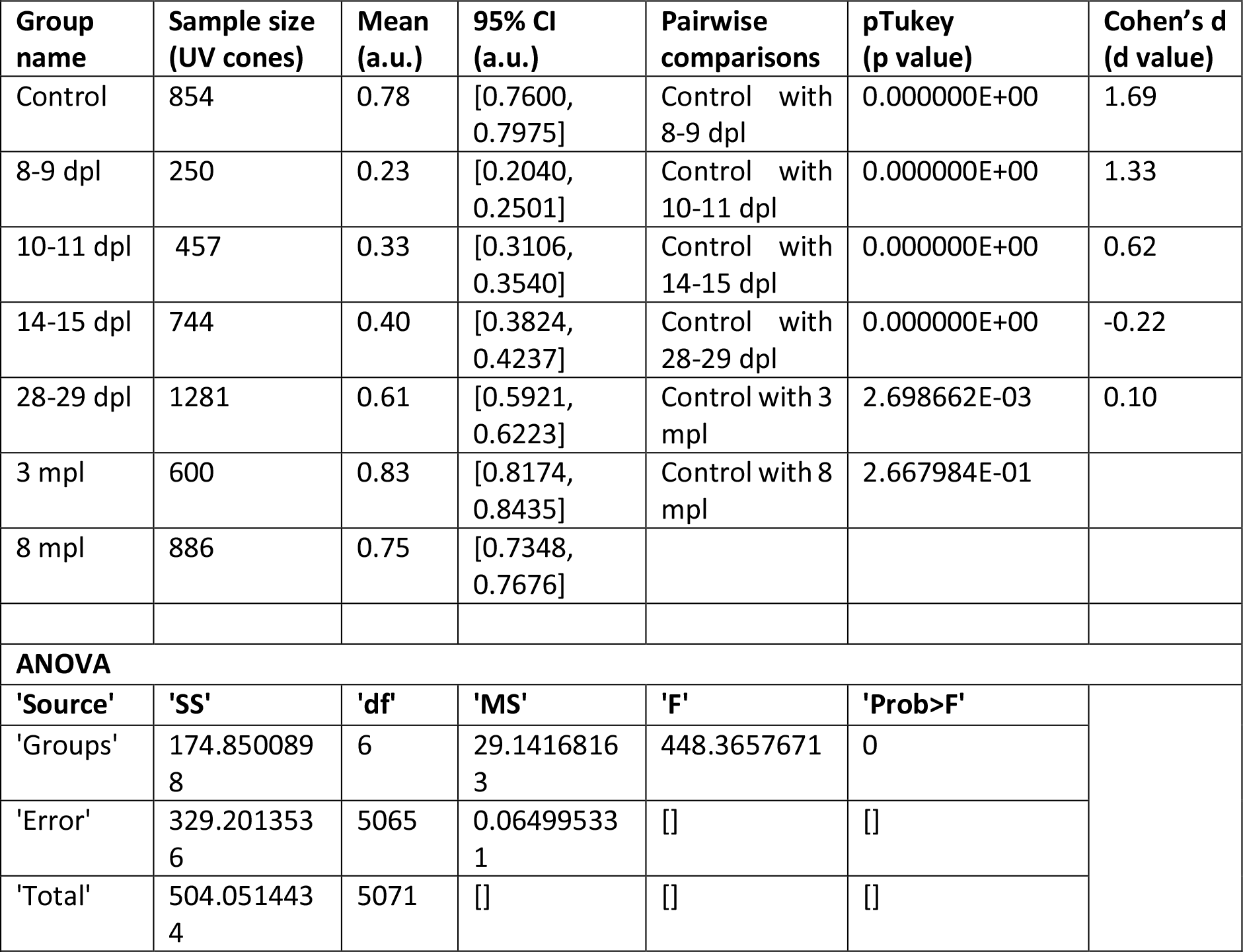
Summary statistics of UV cone response quality during regeneration: related to Figure 3E. Time course of development of Quality index (QI) of UV cones during regeneration time course. One-way analysis of variance or ANOVA followed by Tukey Kramer post-hoc analysis for unequal sample sizes was used whenever multiple pairwise comparisons of independent control and experimental groups were done. Summary of groups, sample size, mean, 95% confidence interval (95% CI) of mean, pairwise comparisons, p values corresponding to significance of multiple comparisons of control group to 6 regenerated time points, and effect size of the difference for each comparison in terms of Cohen’s d. ANOVA results prior to Tukey Kramer post-hoc analysis for unequal sample sizes summarized. ‘SS’-sum of squares, ‘df’ - degrees of freedom, ‘MS’-mean squared error, ‘F’-*F* statistic. The ‘Prob’-is the *p*-value or probability that the test statistic can take a value greater than the value of the computed test statistic. Cohen’s d (d value) can be read as Very small-0.01, Small-0.20, Medium-0.50, Large-0.80, Very large-1.20, Huge-2.0. Sign indicates direction of the effect.

**Table 5:**
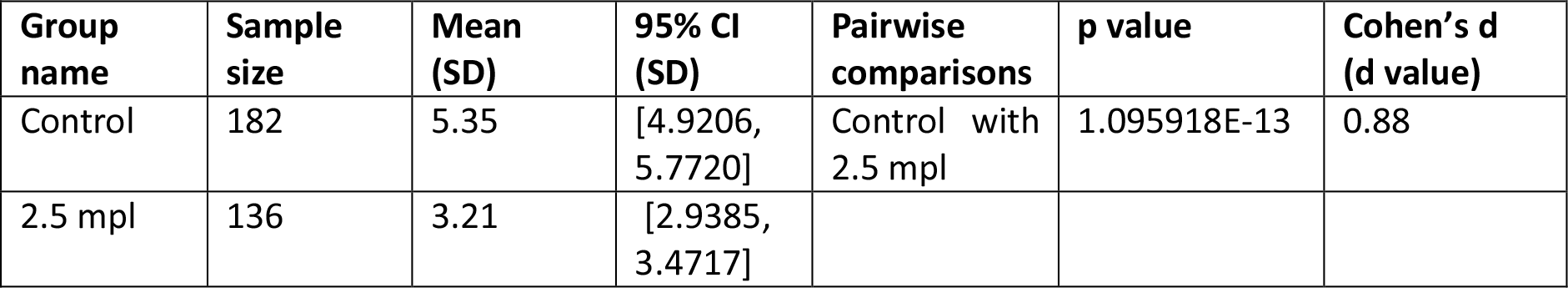
Summary statistics of UV cone On response amplitude during regeneration: related to Figure 4B. Time course of development of UV cone On response amplitude in regenerating UV On cones. Two sample t-test for unequal sample sizes and independent groups was used to compare control and experimental group. Summary of groups, sample sizes, mean, 95% confidence interval (95% CI), pairwise comparison, p values, and effect size of the difference in terms of Cohen’s d value are indicated. Cohen’s d (d value) can be read as Very small-0.01, Small-0.20, Medium-0.50, Large-0.80, Very large-1.20, Huge-2.0. Sign indicates direction of the effect.

**Table 6:**
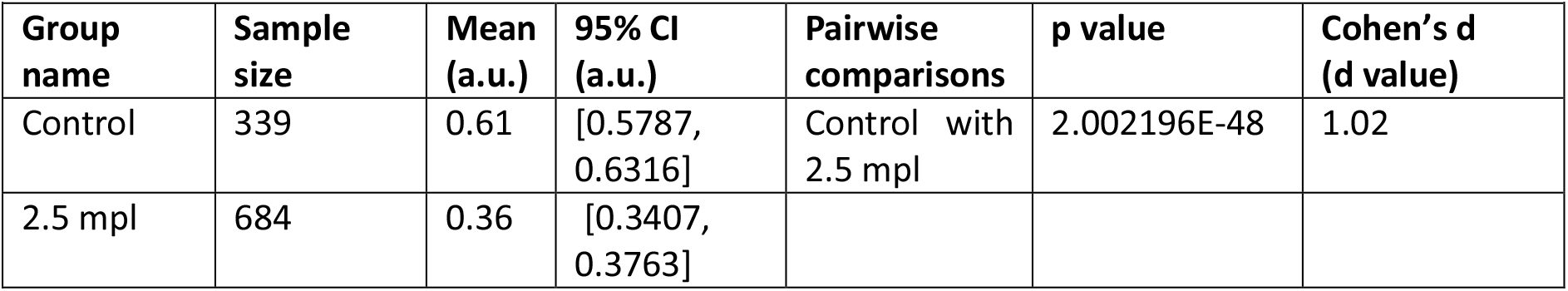
Summary statistics of UV cone response quality during regeneration: related to Figure 4C. Time course of development Quality Index (QI) in regenerating UV cones. Two sample t-test for unequal sample sizes and independent groups was used to compare control and experimental group. Summary of groups, sample sizes, mean, 95% confidence interval (95% CI), pairwise comparison, p values, and effect size of the difference in terms of Cohen’s d value are indicated. Cohen’s d (d value) can be read as Very small-0.01, Small-0.20, Medium-0.50, Large-0.80, Very large-1.20, Huge-2.0. Sign indicates direction of the effect.

