## Supplemental data and methods for "Restoration of Cone Circuit Functionality in the Regenerating Adult Zebrafish Retina"

### SUPPLEMENTARY FIGURES

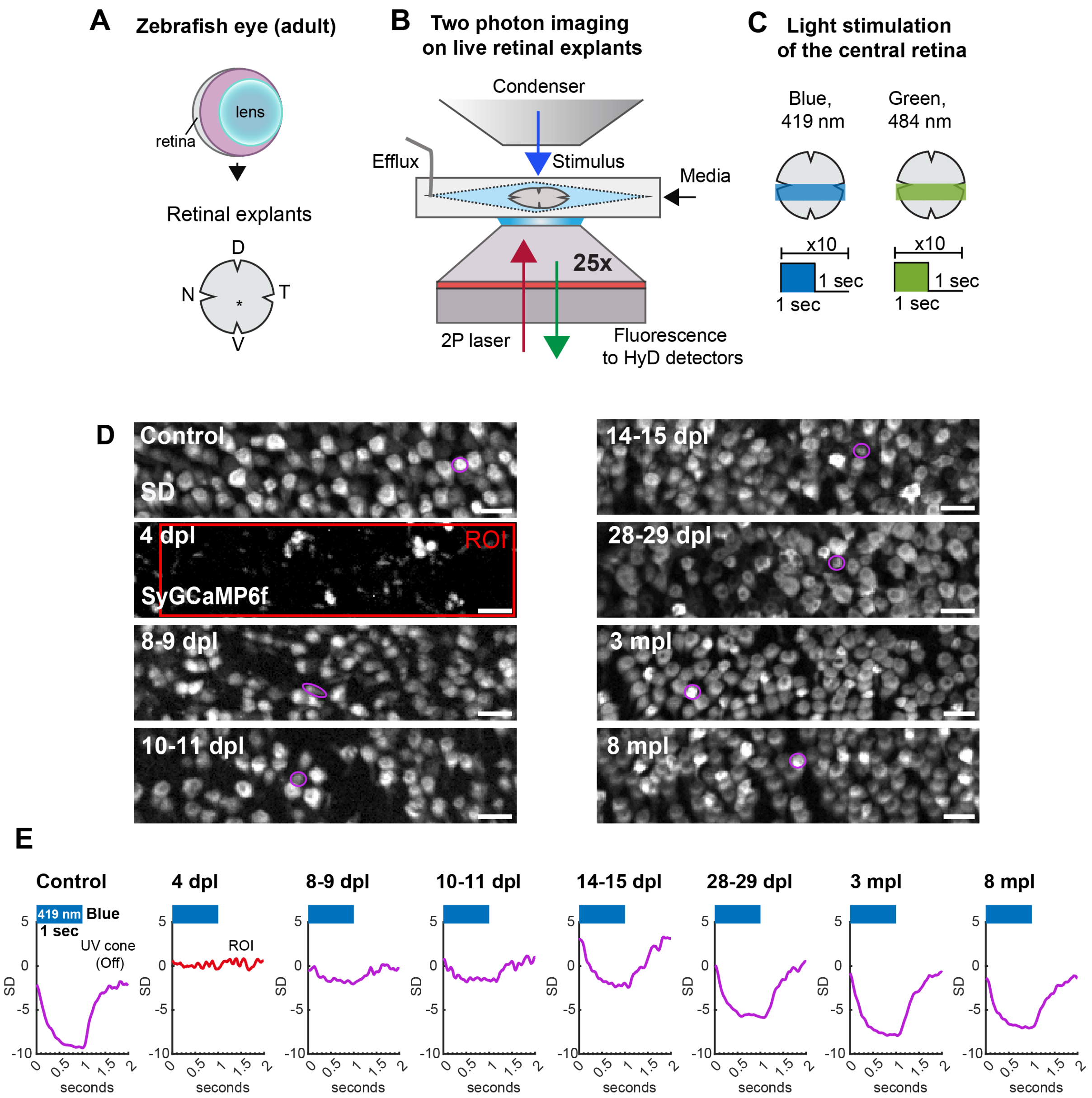

**Figure S1: (A-C) 2P calcium imaging with light stimulation in adult zebrafish retinal explants | (A)** Zebrafish adult retinal whole mount preparation (explants), orientation preserved. **(B)** Retinal explant live-imaging with continuous oxygenated media perfusion under an inverse 2P microscope. 2P excitation (red arrow), light stimulus through condenser (blue arrow), emitted fluorescence (green arrow). **(C)** Light stimulation protocol: LED ON for 1 second and interval for 1 sec, repeated 10 times for each scan field in the central retina. nasal (N), temporal (T), dorsal (D), and ventral (V). **(D-E) UV cone functional recovery in the central retina at the single cone level | (D)** Standard deviation (SD) projection of SyGCaMP6f imaging stack from representative scans shown in Fig. 2D. SD projection is used for placing cone ROIs; here single UV cone ROIs are shown encircled in magenta. At 4 dpl, cones are still ablated in the central retina, with only debris remaining; large rectangular ROI shown in red. Note that non-UV cones ROIs can also be analyzed in this way. **(E)** Light responses of single UV cones from A. Blue bar indicates blue LED stimulus presentation for 1 sec. Control, and 8-9 dpl - 8 mpl shows UV cone Off responses, whereas 4 dpl shows loss of function or random noise. Y- axis: standard deviation (SD) and X- axis: time (seconds). Grey: SyGCaMP6f signal. Scale bar = 10 micrometre. See also Fig. 1-2.

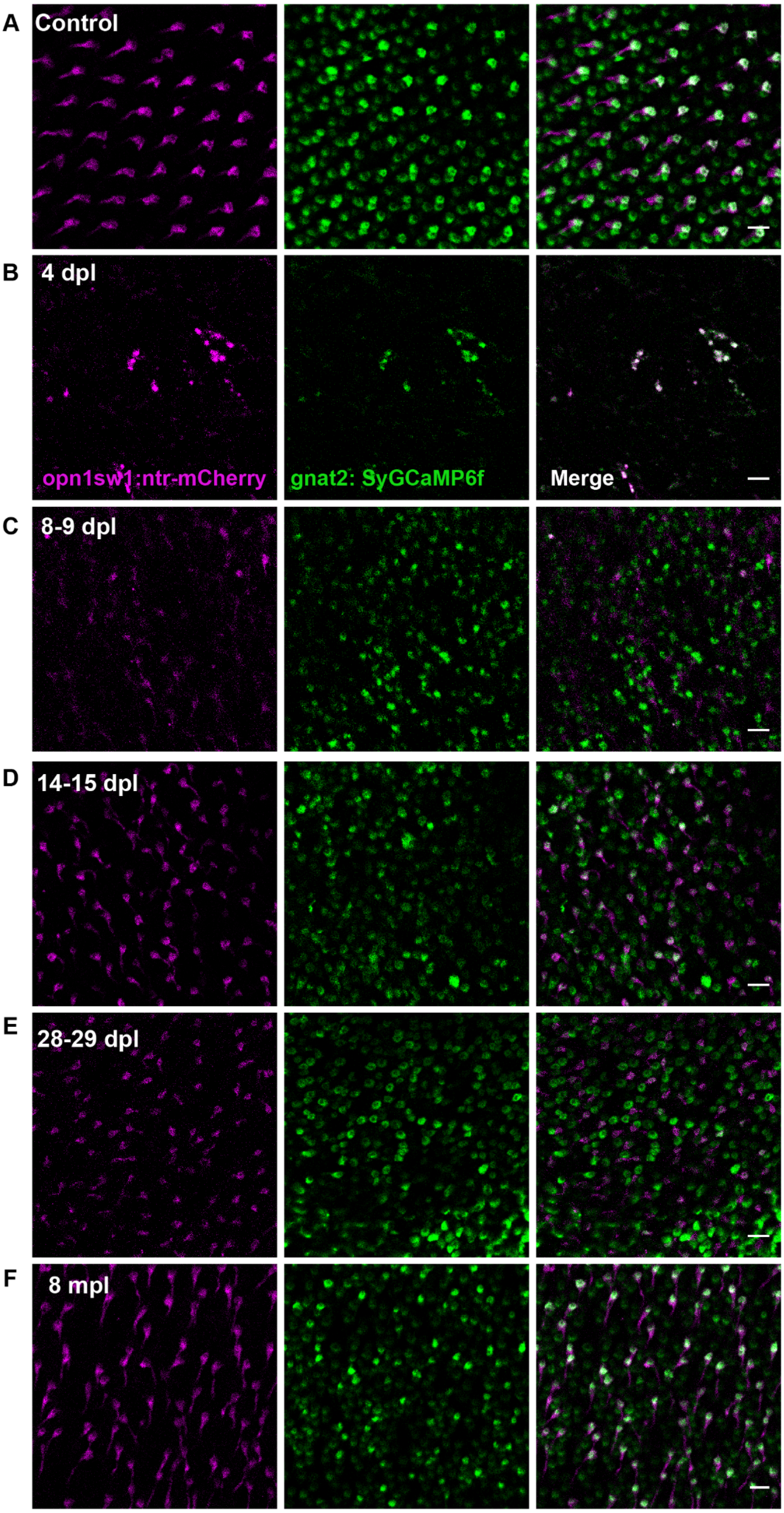
A
Control
B
4 dpl
opn1sw1:ntr-mCherry
gnat2: SyGCaMP6f
Merge
C
8-9 dpl
D
14-15 dpl
E
28-29 dpl
F
8 mpl

**Figure S2: Cone photoreceptors in the central retina are completely ablated after light lesion, but continue to regenerate from 8 dpl to 8 mpl- exemplary morphological scans shown |** Regenerating cones in the central retina in double transgenic *Tg(gnat2:SyGCaMP6f); Tg(opn1sw1:nfsb-mCherry)* reporter after diffuse light lesion **(A)** Control, **(B)** 4 dpl, **(C)** 8-9 dpl, **(D)** 14-15 dpl, **(E)** 28-29 dpl, **(F)** 8 mpl. Several morphological scan fields from respective time points were examined during functional recordings, and representative images are shown. Compared to controls, 4 dpl central retina is completely ablated of UV cones and the bulk of blue, green and red cones. The bulk of regenerated cones are captured in the central retina where the UV cone row mosaic is disrupted, starting at 8-9 dpl and remains prominent at all other examined time points 10-11 dpl (not shown), 14-15 dpl (shown), 28-29 dpl (shown), 3 mpl (not shown) and 8 mpl (shown). Green: SyGCaMP6f, magenta: UV cone mCherry. Scale bar = 10 micrometre.

Abraham et al

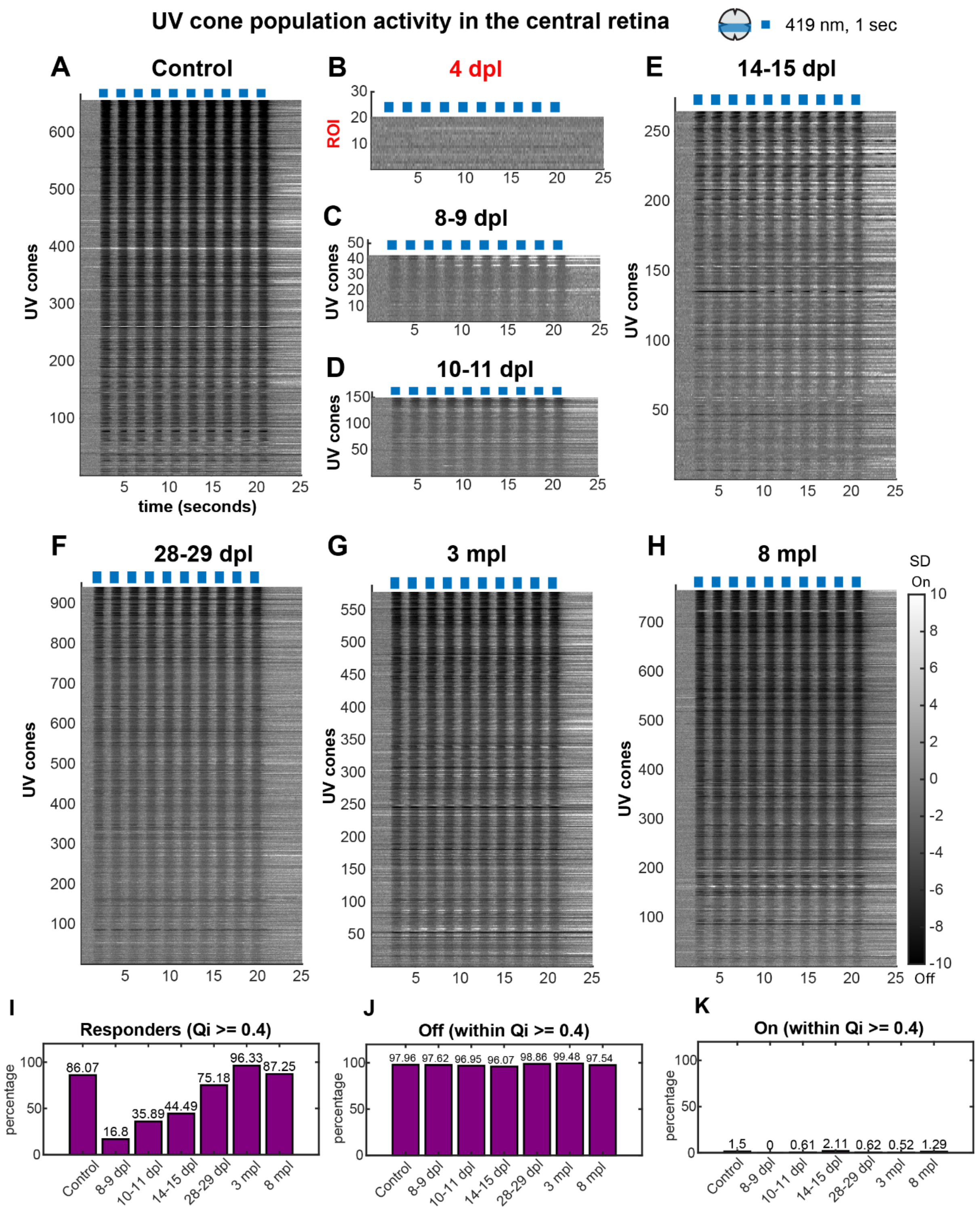

**Figure S3: A-H: Progressive functional recovery of UV cone Off responses in the central retina at the population level after light lesion |** (**A-H)** Heatmaps representing UV cone responses to blue (419 nm) light at the population level in Z scores (SD), in control and regenerating retinas. **(A)** Control, **(B)** 4 dpl, **(C)** 8-9 dpl, **(D)** 10-11 dpl, **(E)** 14-15 dpl, **(F)** 28-29 dpl, **(G)** 3 mpl, **(H)** 8 mpl. Darker shades indicate a drop in calcium level relative to baseline, denoting a cone's intrinsic light response (Off), while lighter shades indicate a rise in calcium level, and reveal sign-inverted inputs from the outer retinal network (On). Qualitative examination of the heatmaps of UV cone- blue light responses in the control group reveals lots of dark responses indicative of a reduction in calcium levels relative to the baseline, signifying Off-responses (A); notably, UV cone responses from the 8-9 dpl to 14-15 dpl reveal attenuated Off responses (C-E). Finally, the responses return to control-like dark shades from 28-29 dpl until 8 mpl (F-H). Intriguingly, the much lighter tones indicating elevated calcium levels above the baseline or On-responses are negligible throughout the time course, validating the recovery of spectrally accurate "Off" responses to short-wavelength blue light in regenerating UV cones. At 4 dpl, the effect of diffuse light lesion is still evident, with loss of cones and function; 4 dpl traces do not represent UV cones; rather a large ROI encompassing the entire scan field, including the debris, as shown in Fig. S1D; 4 dpl. Sample size (n) Control- 656, 8-9 dpl- 42, 10-11 dpl- 150, 14-15 dpl- 264, 28-29 dpl- 941, 3 mpl- 578, 8 mpl-765 (n = UV cones); from N=10, 3, 6, 7, 6, 3, and 3 retinae at respective time points; 4 dpl: 20 ROIs from 3 retinae. UV cone traces indicate UV cones which passed cutoff QI >= 0.4 sorted for QI (most reliable responses on top). Only the UV cone response traces where all 10 stimulus triggers were detected during pre-processing are shown. Blue bars are stimulus presentations with blue (419 nm) light for 1 second. Grayscale bars are in Z scores (SD). **(I-K) Percentage of UV cone responders, Off and On responses to blue light recovered during regeneration | (A)** Percentage of UV cones satisfying QI >= 0.4 cutoff out of total UV cones; called UV cone responders; that reliably responded to stimulus. **(B)** Percentage of UV cone Off responses within UV cone responder population from A. **(C)** Percentage of UV cone On responses within the UV cone responder population from A. Sample size: Total UV cones: Control- 854, 8-9 dpl- 250, 10-11 dpl-457, 14-15 dpl- 744, 28-29 dpl-1281, 3 mpl- 600, 8 mpl- 886; Responding UV cones: Control- 735, 8-9 dpl- 42, 10-11 dpl- 164, 14-15 dpl- 331, 28-29 dpl- 963, 3 mpl-578, 8 mpl- 773; Off responses: Control- 720, 8-9 dpl- 41, 10-11 dpl- 159, 14-15 dpl- 318, 28-29 dpl-952, 3 mpl- 575, 8 mpl- 754; On responses: Control- 11, 8-9 dpl- 0, 10-11 dpl- 1, 14-15 dpl- 7, 28-29 dpl- 6, 3mpl- 3, 8 mpl- 10; n = UV cones. See also Fig. 3.

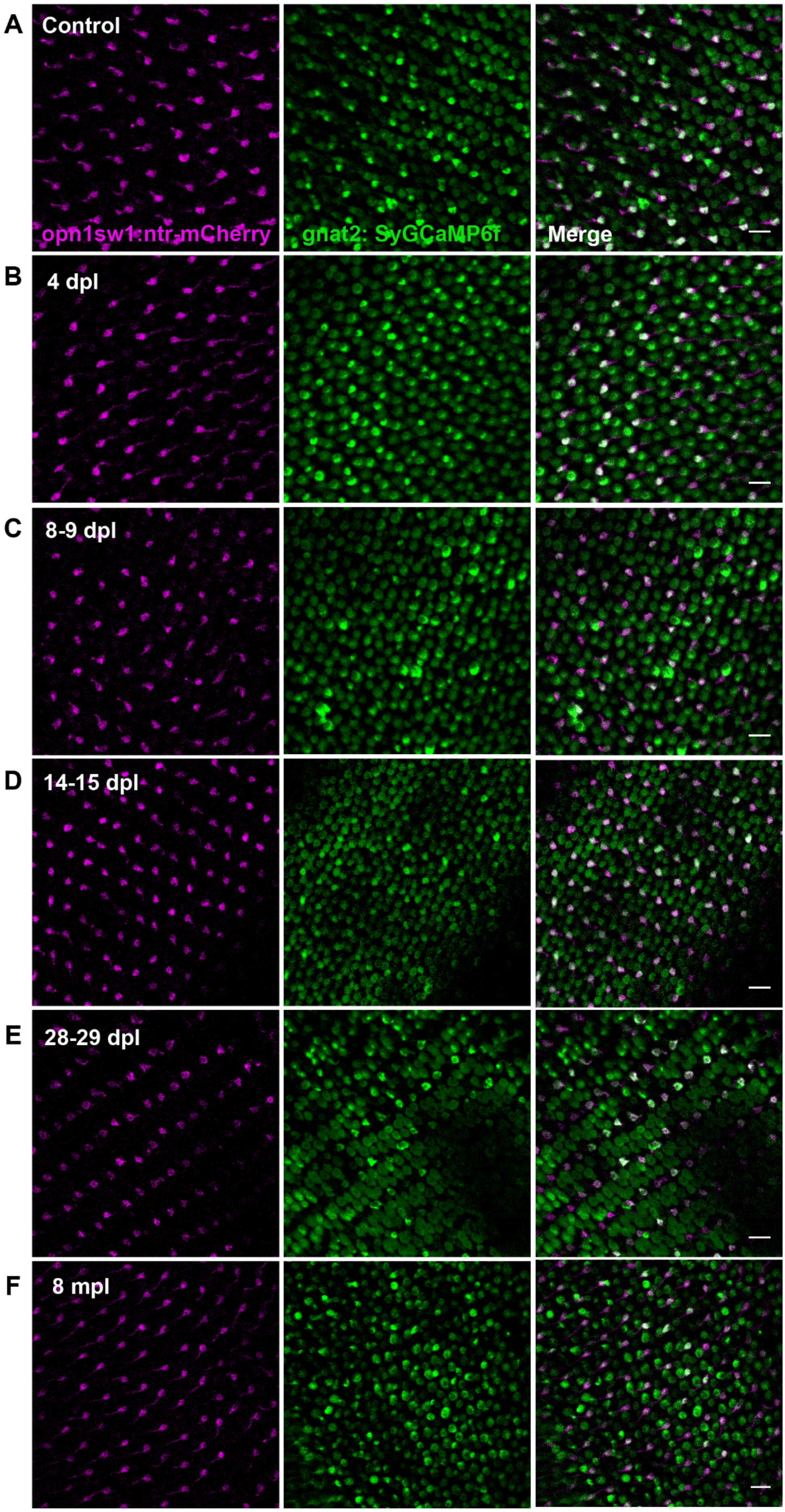
A
Control
opn1sw1:ntr-mCherry
gnat2: SyGCaMP6f
Merge
B
4 dpl
C
8-9 dpl
D
14-15 dpl
E
28-29 dpl
F
8 mpl

**Figure S4: Cone photoreceptors in the ventral retina remain morphologically unlesioned after diffuse light lesion: exemplary morphological scans shown** | Cones in the ventral retina in the double transgenic *Tg(gnat2:SyGCaMP6f); Tg(opn1sw1:nfsb-mCherry)* reporter after diffuse light lesion remain unlesioned. **(A)** Control**, (B)** 4 dpl, **(C)** 8-9 dpl, **(D)** 14-15 dpl, **(E)** 28-29 dpl, **(F)** 8 mpl. Similar to controls, 4 dpl ventral retina is intact with UV, blue, green and red cones, characterized by the intact cone row mosaic arrangement at all examined time points 10-11 dpl (not shown), 14-15 dpl (shown), 28-29 dpl (shown), 3 mpl (not shown) and 8 mpl (shown). Green: SyGCaMP6f, magenta: UV cone mCherry. Scale bar = 10 micrometre.

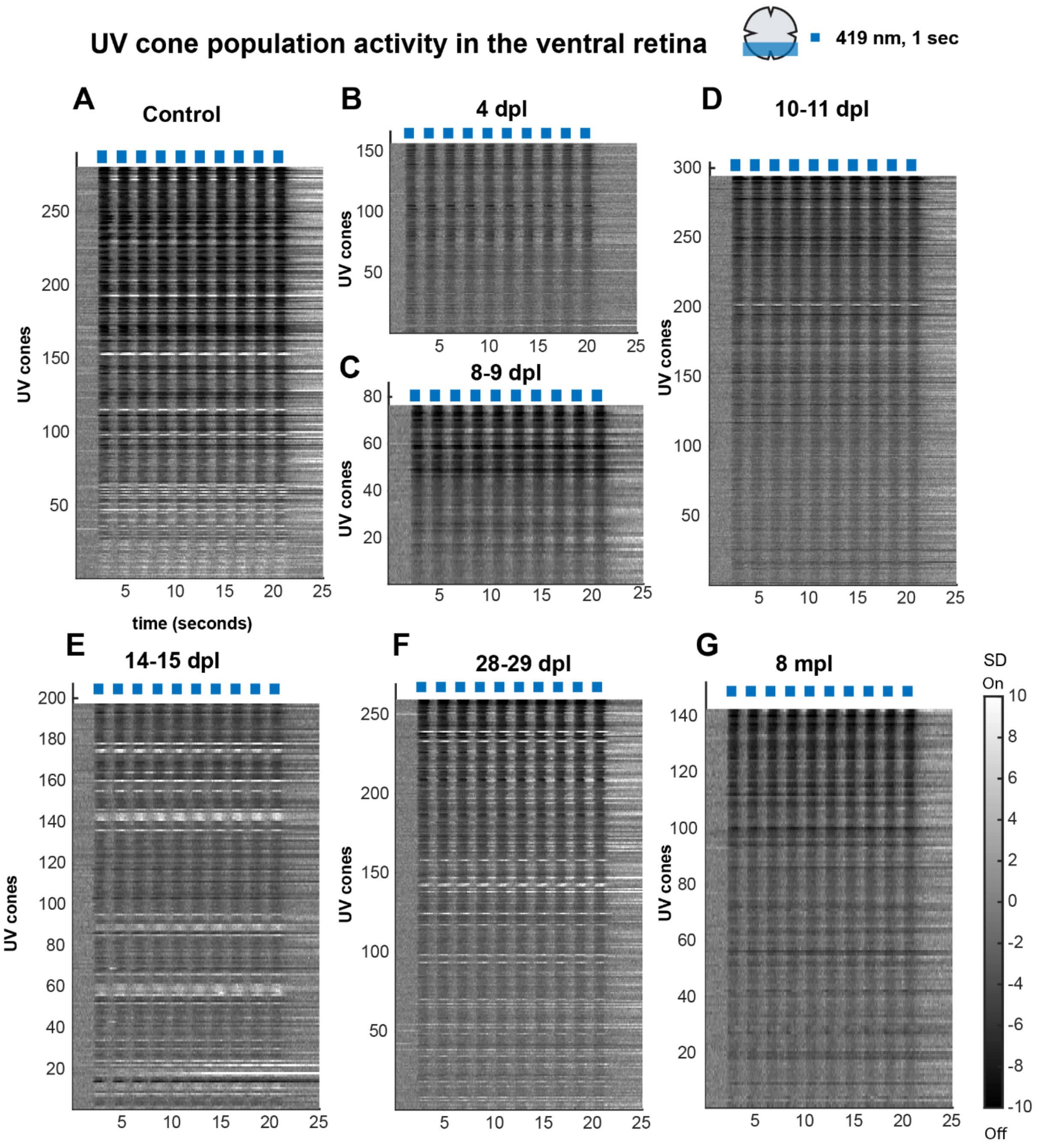

**Figure S5: | UV cone Off responses in the ventral retina is not affected by light lesion at 4 dpl and is an excellent internal control during regeneration | A-G:** Heatmaps representing UV cone responses to blue (419 nm) light at the population level in the ventral unlesioned zone in Z scores (SD) in control and regenerating retinas. (**A**) Control, **(B)** 4 dpl, **(C)** 8-9 dpl, (**D**) 10-11 dpl, (**E**) 14-15 dpl, **(F)** 28-29 dpl, (**G)** 8 mpl. Sample size (n) Control- 280, 4 dpl- 156, 8-9 dpl- 76, 10-11 dpl- 294, 14-15 dpl-197, 28-29 dpl- 259, 8 mpl-142 (n = UV cones); from N= 4, 3, 3, 5, 5, 5, and 3 retinae at respective time points. UV cone traces indicate UV cones which passed cutoff QI >= 0.4 sorted for QI (most reliable responses on top). Only the UV cone response traces where all 10 stimulus triggers were detected during pre-processing are included. Blue bars are stimulus presentations with blue (419 nm) light for 1 second. Grayscale bars are in Z scores (SD). Darker shades indicate a drop in calcium level relative to baseline, denoting a cone's intrinsic light response (Off), while lighter shades indicate a rise in calcium level, and reveal sign-inverted inputs from the outer retinal network (On).

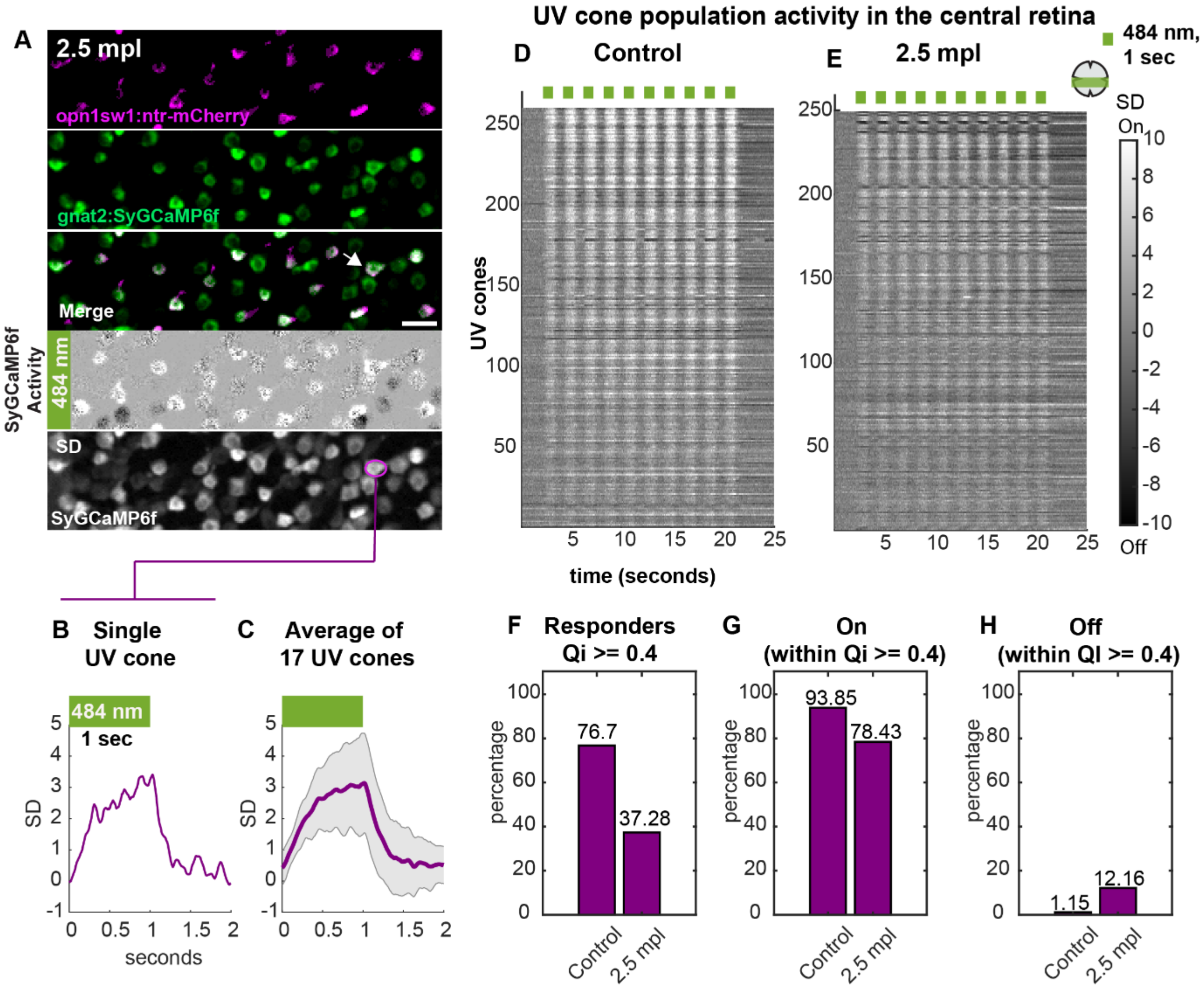

**Figure S6**: **Functional recovery of On response in regenerating UV cones in the central retina after light lesion | (A)** Example functional scan fields from the 2.5 mpl regenerated central retina of the double transgenic *Tg(gnat2:SyGCaMP6f); Tg(opn1sw1:nfsb-mCherry)*, stimulated with green (484 nm) LED, Green- SyGCaMP6f, magenta- UV cone mCherry, Scale bar = 10 micrometre**.** Cone activity and standard deviation (SD) projection of SyGCaMP6f from A shown; with an example UV cone (ROI) marked in magenta. Activity image: Dark terminals represent Off responses, whereas lighter terminals represent On responses; see also **Movie 11**. **(B)** Single UV cone light On response from A. **(C)** Averaged light responses from all n = 17 UV cones from A, represented as mean and standard deviation (shaded error bar). Y- axis: standard deviation (SD) and X- axis: time (seconds). **(D, E)** Heatmaps representing UV cone responses to green (484 nm) light at the population level in Z scores (SD) in control and regenerating retinas. **(D)** Control, **(E)** 2.5 mpl, Sample size (n) Control- 260, 2.5 mpl- 249, (n = UV cones); from N= 5, and 6 retinae, at respective time points. UV cone traces indicate UV cones which passed cutoff QI >= 0.4 sorted for QI (most reliable responses on top). Only the UV cone response traces where all 10 stimulus triggers were detected during pre-processing are shown. Grayscale bars are in Z scores (SD). Darker shades indicate a drop in calcium level relative to baseline, denoting a cone's intrinsic light response (Off), while lighter shades indicate a rise in calcium level, and reveal sign-inverted inputs from the outer retinal network (On). **(F)** Percentage of UV cones satisfying QI >= 0.4 cutoff out of total UV cones; called UV cone responders; that reliably responded to stimulus. **(G)** Percentage of UV cone On responses within UV cone responder population from D. **(H)** Percentage of UV cone Off responses within UV cone responder population from D. Sample size: Total UV cones: Control- 339, 2.5 mpl- 684; Responding UV cones: Control-260, 2.5 mpl- 255; On responses: Control-244, 2.5 mpl- 200; Off responses: Control- 3, 2.5 mpl- 31. Green bar: green (484 nm) stimulus for 1 sec. See also Fig. 4.

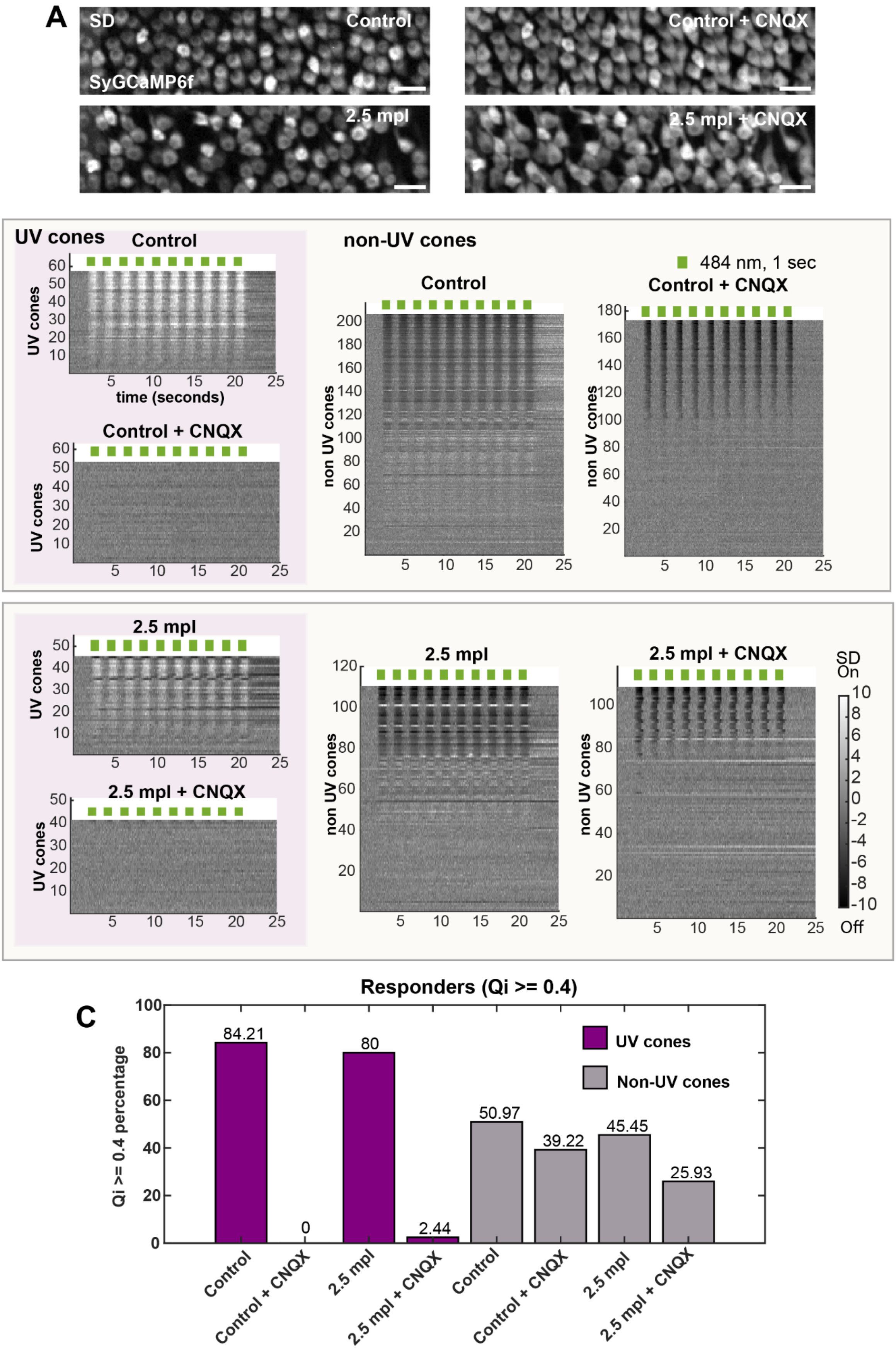
A
SD
Control
SyGCaMP6f
Control + CNQX
2.5 mpl
2.5 mpl + CNQX
B
UV cones
Control
UV cones
time (seconds)
Control + CNQX
non-UV cones
484 nm, 1 sec
Control
Control + CNQX
non UV cones
B'
2.5 mpl
2.5 mpl + CNQX
SD
On
Off
C
Responders (Qi >= 0.4)
Qi >= 0.4 percentage
84.21
0
80
2.44
50.97
39.22
45.45
25.93
UV cones
Non-UV cones
Control
Control + CNQX
2.5 mpl
2.5 mpl + CNQX

**Figure S7: CNQX treatment in unique scan fields from the same retina in the control and 2.5 mpl regenerated retina successfully eliminates all On responses and preserves Off responses | (A)** Standard deviation (SD) projection of SyGCaMP6f from representative scans shown in Fig. 4D and 4F. SD projection is used for placing cone ROIs. Grey: SyGCaMP6f signal. Scale bar = 10 micrometre. **(B)** Heatmap of response traces of UV cones and non-UV cones (in Z scores) from 4 unique scan fields (N= 4 retinae) fields in the control group to green light flashes (green bars on top) before and after HC block, as indicated. Sample size (n): Control- 57, control + HC block- 53 (n = UV cones); Sample size (n): Control- 206, control + HC block- 173, (n = non-UV cones) **(B')**- Heatmap of response traces of UV cones and non UV cones (in Z scores) from 3 unique scan fields (N= 3 retinae) in 2.5 mpl group to green light flashes (green bars on top) before and after HC block, as indicated. Sample size (n): 2.5 mpl- 45, 2.5 mpl + HC block- 41, (n = UV cones); Sample size (n): 2.5 mpl- 110, 2.5 mpl + HC block- 108, (n = non-UV cones). Grayscale bars are in Z scores (SD). Darker shades indicate calcium drop relative to baseline, denoting a cone's intrinsic light response (Off), while lighter shades indicate a rise in calcium and reveal sign-inverted inputs from the outer retinal network (On). In all heatmaps, cone response traces are sorted for increasing QI (most reliable responses on top) and all responses passed cutoff QI >= 0, where 10 stimulus triggers were detected are shown. **(C)** Percentage of UV cone responders before and after CNQX treatment in the control and 2.5 mpl groups. Sample size: Total UV cones: Control- 57, Control + CNQX- 53; Responding UV cones: Control- 48, Control + CNQX- 0; Total UV cones: 2.5 mpl- 45, 2.5 mpl + CNQX- 41; Responding UV cones: 2.5 mpl- 36, 2.5 mpl + CNQX- 1; Total non-UV cones: Control- 206, Control + CNQX- 204; Responding non-UV cones: Control- 105, Control + CNQX- 80; Total non-UV cones: 2.5 mpl-110, 2.5 mpl + CNQX- 108; Responding non-UV cones: 2.5 mpl- 50, 2.5 mpl + CNQX- 28. See also Fig. 4.

Abraham et al

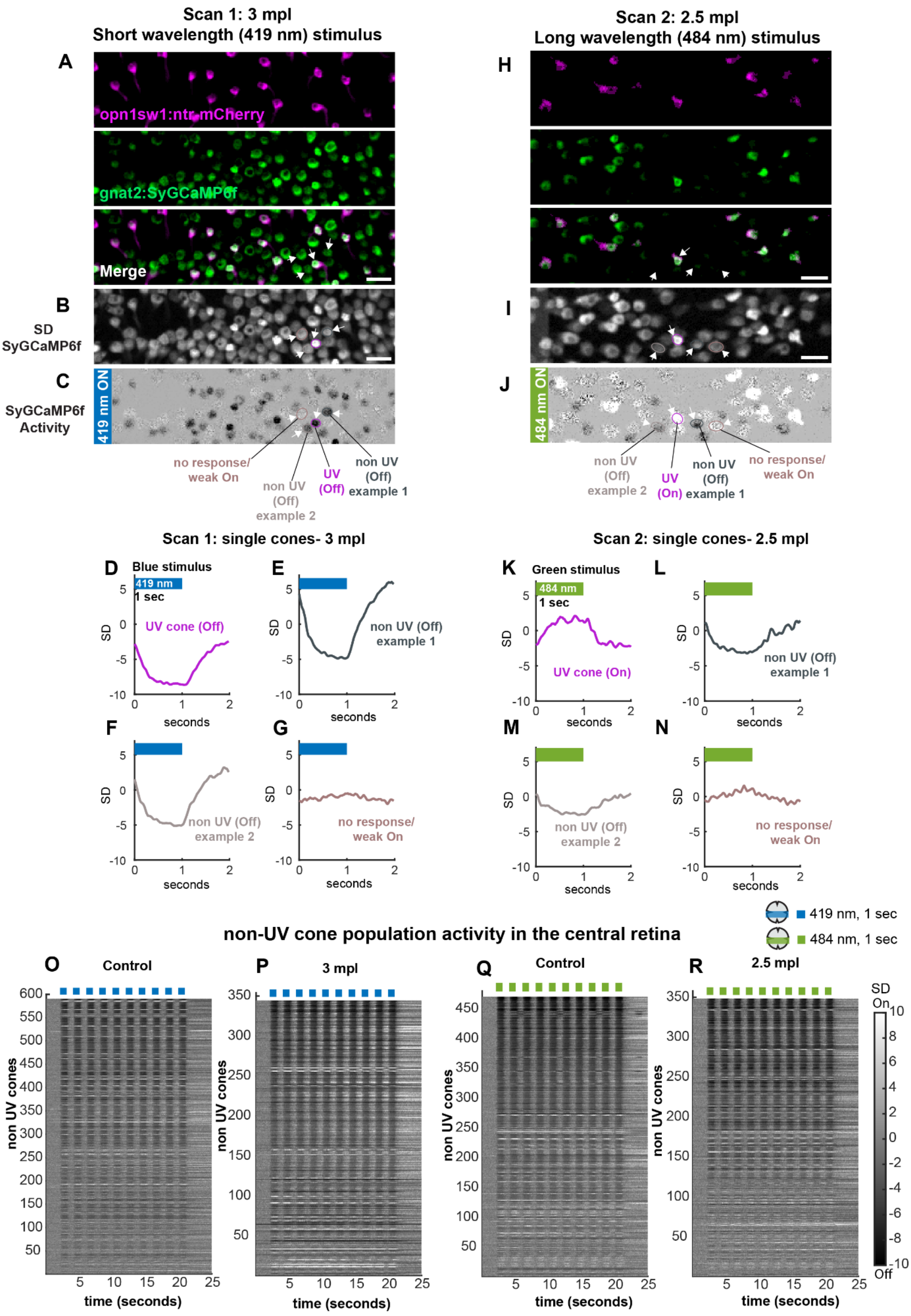

**Figure S8: Non-UV cone responses to short wavelength (blue, 419 nm) and long wavelength (green, 484 nm) light stimulation are recovered in the regenerating retina | (A)** Example functional scan fields from the central retina of double-transgenic *Tg(gnat2:SyGCaMP6f); Tg(opn1sw1:nfsb-mCherry)* retinae at 3 mpl, stimulated with blue light. **(B)** Standard deviation (SD) projection of SyGCaMP6f from A with quartet of ROIs, one UV cone and three non-UV cone ROI marked (also in A and C). **(C)** Activity image of A, showing cone SyGCaMP6f responses to blue light. Dark terminals represent Off responses, whereas lighter terminals represent On responses. **(D-G)** Example cone responses from ROIs in B (also in A and C): UV cone Off response, non UV cone Off response example 1, non UV cone Off response example 2, and non UV cone (non-responsive/ weakly On), respectively. **(H)** Example functional scan fields from the central retina of double-transgenic *Tg(gnat2:SyGCaMP6f); Tg(opn1sw1:nfsb-mCherry)* retinae at 2.5 mpl, stimulated with green light. **(I)** Standard deviation (SD) projection of SyGCaMP6f from H with quartet of ROIs, one UV cone and three non-UV cone ROI marked (also in H and J). **(J)** Activity image of H, showing cone SyGCaMP6f responses to green light. Dark terminals represent Off responses, whereas lighter terminals represent On responses. **(K-N)** Example cone responses from ROIs in I (also in H and J): UV cone On response, non UV cone Off response example 1, non UV cone Off response example 2, and non UV cone (non-responsive/ weakly On), respectively. A and H are separate scans from different retinas. Green- SyGCaMP6f, magenta- UV cone mCherry; Scale bar = 10 micrometre. **(O)** Control, **(P)** 3 mpl, **(Q)** Control, **(R)** 2.5 mpl. In some control scans we also see an On response artifact from blue cones to blue light; this was reasoned as a consequence of calcium responses from an imaing plane closer to the soma. Sample size(n): blue stimulus: Control- 590, 3 mpl- 344, (n = non UV cones); from N= 5, and 3 retinae; green stimulus: Control- 470, 2.5 mpl- 348, (n = non UV cones); from N= 5, and 6 retinae. Non-UV cone traces indicate non-UV cones which passed cutoff QI >= 0.4 sorted for QI (most reliable responses on top). Only the non UV cone response traces where all 10 stimulus triggers were detected during pre-processing are included. Blue and green bars are stimulus presentations with blue (419 nm) and green (484 nm) light for 1 second respectively. Grayscale bars are in Z scores (SD). Darker shades indicate a drop in calcium level relative to baseline, denoting a cone's intrinsic light response (Off), while lighter shades indicate a rise in calcium level, and reveal sign-inverted inputs from the outer retinal network (On). See also Fig. 1, 2 and 4.

### SUMMARY STATISTICS

| TABLE 1: AMPLITUDE (SD) | | | | | | |
|---|---|---|---|---|---|---|
| **Group name** | **Sample size (UV cones)** | **Mean (SD)** | **95% CI (SD)** | **Pairwise comparisons** | **pTukey (p value)** | **Cohen's d (d value)** |
| Control | 576 | -7.11 | [-7.3636, -6.8648] | Control with 8-9 dpl | 0.000000E+00 | -1.74 |
| 8-9 dpl | 28 | -1.91 | [-2.2342, -1.5816] | Control with 10-11 dpl | 0.000000E+00 | -1.52 |
| 10-11 dpl | 89 | -2.71 | [-3.0568, -2.3586] | Control with 14-15 dpl | 0.000000E+00 | -1.38 |
| 14-15 dpl | 182 | -3.25 | [-3.5174, -2.9896] | Control with 28-29 dpl | 0.000000E+00 | -1.20 |
| 28-29 dpl | 643 | -4.02 | [-4.1806, -3.8595] | Control with 3 mpl | 0.000000E+00 | -0.68 |
| 3 mpl | 469 | -5.26 | [-5.4631, -5.0548 | Control with 8 mpl | 2.622376E-21 | -4.95E-01 |
| 8 mpl | 571 | -5.80 | [-5.9794, -5.6173] | | | |
| **ANOVA** | | | | | | |
| **'Source'** | **'SS'** | **'df'** | **'MS'** | **'F'** | **'Prob>F'** | |
| 'Groups' | 4757.022105 | 6 | 792.8370174 | 143.6060936 | 3.32E-157 | |
| 'Error' | 14083.85384 | 2551 | 5.520914871 | [] | [] | |
| 'Total' | 18840.87594 | 2557 | [] | [] | [] | |

**Table 1: Summary statistics of UV cone Off response amplitude during regeneration: related to Figure 3B |** Time course of development of UV cone Off response amplitude during regeneration time course. One-way analysis of variance or ANOVA followed by Tukey Kramer post-hoc analysis for unequal sample sizes was used whenever multiple pairwise comparisons of independent control and experimental groups were done. Summary of groups, sample size, mean, 95% confidence interval (95% CI) of mean, pairwise comparisons, p values corresponding to significance of multiple comparisons of control group to 6 regenerated time points, and effect size of the difference for each comparison in terms of Cohen's d. ANOVA results prior to Tukey Kramer post-hoc analysis for unequal sample sizes summarized. 'SS'- sum of squares, 'df' - degrees of freedom, 'MS'- mean squared error, 'F'- *F* statistic. The 'Prob'- is the *p*-value or probability that the test statistic can take a value greater than the value of the computed test statistic. Cohen's d (d value) can be read as Very small- 0.01, Small- 0.20, Medium- 0.50, Large- 0.80, Very large-1.20, Huge- 2.0. Sign indicates direction of the effect.

| TABLE 2: TAU DECAY (seconds) | | | | | | |
|---|---|---|---|---|---|---|
| **Group name** | **Sample size (UV cones)** | **Mean (sec)** | **95% CI (sec)** | **Pairwise comparisons** | **pTukey (p value)** | **Cohen's d (d value)** |
| Control | 576 | 0.24 | [0.2347, 0.2413] | Control with 8-9 dpl | 2.413950E-04 | -0.76 |
| 8-9 dpl | 28 | 0.27 | [0.2377, 0.3037] | Control with 10-11 dpl | 1.938520E-08 | 0.64 |
| 10-11 dpl | 89 | 0.21 | [0.2003, 0.2218] | Control with 14-15 dpl | 9.461147E-01 | 0.08 |
| 14-15 dpl | 182 | 0.23 | [0.2261, 0.2430] | Control with 28-29 dpl | 0.000000E+00 | 0.70 |
| 28-29 dpl | 643 | 0.21 | [0.2078, 0.2136] | Control with 3 mpl | 1.821909E-10 | 0.46 |
| 3 mpl | 469 | 0.22 | [0.2191, 0.2242] | Control with 8 mpl | 1.801557E-10 | 0.43 |
| 8 mpl | 571 | 0.22 | [0.2198, 0.2251] | | | |
| **ANOVA** | | | | | | |
| **'Source'** | **'SS'** | **'df'** | **'MS'** | **'F'** | **'Prob>F'** | |
| 'Groups' | 0.326418179 | 6 | 0.05440303 | 36.4435769 | 1.44E-42 | |
| 'Error' | 3.808136873 | 2551 | 0.001492802 | [] | [] | |
| 'Total' | 4.134555052 | 2557 | [] | [] | [] | |

**Table 2: Summary statistics of UV cone Off response Tau decay during regeneration: related to Figure 3C |** Time course of development of UV cone Off response Tau decay during regeneration time course. One-way analysis of variance or ANOVA followed by Tukey Kramer post-hoc analysis for unequal sample sizes was used whenever multiple pairwise comparisons of independent control and experimental groups were done. Summary of groups, sample size, mean, 95% confidence interval (95% CI) of mean, pairwise comparisons, p values corresponding to significance of multiple comparisons of control group to 6 regenerated time points, and effect size of the difference for each comparison in terms of Cohen's d. ANOVA results prior to Tukey Kramer post-hoc analysis for unequal sample sizes summarized. 'SS'- sum of squares, 'df' - degrees of freedom, 'MS'- mean squared error, 'F'- *F* statistic. The 'Prob'- is the *p*-value or probability that the test statistic can take a value greater than the value of the computed test statistic. Cohen's d (d value) can be read as Very small- 0.01, Small- 0.20, Medium- 0.50, Large- 0.80, Very large-1.20, Huge- 2.0. Sign indicates direction of the effect.

| TABLE 3: TAU RECOVERY (seconds) | | | | | | |
|---|---|---|---|---|---|---|
| **Group name** | **Sample size (UV cones)** | **Mean (sec)** | **95% CI (sec)** | **Pairwise comparisons** | **pTukey (p value)** | **Cohen's d (d value)** |
| Control | 576 | 0.32 | [0.3090, 0.3293] | Control with 8-9 dpl | 4.692240E-20 | -2.43 |
| 8-9 dpl | 28 | 0.73 | [0.5133, 0.9546] | Control with 10-11 dpl | 0.000000E+00 | -1.90 |
| 10-11 dpl | 89 | 0.75 | [0.6395, 0.8686] | Control with 14-15 dpl | 0.000000E+00 | -0.93 |
| 14-15 dpl | 182 | 0.53 | [0.4673, 0.5838] | Control with 28-29 dpl | 4.708746E-12 | -0.47 |
| 28-29 dpl | 643 | 0.42 | [0.3964, 0.4360] | Control with 3 mpl | 1.453363E-02 | -0.34 |
| 3 mpl | 469 | 0.37 | [0.3528, 0.3819] | Control with 8 mpl | 9.634973E-01 | -0.10 |
| 8 mpl | 571 | 0.33 | [0.3209, 0.3435] | | | |
| **ANOVA** | | | | | | |
| **'Source'** | **'SS'** | **'df'** | **'MS'** | **'F'** | **'Prob>F'** | |
| 'Groups' | 23.92810984 | 6 | 3.988018307 | 74.19481218 | 1.44E-85 | |
| 'Error' | 137.1178712 | 2551 | 0.053750636 | [] | [] | |
| 'Total' | 161.0459811 | 2557 | [] | [] | [] | |

**Table 3: Summary statistics of UV cone Off response Tau recovery during regeneration: related to Figure 3D |** Time course of development of UV cone Off response Tau recovery during regeneration time course. One-way analysis of variance or ANOVA followed by Tukey Kramer post-hoc analysis for unequal sample sizes was used whenever multiple pairwise comparisons of independent control and experimental groups were done. Summary of groups, sample size, mean, 95% confidence interval (95% CI) of mean, pairwise comparisons, p values corresponding to significance of multiple comparisons of control group to 6 regenerated time points, and effect size of the difference for each comparison in terms of Cohen's d. ANOVA results prior to Tukey Kramer post-hoc analysis for unequal sample sizes summarized. 'SS'- sum of squares, 'df' - degrees of freedom, 'MS'- mean squared error, 'F'- *F* statistic. The 'Prob'- is the *p*-value or probability that the test statistic can take a value greater than the value of the computed test statistic. Cohen's d (d value) can be read as Very small- 0.01, Small- 0.20, Medium- 0.50, Large- 0.80, Very large-1.20, Huge- 2.0. Sign indicates direction of the effect.

| TABLE 4: QUALITY INDEX - Qi (a.u.) | | | | | | |
|---|---|---|---|---|---|---|
| **Group name** | **Sample size (UV cones)** | **Mean (a.u.)** | **95% CI (a.u.)** | **Pairwise comparisons** | **pTukey (p value)** | **Cohen's d (d value)** |
| Control | 854 | 0.78 | [0.7600, 0.7975] | Control with 8-9 dpl | 0.000000E+00 | 1.69 |
| 8-9 dpl | 250 | 0.23 | [0.2040, 0.2501] | Control with 10-11 dpl | 0.000000E+00 | 1.33 |
| 10-11 dpl | 457 | 0.33 | [0.3106, 0.3540] | Control with 14-15 dpl | 0.000000E+00 | 0.62 |
| 14-15 dpl | 744 | 0.40 | [0.3824, 0.4237] | Control with 28-29 dpl | 0.000000E+00 | -0.22 |
| 28-29 dpl | 1281 | 0.61 | [0.5921, 0.6223] | Control with 3 mpl | 2.698662E-03 | 0.10 |
| 3 mpl | 600 | 0.83 | [0.8174, 0.8435] | Control with 8 mpl | 2.667984E-01 | |
| 8 mpl | 886 | 0.75 | [0.7348, 0.7676] | | | |
| **ANOVA** | | | | | | |
| **'Source'** | **'SS'** | **'df'** | **'MS'** | **'F'** | **'Prob>F'** | |
| 'Groups' | 174.8500898 | 6 | 29.14168163 | 448.3657671 | 0 | |
| 'Error' | 329.2013536 | 5065 | 0.064995331 | [] | [] | |
| 'Total' | 504.0514434 | 5071 | [] | [] | [] | |

**Table 4: Summary statistics of UV cone response quality during regeneration: related to Figure 3E |** Time course of development of Quality index (QI) of UV cones during regeneration time course. One-way analysis of variance or ANOVA followed by Tukey Kramer post-hoc analysis for unequal sample sizes was used whenever multiple pairwise comparisons of independent control and experimental groups were done. Summary of groups, sample size, mean, 95% confidence interval (95% CI) of mean, pairwise comparisons, p values corresponding to significance of multiple comparisons of control group to 6 regenerated time points, and effect size of the difference for each comparison in terms of Cohen's d. ANOVA results prior to Tukey Kramer post-hoc analysis for unequal sample sizes summarized. 'SS'- sum of squares, 'df' - degrees of freedom, 'MS'- mean squared error, 'F'- *F* statistic. The 'Prob'- is the *p*-value or probability that the test statistic can take a value greater than the value of the computed test statistic. Cohen's d (d value) can be read as Very small- 0.01, Small- 0.20, Medium- 0.50, Large- 0.80, Very large-1.20, Huge- 2.0. Sign indicates direction of the effect.

| **TABLE 5: AMPLITUDE (SD)** | | | | | | |
|---|---|---|---|---|---|---|
| **Group name** | **Sample size** | **Mean (SD)** | **95% CI (SD)** | **Pairwise comparisons** | **p value** | **Cohen's d (d value)** |
| Control | 182 | 5.35 | [4.9206, 5.7720] | Control with 2.5 mpl | 1.095918E-13 | 0.88 |
| 2.5 mpl | 136 | 3.21 | [2.9385, 3.4717] | | | |

**Table 5: Summary statistics of UV cone On response amplitude during regeneration: related to Figure 4B |** Time course of development of UV cone On response amplitude in regenerating UV On cones. Two sample t-test for unequal sample sizes and independent groups was used to compare control and experimental group. Summary of groups, sample sizes, mean, 95% confidence interval (95% CI), pairwise comparison, p values, and effect size of the difference in terms of Cohen's d value are indicated. Cohen's d (d value) can be read as Very small- 0.01, Small- 0.20, Medium- 0.50, Large- 0.80, Very large-1.20, Huge- 2.0. Sign indicates direction of the effect.

| **TABLE 6: QUALITY INDEX – Qi (a.u.)** | | | | | | |
|---|---|---|---|---|---|---|
| **Group name** | **Sample size** | **Mean (a.u.)** | **95% CI (a.u.)** | **Pairwise comparisons** | **p value** | **Cohen's d (d value)** |
| Control | 339 | 0.61 | [0.5787, 0.6316] | Control with 2.5 mpl | 2.002196E-48 | 1.02 |
| 2.5 mpl | 684 | 0.36 | [0.3407, 0.3763] | | | |

**Table 6: Summary statistics of UV cone response quality during regeneration: related to Figure 4C |** Time course of development Quality Index (QI) in regenerating UV cones. Two sample t-test for unequal sample sizes and independent groups was used to compare control and experimental group. Summary of groups, sample sizes, mean, 95% confidence interval (95% CI), pairwise comparison, p values, and effect size of the difference in terms of Cohen's d value are indicated. Cohen's d (d value) can be read as Very small- 0.01, Small- 0.20, Medium- 0.50, Large- 0.80, Very large-1.20, Huge- 2.0. Sign indicates direction of the effect.
