## Supplementary material for "Restoration of Cone Circuit Functionality in the Regenerating Adult Zebrafish Retina": Movies 2 Photon

**Supplemental Movies:**

**SyGCaMP6f light response and corresponding activity (grey)**

Activity (see methods) shows synaptic cone SyGCaMP6f calcium responses to blue and green stimulus respectively. Dark terminals represent Off responses, whereas lighter terminals represent On responses.

Stimulus: Flash ON for 1 sec, and Flash OFF for 1 sec.

Total duration: 2 sec.

**Movie 1:** Fig. 1D, Control_1 *(blue, 419 nm)*

**Movie 2:** Fig. 1F, Control_2 *(green, 484 nm)*

**Movie 3:** Fig. 2D, Control_3 *(blue, 419 nm)*

**Movie 4:** Fig. 2D, 4 dpl *(blue, 419 nm)*

**Movie 5:** Fig. 2D, 8-9 dpl *(blue, 419 nm)*

**Movie 6:** Fig. 2D, 10-11 dpl *(blue, 419 nm)*

**Movie 7:** Fig. 2D, 14-15 dpl *(blue, 419 nm)*

**Movie 8:** Fig. 2D, 28-29 dpl *(blue, 419 nm)*

**Movie 9:** Fig. 2D, 3 mpl *(blue, 419 nm)*

**Movie 10:** Fig. 2D. 8 mpl *(blue, 419 nm)*

**Movie 11:** Fig. S6A, 2.5 mpl_1 *(green, 484 nm)*

**Movie 12:** Fig. 4D, Control_4 +CNQX *(green, 484 nm)*

**Movie 13:** Fig. 4F, 2.5 mpl_2 + CNQX *(green, 484 nm)*

**Movie 1: Control_1 (blue, 419 nm)	Movie 2: Control _2 (green, 484 nm)**

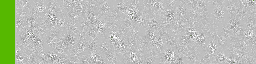

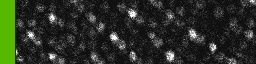

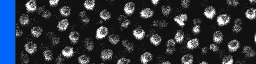

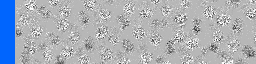

**Movie 3: Control_3 (blue, 419 nm)	Movie 4: 4 dpl (blue, 419 nm)**

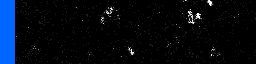

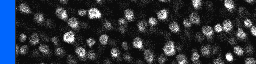

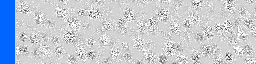

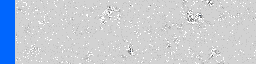

**Movie 5: 8-9 dpl (blue, 419 nm)	Movie 6: 10-11 dpl (blue, 419 nm)**

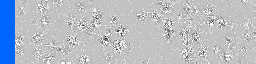

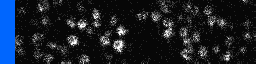

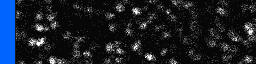

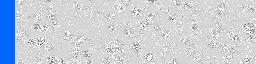

**Movie 7: 14-15 dpl (blue, 419 nm)	Movie 8: 28-29 dpl (blue, 419 nm)**

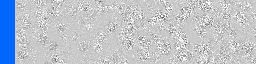

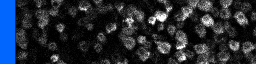

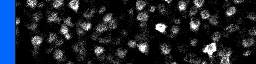

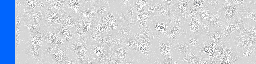

**Movie 9: 3 mpl (blue, 419 nm)	Movie 10: 8 mpl (blue, 419 nm)**

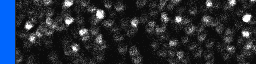

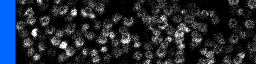

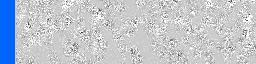

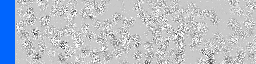

**Movie 11: 2.5mpl_1 (green, 484 nm)**

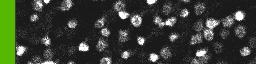

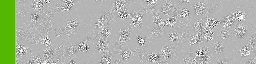

**Movie 12: Control_4	+ HC block (green, 484 nm)**

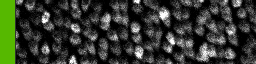

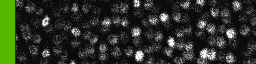

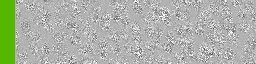

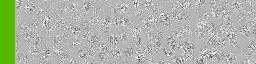

**Movie 13: 2.5 mpl_2	+ HC block (green, 484 nm)**

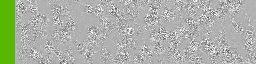

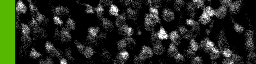

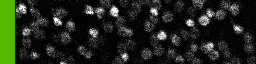

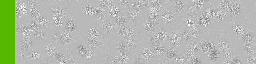
